# A novel therapeutic antibody screening method using bacterial high-content imaging reveals functional antibody binding phenotypes of *Escherichia coli* ST131

**DOI:** 10.1101/2020.05.22.110148

**Authors:** Mailis Maes, Zoe A. Dyson, Sarah E. Smith, David A. Goulding, Catherine Ludden, Stephen Baker, Paul Kellam, Stephen T. Reece, Gordon Dougan, Josefin Bartholdson Scott

## Abstract

The increase of antimicrobial resistance (AMR), and lack of new classes of licensed antimicrobials, have made alternative treatment options for AMR pathogens increasingly attractive. Recent studies have demonstrated anti-bacterial efficacy of a humanised monoclonal antibody (mAb) targeting the O25b O-antigen of *Escherichia coli* ST131. To evaluate the phenotypic effects of antibody binding to diverse clinical *E. coli* ST131 O25b bacterial isolates in high-throughput, we designed a novel mAb screening method using high-content imaging (HCI) and image-based morphological profiling to screen a mAb targeting the O25b O-antigen. Screening the antibody against a panel of 86 clinical *E. coli* ST131 O25:H4 isolates revealed 4 binding phenotypes: no binding (18.60%), weak binding (4.65%), strong binding (69.77%) and strong agglutinating binding (6.98%). Impaired antibody binding could be explained by the presence of insertion sequences or mutations in O-antigen or lipopolysaccharide core biosynthesis genes, affecting the amount, structure or chain length of the O-antigen. The agglutinating binding phenotype was linked with lower O-antigen density, enhanced antibody-mediated phagocytosis and increased serum susceptibly. This study highlights the need to screen candidate mAbs against large panels of clinically relevant isolates, and that HCI can be used to evaluate mAb binding affinity and potential functional efficacy against AMR bacteria.

## Introduction

Antimicrobial resistance (AMR) is one of the greatest current challenges in global health^1^. Increasing AMR and a lack of antimicrobials in the pharmaceutical pipeline makes alternative therapeutic approaches increasingly important. Passive antibody transfer has historically been used to treat bacterial infections, such as diphtheria, tetanus, and pneumococcal pneumonia^2,3^, making antibodies a potential therapeutic approach. Many biologics, including monoclonal antibodies (mAbs) are being used increasingly in oncology, autoimmune diseases, and the prevention of some viral infections^4,5^. However, mAbs have been found to have limited utility against bacteria^6^. This lack of effectiveness is, in part, because bacterial species are generally antigenically diverse and conserved immunogenic surface components can be masked by large structures such as capsules. Identifying tractable therapeutic antibody targets that are generic or specific for particular pathogenic or AMR bacteria would be a valuable addition to our current arsenal of therapeutic options.

Numerous globally dispersed clades of pathogenic bacteria associated with broad AMR phenotypes have emerged in recent decades^7–9^. One example is the ST131 O25b:H4 clonal group of *Escherichia coli* that is characterised by the acquisition of extended-spectrum beta-lactamases (ESBLs) and fluoroquinolone resistance^10^. Notably, *E. coli* ST131 O25b:H4 have a specific O-antigen, which is potentially an attractive target for mAbs. The humanised monoclonal antibody, 3E9-11, specifically targeting this O25b O-antigen has recently demonstrated promising efficacy^11^. This antibody, which is in preclinical development, exhibits multiple modes of antibacterial activity and exhibited protection in mice^11,12^. To be of clinical utility it is important to demonstrate that these anti-bacterial activities function against a diverse collection of *E. coli* ST131 O25b associated with disease in healthcare settings.

High-content imaging (HCI) is a powerful phenotypic screening approach that combines high-throughput automated microscopy with comprehensive image analysis to quantify multiple morphological and functional cellular features. This type of image-based morphological profiling can be used for high-throughput screening of drugs, simultaneously evaluating potency as well as mode-of-action^13,14^. HCI has been predominantly applied to mammalian cells and tissue where the examined variables include cell and organelle shape, signal transduction, gene expression and metabolism. In addition to studying mammalian cells, HCI has also been used to study intracellular pathogens such as *Mycobacterium tuberculosis*^15,16^, and more recently individual bacteria under drug exposure^17^. HCI has now reached a point where populations of bacteria can be phenotyped at single cell resolution in high-throughput to enable the simultaneous screening of multiple isolates of diverse bacterial clades.

Here we designed a novel mAb screening method using HCI and image-based morphological profiling to measure the antimicrobial potential of a variant of 3E9-11, which targets the O-antigen of *E. coli* ST131 O25b:H4. We profiled 86 *E. coli* ST131 O25b:H4 clinical isolates in imaging assays at a level of resolution that identified individual bacteria in 96 well plates. Our analysis revealed distinct mAb binding phenotypes within the *E. coli* ST131 O25b:H4 population that were directly associated with mAb function.

## Results

### Screening antibodies against bacteria using high-content imaging

To evaluate HCI as a method for screening candidate mAbs against large panels of clinical isolates, whilst simultaneously determining the diagnostic and functional potentials of the antibody, we synthesised KM467, an IgG1 antibody based on the VH and VL sequences of 3E9-11, which specifically targets the O25b O-antigen of *E. coli* ST131. KM467 was tested for the ability to bind lipopolysaccharide (LPS) isolated from the *E. coli* ST131 O25b reference strain NCTC13441 using ELISA (Supplementary Fig. S1), and direct binding to whole bacteria was tested using the Perkin Elmer Opera Phenix high-content confocal microscope (Fig. 1a). KM467 recognized the target in both assays: exhibiting a clear titration curve in the ELISA and a strong staining pattern of the bacterial surface by confocal imaging.

**Figure 1:**
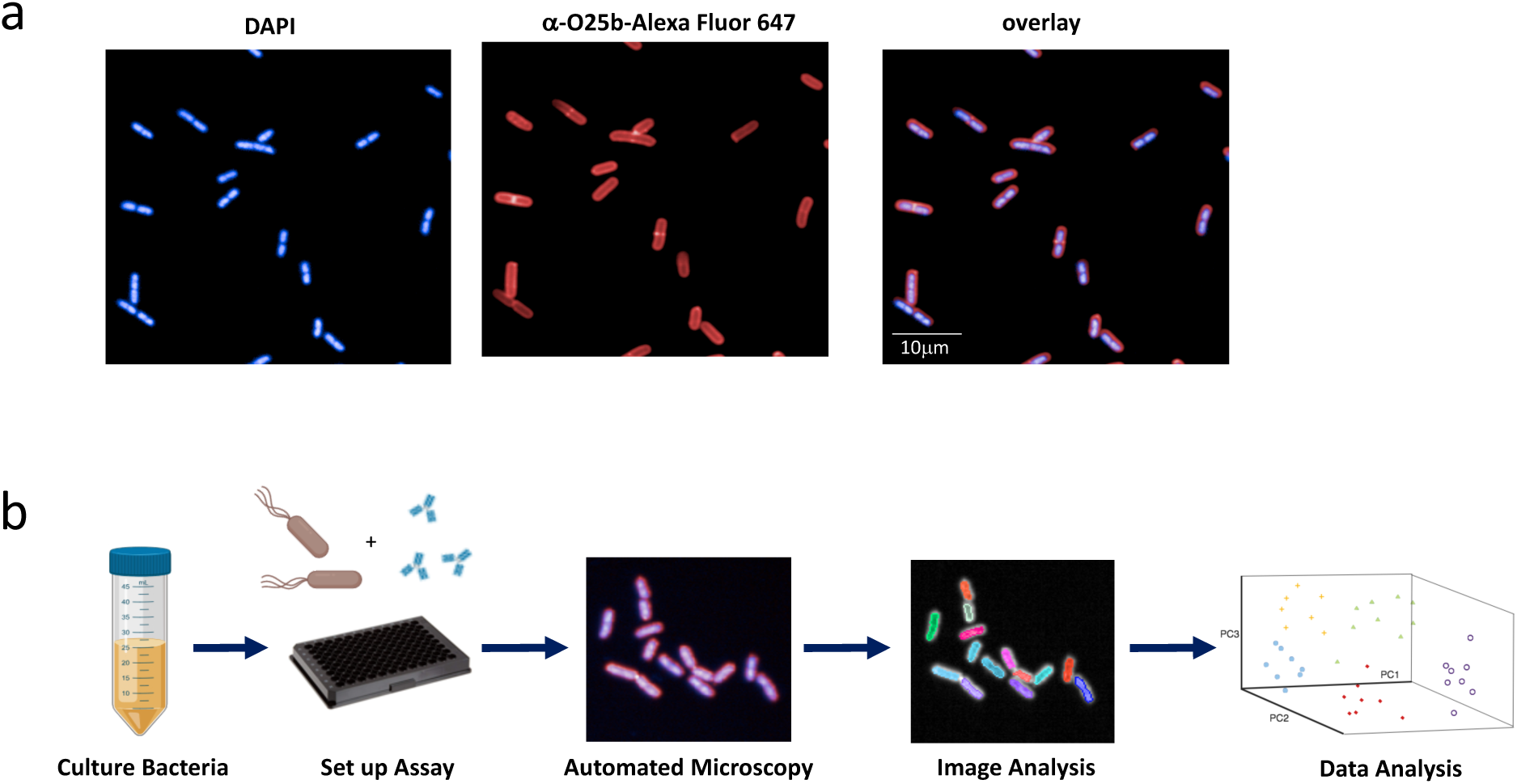
High-content imaging to screen mAbs against bacteria. *E. coli* ST131 NCTC13441 were stained with DAPI (nucleic acids) and KM467 followed by an Alexa Flour 647-conjugated secondary antibody (**a**). Bacterial high-content imaging workflow (**b**): bacterial overnight cultures were diluted and added to microtiter plates and left to adhere for 2 hours at 37°C. Plates were fixed, and incubated with KM467 for 1 hour, followed by an Alexa Flour 647-conjugated secondary antibody and DAPI (nucleic acid stain) for 30 min. The plates were imaged on the Opera Phenix using a 63x water immersion objective, and the images were analysed using the Harmony software. Data was exported into R for further analysis.

A bacterial antibody high-content screening workflow was developed for higher throughput screening as outlined in Fig. 1b. For bacterial imaging, overnight liquid cultures of *E. coli* NCTC13441 were diluted and added to ultra-thin, flat-bottom 96 well plates and left to adhere for 2 hours at 37°C. Bacteria were fixed with paraformaldehyde and incubated with KM467. Finally, bacteria were stained with DAPI and an Alexa Fluor 647-conjugated anti-human IgG secondary antibody *in situ* (Fig. 1a). The plates were imaged on an Opera Phenix using a 63x water immersion objective and the images were analysed using the Harmony software. A full 96 well plate with 16 fields and 3 Z-stacks per well took 1 hour to image, and the images displayed clearly defined individual bacteria (Fig. 1a). Images were segmented using the DAPI channel and the Harmony software ‘Find Spots’ building block, then size filters were applied to distinguish individual bacteria. A small outer border around each bacterium was included to identify Alexa Fluor 647 bound around the bacterial cell surface (Fig. 2a), and the Alexa Fluor 647 intensity was calculated for each segmented bacterium (Fig. 2b). An average of 2×10^3^ bacteria per well were analysed using this approach.

**Figure 2:**
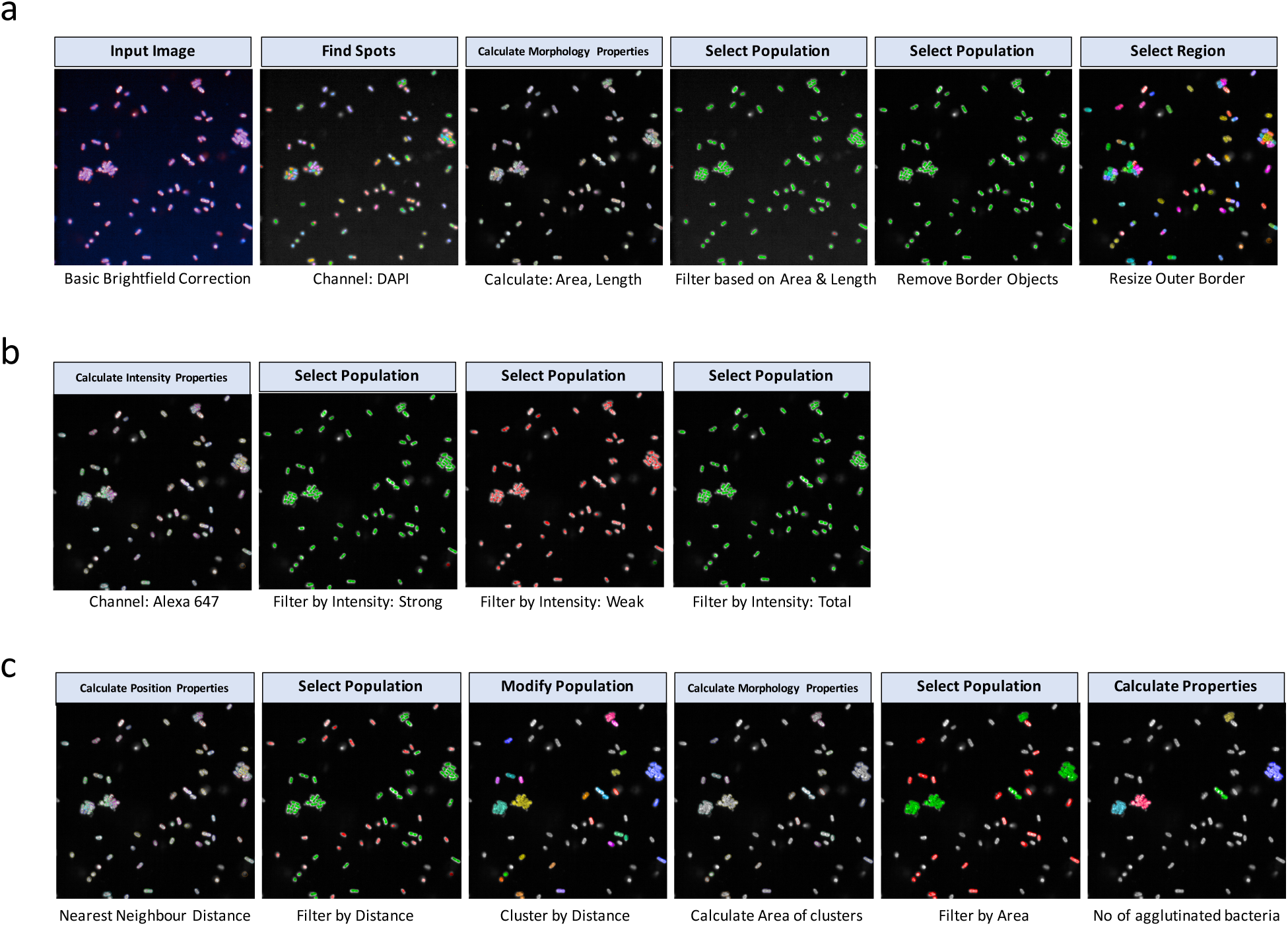
Antibody binding image analysis. Images were segmented in Harmony using the Find Spots building block and size filters were applied to distinguish individual bacteria and the outer border of each bacteria was resized to maximize antibody stain coverage (**a**). Intensity properties were calculated based on Alexa Fluor 647 intensity (antibody binding) and three sub-populations were defined based on strong, weak and total Alexa Fluor 647 intensity (**b**). Agglutination was defined based on bacterial position properties: nearest neighbour distance was used to filter and define bacterial clusters (**c**). Clusters were further filtered based on area to define true agglutination, finally, the number of agglutinating bacteria was calculated.

A KM467 titration curve was performed using the NCTC13441 O25b reference strain, as well as an *E. coli* ST131 isolate with O16 O-antigen (VRES0673) (Supplementary Fig. S2). No KM467 binding was observed to the O16 isolate, confirming specificity to the O25b O-antigen. The titration curve generated using image analysis (Supplementary Fig. S2b) correlated with the standard curve from the ELISA (Supplementary Fig. S1), and demonstrated that the optimal concentration of KM467 was 1μg/ml. Bacterial imaging and antibody binding were found to be reproducible with no significant background from the secondary antibody in the negative control or when using the *E. coli* O16 isolate (Supplementary Fig. S2).

### Screening KM467 against clinical *E. coli* ST131 isolates revealed four distinct binding phenotypes

KM467 was screened against a panel of 86 *E. coli* ST131 O25:H4 clinical isolates (Supplementary Table S1) in 96 well format and images were captured on the Opera Phenix and segmented using the Perkin Elmer Harmony software as described above (Fig. 2a-c, Supplementary Table S2). This screen revealed four distinct binding phenotypes (Fig. 3a) that were classified as: no binding (NB), weak binding (WB), strong binding (SB) and strong agglutinating binding (SAB). To confirm that the weak binding was not an artefact, titrated antibody was tested against representative isolates for each of the phenotypic classes (Supplementary Fig. S3). Although the percentage binding observed for the weak binders was much lower than that observed for the SB and SAB isolates, weak binding was genuine, as evidenced by increased binding observed with higher mAb concentrations and no binding observed with the secondary antibody only. The NB isolates displayed no observable KM467 binding at any concentration, even at high exposure (Supplementary Fig. S3). Based on the Alexa Fluor 647 intensities observed for the different phenotypes, intensity thresholds could be determined to differentiate between weak and strong antibody binding (Fig. 2b).

**Figure 3:**
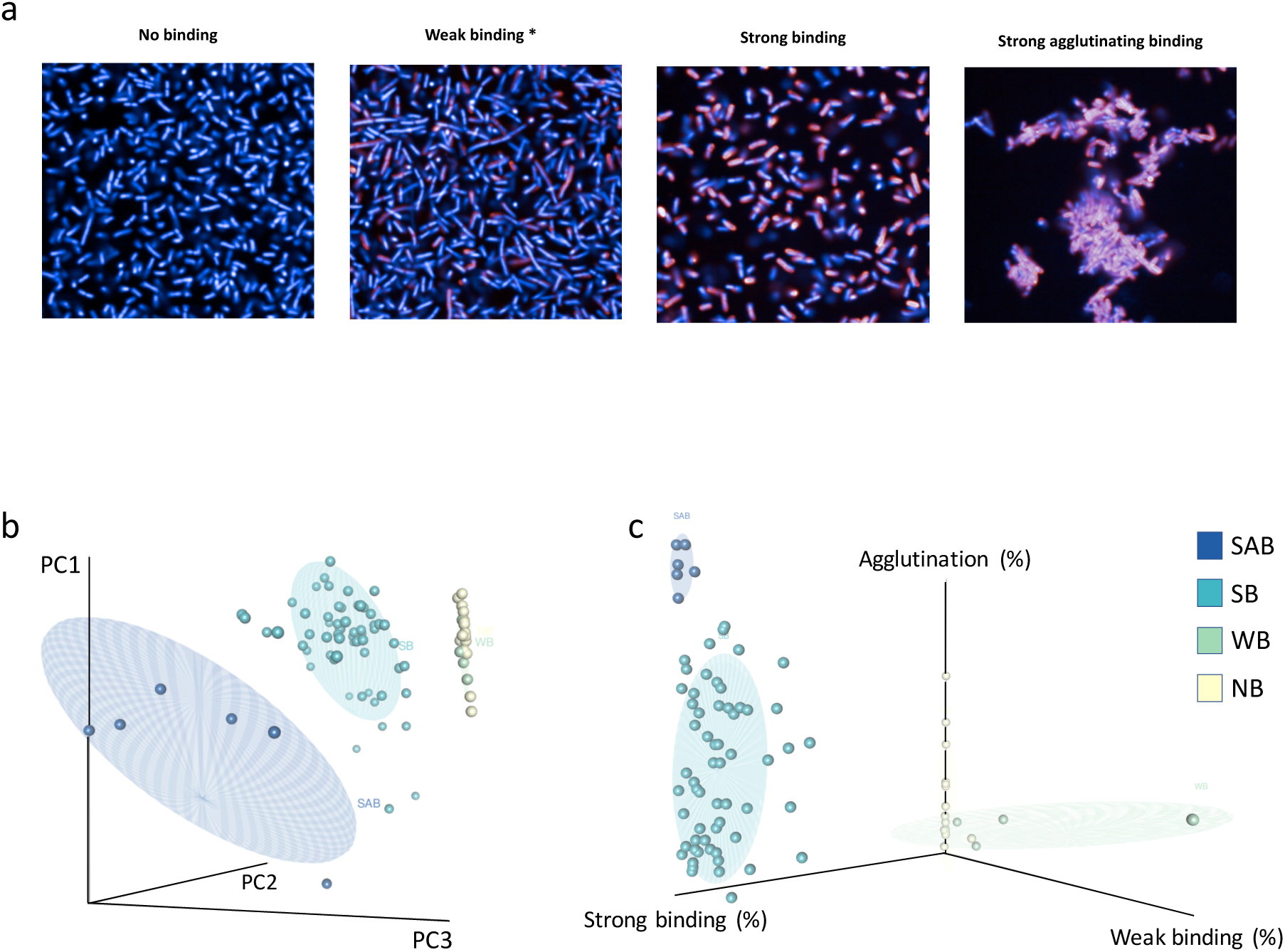
Screening KM467 against clinical *E. coli* ST131 isolates revealed four distinct binding phenotypes. Four antibody binding phenotypes were observed (**a**): no binding, weak binding, strong binding and strong agglutinating binding. *Over-exposed compared to other images. Principal Component Analysis distinguishing the different binding phenotypes (**b**) and a graph separating the phenotypes based on percentage weak binding, strong binding and agglutination (**c**) were generated in R.

For efficient image segmentation, the bacterial density in the well needed to be optimal so that bacteria were sufficiently separated. This was particularly challenging for the agglutinating bacteria that were tightly clustered together. Conversely, this challenge formed the basis for measuring agglutination, allowing for automated identification of the SAB phenotype. To perform this analysis, the segmented bacterial positional properties were calculated: i.e. measuring distance to the “nearest neighbour” and by defining agglutination by cluster area and the quantification of bacteria within clusters (Fig. 2c, Supplementary Table S2). This approach, in combination with intensity measurements, convincingly separated the four phenotypes by principal component analysis (PCA) (Fig. 3b). The WB isolates clustered closely with the NB, but the SB isolates separated markedly from the WB and NB, and SAB isolates could be clearly distinguished from the non-agglutinating phenotypes. In addition, the four phenotypes could be effectively separated based on the proportions showing weak binding, strong binding and agglutination (Fig. 3c). Out of the 86 isolates 16 exhibited no KM467 binding (18.60%), 4 WB (4.65%), 60 SB (69.77%) and 6 SAB (6.98%) (Supplementary Table S1, Supplementary Table S3). One isolate (ECO0237) agglutinated both in the presence and absence of antibody but grouped with the SB isolates in all analyses, therefore this isolate was classified as SB since the antibody did not induce the agglutination. These results demonstrate that HCI can be used for high-throughput antibody screening and found that a single antibody can induce distinct *E. coli* ST131 isolate-specific phenotypic effects due to binding.

### Differences in O-antigen correlate with binding phenotypes

To investigate the different binding phenotypes, LPS was extracted from 35 representative isolates including the NCTC13441 reference. Silver staining revealed that a lack of binding or WB appeared to be associated with a lack of O-antigen, reduced O-antigen production or differing O-antigen lengths compared to the conserved O-antigen of the binders (Fig. 4a, and Supplementary Fig. S4). We could observe no obvious differences between the SB and the SAB organisms.

**Figure 4:**
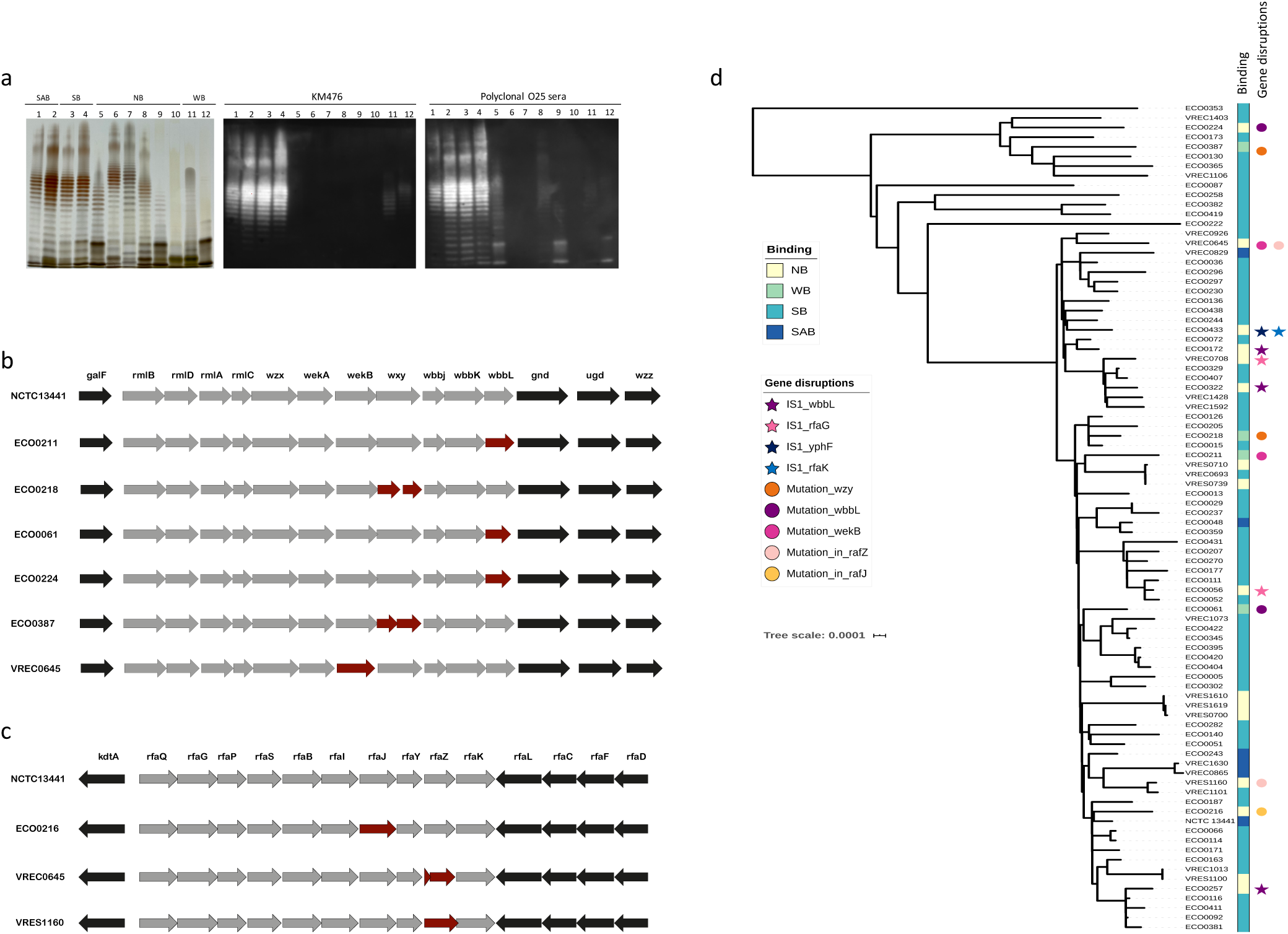
O-antigen integrity influences binding phenotypes. Isolated LPS was analysed by gel electrophoresis followed by silver staining (left) and immunoblotting using KM467 (middle) or O25 polyclonal antisera (right) (**a**). Lanes 1-12: NCTC13441, VREC0829, VREC1073, VREC1403, VREC0708, VRES1100, VRES1160, VRES1610, ECO0056, ECO0172, ECO0061, ECO0218. Binding phenotypes are indicated above the gel (lane 1-2 are SAB, lane 3-4 are SB, lane 5-10 are NB and lane 11-12 are WB). The O25b O-antigen genes (**b**) and LPS core genes (**c**) of NCTC13441 and a selection of isolates that did not bind KM467 were aligned using Prokka annotated fasta files and plotted using EasyFig2.2.2. Genes affected by mutations are highlighted in red. A core genome maximum likelihood phylogenetic tree of the *E. coli* ST131 O25b isolates was aligned with the antibody binding phenotypes and presence of mutations (circles) or IS*1* (stars) (**d**).

A selection of LPS representing each binding phenotype was further analysed by immunoblotting using KM467 and polyclonal anti-O25 typing sera (Fig. 4a). Immunoblotting using KM467 correlated with the HCI data, with strong bands only observed for those isolates with SB or SAB binding phenotypes. Isolates classified as WB by HCI also had weak staining by immunoblotting with KM467; however, their banding pattern was distinct from that of the SB and SAB isolates. The polyclonal anti-O25 sera proficiently stained the O-antigen of the SB and SAB isolates, but also showed weaker binding to some of the isolates that showed weak or no KM467 binding. Notably, the NB isolates with higher molecular weight O-antigen exhibited no binding even in the presence of the polyclonal sera. These results show that the inability of KM467 to bind some isolates is associated with differences in or lack of O-antigen.

### Correlating genome content with phenotype

To address the observed differences in O-antigen in terms of genetics, we obtained whole genome sequence data for each isolate in our study and analysed the read data using SRST2^18^ in combination with an *E. coli in silico* serotyping scheme^19^. This analysis confirmed that all 86 isolates were genotypically serotype O25b (Supplementary Table S1). Raw read data were assembled with Unicycler^20^ and alignments of the O-antigen gene cluster and flanking genes (*galF, rmlB, rmlD, rmlA, rmlC, wzx, wekA, wekB, wzy, wbbj, wbbK, wbbL, gnd, ugd, wzz*) from each isolate were extracted manually to identify any genetic variation in this region (Fig. 4b, Table 1, Supplementary Fig. S5).

**Table 1:**
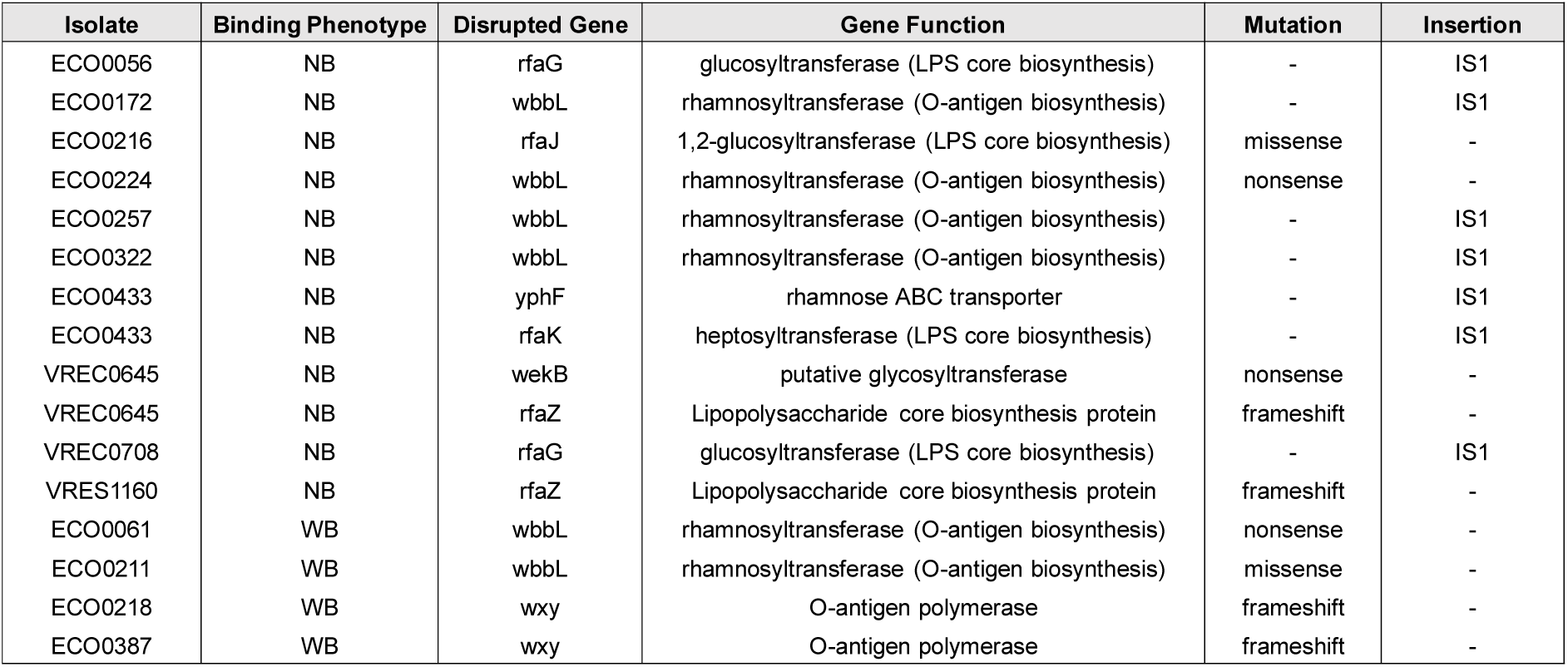
O-antigen and LPS core genes disrupted by mutations or *IS*1.

Two of the WB isolates, ECO0218 and ECO0387, harboured frameshift mutations resulting in a split of *wzy* into two open reading frames. The *wzy* gene encodes an O-antigen polymerase that regulates O-antigen chain length, and disruption of this gene may explain reduced KM467 binding. This observation is consistent with the lack of O-antigen observed by silver staining (Fig. 4a lane 12, Supplementary Fig. S4 lanes 27 and 30). A third WB isolate (ECO0061) and one NB isolates (ECO0224) had a truncated *wbbL* gene which encodes a rhamnosyltransferase. In addition, the fourth WB isolate (ECO0211) had a missense mutation in the same gene resulting in a change from glycine to glutamic acid at codon 60 (G60E). These mutations likely impact both O-antigen integrity and the antibody binding affinity as each O25b O-antigen repeat contains both rhamnose and *O*-acetyl-rhamnose^21^. A second NB isolate (VREC0645) had a premature stop codon in the putative glycosyltransferase *wekB*. The remaining 10 isolates that displayed no binding to the antibody appeared to have intact O-antigen operons, despite having atypical O-antigen repeat patterns.

To further investigate the lack of KM467 binding to these NB isolates, hybrid short and long-read assemblies from one SAB, one SB and three NB isolates were aligned using Mauve^22^, and inspected manually to identify potential inversions and indels in the O-antigen, LPS core and capsule biosynthesis regions. Insertion sequence IS*Ec10* was detected upstream of *kpsF* and IS*1* downstream of *kpsT* in the capsule biosynthesis regions of the NCTC13441 reference isolate, and subsequently we screened for the presence of IS*Ec10* and IS*1* using ISmapper^23^ in all genomes. The presence of IS*Ec10* upstream of *kpsF* did not appear to correlate with binding phenotype, nor was this type of IS found in any other relevant genomic regions (Supplementary Table S4). In addition, the presence of IS*1* downstream of *kpsT* was only observed in the NCTC13441 reference isolate that had an SAB antibody binding phenotype. However, IS*1* was present in three NB isolates in *wbbL* (ECO0172, ECO0257 and ECO0322), and in two NB isolates (ECO0056 and VREC0708) in *rfaG* encoding a glucosylranferase involved in LPS core biosynthesis (Table 1). Mutations in *rfaG* have previously been associated with a rough LPS phenotype^24^. However, both these isolates displayed some shorter O-antigen repeating units, indicating that RfaG was not completely inactive (Fig. 4a lanes 5 and 9, Supplementary Fig. S4 lanes 11 and 16). ECO0433 had an IS*1* in rhamnose ABC transporter gene *yphF*, which could potentially affect the rhamnose composition of the O-antigen, and also an IS*1* in *rfaK*, which encodes a heptosyltransferase involved in core biosynthesis, which may explain the lack of O-antigen observed by LPS gel electrophoresis (Supplementary Fig. S4, lane 35).

Since IS*1* was found in genes affecting the LPS core, the core biosynthesis gene cluster (*kdtA, rfaQ, rfaG, rfaP, rfaS, rfaB, rfaI, rfaJ, rfaY, rfaZ, rfaK, rfaL, rfaC, rfaF, rfaD*) was extracted manually from assemblies and analysed by multiple sequence alignment (Fig. 4c, Table 1, Supplementary Fig. S5), and mutations were identified in two additional genes. NB isolate ECO0216 had a mutation resulting in a premature stop codon in *rfaJ* which encodes a 1,2-glucosyltransferase, and VREC0645 and VRES1160 (both NB) had frameshift mutations in *rfsZ* which is involved in 3-deoxy-D-manno-oct-2-ulosonic acid (Kdo) III attachment in the inner LPS core^25^. Although the core bands appear to be affected in both isolates, interestingly, VRES1160 displayed longer O-antigen repeats that were not detected by the polyclonal anti-O25 sera (Fig. 4a lane 7). In total, the identified mutations and the presence of IS*1* accounted for 10 of 16 NB and all four of the WB isolates.

A maximum-likelihood phylogenetic tree was inferred from core genome SNP alignments of the study isolates. Phenotypic binding data and the presence of mutations in the O-antigen or core biosynthesis region and presence of IS*1* within these was overlaid as metadata to observe any potential relationship between phenotype and genotype (Fig. 4d). There was no relationship between phylogenetic cluster and KM467 binding phenotype in this data set, and gene disruptions were associated with mutation and recombination events in individual isolates.

### Antibody-induced phagocytosis is more efficient in agglutinating isolates

To identify any potential functional differences between the antibody binding phenotypes, macrophage phagocytosis and opsonophagocytosis assays were carried out in the presence and absence of KM467. Assays were conducted in ultra-thin, flat bottom 96 well plates and imaged on the Opera Phenix using the 40x air objective. Cells were stained with CellMask Orange (a plasma membrane stain), and DAPI (nuclear stain) and bacteria were visualised using KM467 labelled with Alexa Flour 647. As the NB and WB isolates could not be visualised using KM467, serum from mice immunised with *E. coli* outer-membrane vesicles followed by an Alexa Fluor 647 secondary was used. Images were analysed in Harmony using predefined building blocks to segment nuclei and cytoplasm in the eukaryotic cells, to define the number of these cells, and to count individual bacteria within the cells (Fig. 5a, Supplementary Table S5). Initially, the macrophage phagocytosis assays were performed in the absence of complement. Although the bacterial cultures were OD600 adjusted to 1.00, there were significant differences in phagocytosis between isolates without antibody present (Supplementary Fig. S6). When plotting the average increase in phagocytosis in the presence of KM467, phagocytosis of the SB isolates increased by ∼0.5 bacteria/cell while the SAB increased by ∼3 bacteria/cell (Fig. 5b). No significant increase was observed for the WB and the NB isolates as expected. These results indicate that the KM467 agglutinated bacteria are more efficiently phagocytosed by macrophages.

**Figure 5:**
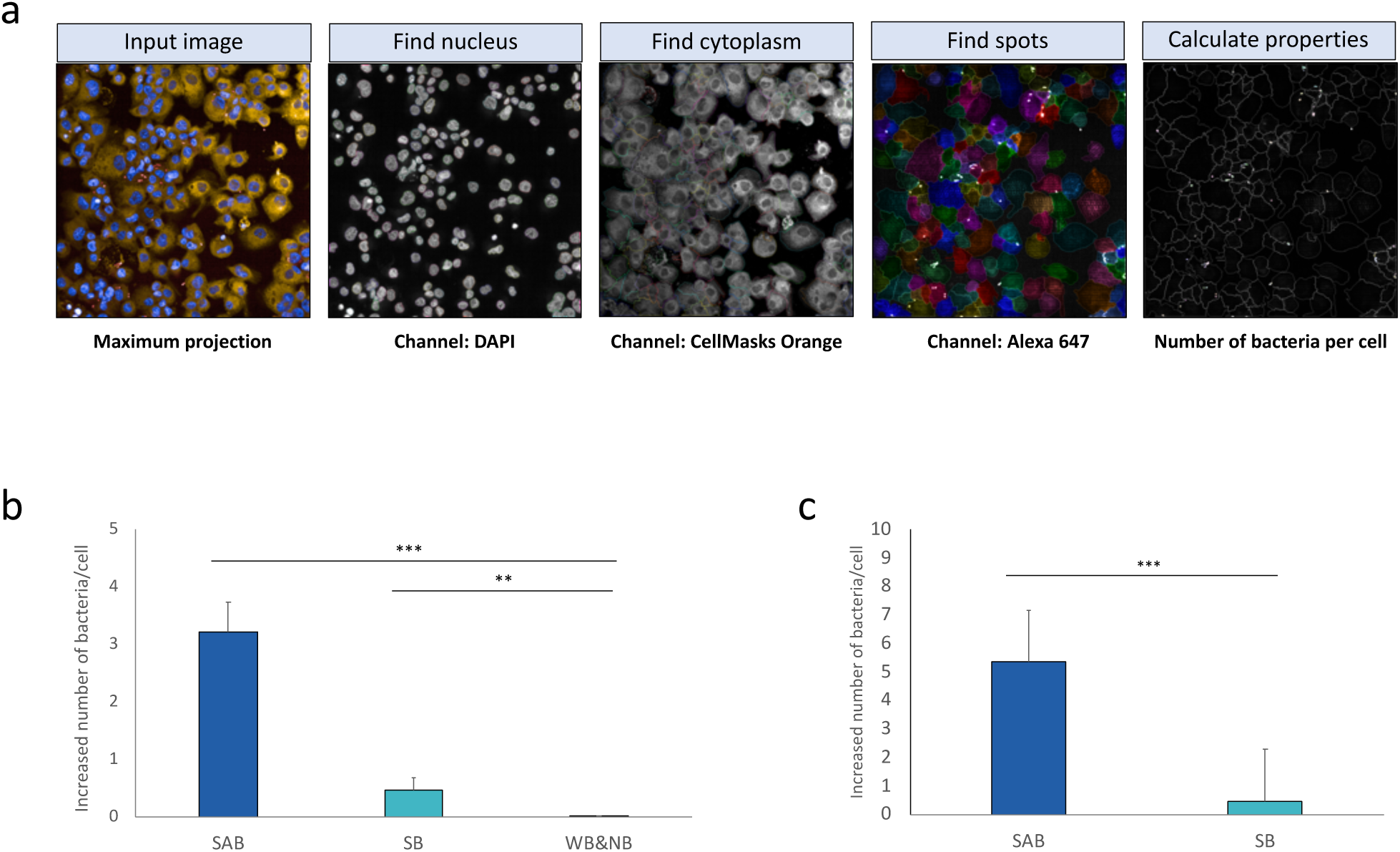
Clearance by phagocytosis and opsonophagocytosis is directly linked to binding phenotype. Macrophage phagocytosis and opsonophagocytosis imaging and analysis (**a**): cells were stained with CellMasks Orange (cytoplasm) and DAPI (nucleic acid) and bacteria were visualised using KM467 labelled with Alexa Flour 647. Images were obtained using a 40x air objective on the Opera Phenix. Images were analysed in Harmony using predefined building blocks to segment nuclei and cytoplasm to define the number of cells, and to count individual bacteria (using Find Spots) within the cells. Macrophage phagocytosis (without complement) (**b**) and macrophage opsonophagocytosis (with complement) (**c**) was plotted as the average increase in bacterial uptake per cell in the presence of antibody. The average of 4 representative isolates per phenotype is shown and error bars represent standard deviation of 3 replicates. Significance was determined by *t*-test (*=<0.05, **= between 0.001 and 0.05, *** <0.001).

The assay was then repeated for the SB and SAB isolates in the presence of baby rabbit complement (BRC) (Fig. 5c). The presence of complement and antibody augmented phagocytosis of the SAB isolates but not of the SB isolates, indicating that the antibody-induced agglutination may be more important than just antibody binding for efficient phagocytosis of *E. coli* ST131.

### Binding phenotypes correlate with differences in serum resistance and antibody-induced complement-dependent cytotoxicity (CDC)

To further evaluate the functional impact of the different antibody binding phenotypes, their serum resistance was evaluated using BRC. *E. coli* belonging to ST131 are the most commonly isolated *E. coli* from blood in bacteraemia patients, and have, not surprisingly, previously been characterised as being serum resistant^26^. Isolates that exhibited a SB phenotype generally grew well in serum (>80% compared to LB) (Fig. 6a). However, the remaining isolates (that show either no, weak and agglutinating binding with the antibody) grew to different extents in serum. The WB and NB isolates showed ∼30% and ∼40% survival respectively, and the SAB isolates showed less than 20% survival compared to bacteria grown in LB (Fig. 6a). Comparable results were observed by using both optical density measurements (OD600) and CFU counting (Supplementary Fig. S7). Serum resistance did not seem to correlate with the source of the isolates nor with ESBL-plasmid carriage (Supplementary Table S1), which has previously been linked to enhanced serum resistance^27^.

**Figure 6:**
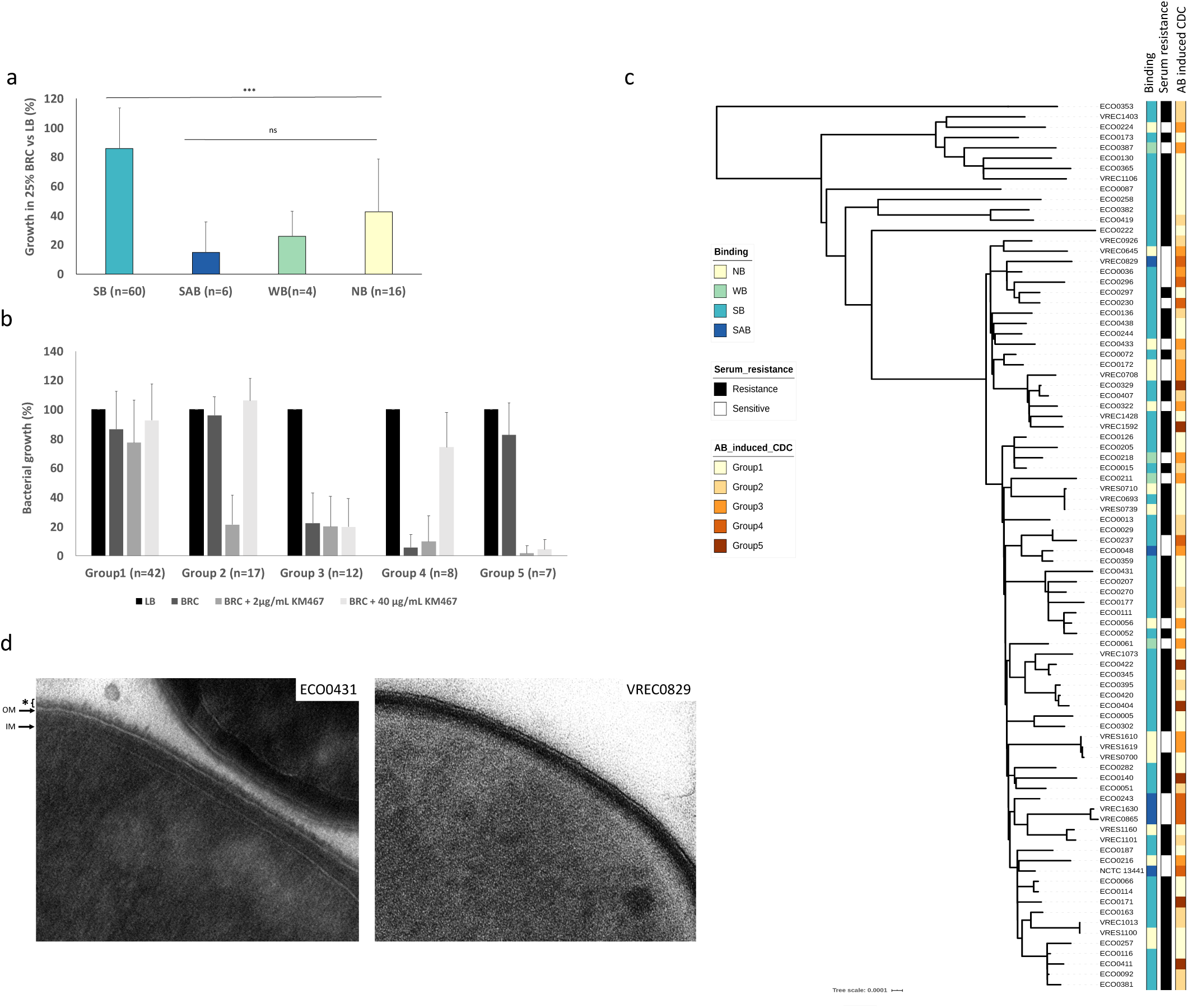
Differences in serum survival and complement-dependent cytotoxicity is linked to O-antigen density and correlate with binding phenotypes. Growth in LB broth in the presence or absence of 25% BRC was measured by OD600 after 4 hours and the average for all isolates of each phenotype was plotted as percentage growth in the presence of serum compared to LB alone (**a**). Error bars represent standard deviation. Significance was determined by *t*-test (*=<0.05, **= between 0.001 and 0.05, *** <0.001). CDC was investigated for all isolates in the presence of 25% BRC alone, 25% BRC plus 2 or 40μg/ml KM467 and plotted as percentage growth compared to LB alone (**b**). Growth patterns were grouped into 5 different CDC groups. Error bars represent standard deviation. A core genome maximum likelihood phylogenetic tree of the *E. coli* ST131 O25b isolates was aligned with the antibody binding phenotypes, serum resistance and antibody-induced CDC (**c**). Transmission electron microscopy images of a SAB (VREC0829) and a SB (ECO0431) isolate (**d**). Arrows indicate outer (OM) and inner membrane (IM), * indicates surface matrix layer.

To determine if KM467 could induce CDC in any of the isolates, survival in the presence of BRC and two different concentrations of KM467 was investigated. Interestingly, the isolates could be classified into five groups based on the CDC results (Fig. 6b). Group 1 (n=42): were resistant to complement even in the presence of antibody. Group2 (n=17): were resistant to complement, but susceptible when 2μg/ml KM467 was present, however, in the presence of increased amount of KM467 (40µg/ml) they were protected from complement cytotoxicity. Group 3 (n=12): were sensitive to complement alone and in both KM467 concentrations. Group 4 (n=8): were susceptible to complement alone and in the presence of 2μg/ml KM467, however, when 40μg/ml KM467 was added these isolates were protected from complement cytotoxicity. Group 5 (n=7): were complement resistant, but susceptible at both KM467 concentrations. These results were aligned to the phylogenetic tree of the isolates with the KM467 binding data (Fig. 6c). The majority of SB isolates were serum resistant, and all SAB and WB isolates were serum sensitive, whereas the NB phenotype did not correlate with serum resistance. The NB isolates that displayed either rough LPS or shorter O-antigen repeats by LPS gel electrophoresis were all serum sensitive. All isolates that have an agglutinating phenotype in the presence of KM467, except ECO0048, fell in CDC group 4, all WB isolates in group 3, and most SB isolates were in groups 1 and 2. In these experiments, the majority of isolates were resistant to complement even in the presence of the KM467, and a significant proportion of the remaining susceptible isolates were protected from complement killing in high antibody concentrations.

The finding that antibody binding phenotypes correlate with different levels of serum resistance might suggest different O-antigen densities. Transmission electron microscopy (TEM) was performed on an SB (group 1, serum and CDC resistant) and an SAB (group 4, serum sensitive) isolate. The TEM images revealed differences in the thickness of surface matrix surrounding their outer membrane, which is likely due to O-antigen density (Fig. 6d). The serum resistant SB isolate ECO0431 displayed a high amount of surface matrix (∼13nm), whereas the serum sensitive SAB isolate VREC0829 only had a low amount (∼2nm). This observation suggests that the surface matrix density is likely important for protection against serum killing in *E. coli* ST131 O25b isolates. Taken together these data suggest that an agglutinating binding phenotype may be linked to a less dense surface matrix layer and that this enhances both antibody-mediated phagocytosis and complement susceptibly.

## Discussion

HCI offers the opportunity to analyse bacterial populations at single cell resolution, which enables the visualisation and analysis of direct antibody binding to their individual bacterial targets at scale. This imaging can be undertaken at high-throughput, in 96-well microtiter plates, and the same analysis can be compared across multiple plates. Consequently, this approach has the potential to enable screening of a large number of candidate mAbs against a reference bacterial isolate, followed by functional screening of the most promising mAbs against large panels of clinically relevant isolates. HCI analysis software can carry out automated morphological image analysis which, for example, measures percentage binding, binding intensity and cell shape (single cell or agglutination).

Here, we have demonstrated that HCI can be used to screen a therapeutic mAb candidate against bacteria in high-resolution and in high-throughput, allowing us to study direct binding of mAbs to their native targets. When screening an O25b O-antigen mAb against a panel of 86 closely related clinical *E. coli* ST131 O25b:H4 isolates, distinct binding phenotypes were identified that correlated with mAb function both in CDC and phagocytosis assays. Despite all isolates being genetically typed as O25b, 16 of the 86 isolates showed no binding and four showed weak binding to the antibody. These binding phenotypes correlated with differences in O-antigen presentation compared to the isolates that showed strong antibody binding. For the majority of isolates (10/16 NB, 4/4 of WB) this could be tentatively explained by the presence of an IS*1* insertion sequence or mutations in genes in O-antigen or LPS core biosynthesis gene loci, which impacted the quantity, structure, or length of the repeating O-antigen units. Thus, it is possible to link genotype to phenotype, directing more intensive experimental investigations.

The majority (66/86) of isolates strongly bound the antibody, with six out of these displaying an agglutinating binding phenotype. Interestingly, the agglutinated isolates were more efficiently phagocytosed by macrophages in comparison to the non-agglutinated isolates and in addition, the agglutinating isolates were markedly more sensitive to complement killing. The association between the agglutinating phenotype with increased phagocytosis is not surprising as larger clusters of bacteria are more efficiently phagocytosed than individual bacteria; this has been observed previously for other bacterial species^28^. Agglutination may be linked with a less dense surface matrix layer (likely O-antigen) compared to the non-agglutinating SB isolates, although more work would be required to confirm this. O-antigen is known to protect against complement killing^29^ and O-antigen density has recently been directly linked to colicin protection in *E. coli* ST131^30^. In addition, complement sensitivity was observed in all NB and WB isolates that displayed either rough LPS or shorter O-antigen repeats.

It was evident that the phylogenetic position of individual isolates did not correlate with the binding phenotypes nor antibody function. This indicates that significant phenotypic variation can occur within highly clonal microbial populations. Similar data has been reported for other bacteria, including *Salmonella* Typhimurium^31^. However, the binding phenotypes correlated with both differences in phagocytosis and serum sensitivity, signifying the importance of having high-throughput methods to phenotypically screen large populations of isolates as genomic data alone cannot not always be predictive of a phenotype.

The majority of isolates screened in this study were resistant to complement, even in the presence of the KM467, and a significant proportion of the remaining susceptible isolates were protected from complement killing in the presence of increased KM467 concentrations. Antibody-mediated protection from complement has been observed previously^32,33,34^ and is a major concern when it comes to antibody therapy. This observation needs to be considered when targeting bacterial O-antigens by therapeutic mAbs. Our study also highlights the importance of screening candidate mAbs against a large panel of clinical isolates as there can be substantial phenotypic differences between genetically related organisms. Of note is that the NCTC13441 reference isolate is part of the minority SAB group and is perhaps not the best representative to be used for studies on *E. coli* ST131 O25b.

Compared to alternative mAb screening methods, HCI offers a much more comprehensive approach than ELISA or flow cytometry based assays. In terms of measuring antibody affinity, this is performed by measuring the intensity of the fluorescently-labelled antibody and does not provide any binding kinetic data (*K*_on_ or *K*_off_ values) that can be obtained by surface plasmon resonance. However, HCI can measure the binding of antibodies to native epitopes on intact bacteria, providing a more realistic *in vivo* representation. Phenotypic antibody screening, using an ELISA-based whole bacterial binding assay, was successfully used to discover a therapeutic mAb against the *Pseudomonas aeruginosa* exopolysaccharide Psl, a target which would have been missed using target-based screening methods^35^. A bispecific antibody MEDI3902, targeting Psl and PcrV (inhibiting type III secretion), recently completed phase 1 clinical trials^36,37^.

HCI can potentially be used to evaluate the binding efficacy and specificity of targeted mAbs against specific high-risk clones such as *E. coli* ST131, *Klebsiella pneumoniae* ST258 or *Staphylococcus aureus* ST22 (MRSA). This approach may be useful both in evaluating antibodies for their diagnostic potential, as well as for therapeutic use in combination with high-throughput opsonophagocytosis assays. It is also technically possible to screen different species simultaneously with the same antibody to evaluate cross-reactivity of broader spectrum targets including common surface antigens found on multiple pathogens. However, one of the potential advantages to more targeted mAbs is that these should not impact the host microbiota diversity, which is a major problem with antimicrobial usage.

Currently, HCI analysis software packages are optimised for mammalian cells and analysing bacteria this way has only just started being possible. We are hopeful that advances in image analysis and machine learning will soon enable more efficient segmentation and image analysis of bacteria. Our study was able to link phenotype with function, generating the possibility to be able to model the efficacy of a mAb based on the HCI phenotypic profile. Future application of HCI will likely not be limited to mAb screening, but also bispecific antibodies and antibody cocktails, as well as mAbs in combination with antimicrobials or novel antibacterial compounds.

## Materials and Methods

### Bacterial isolates

The *E. coli* reference strain NCTC13441 was obtained from the Public Health England NCTC culture collection. Clinical *E. coli* ST131 (Supplementary Table S1) were isolated between 2007 and 2015 at Addenbrooke’s Hospital, Cambridge, UK, and whole genome sequence data was previously published^38^. Bacteria were cultured on LB agar plates and single colonies picked for overnight culture in LB broth at 37°C at 200rpm.

### Antibody

DNA fragments encoding the VH and VL amino acid sequences for 3E9-11 were cloned into a plasmid vector for expression on a human IgG1 backbone and transfected into Chinese Hamster Ovary cell lines for transient expression and purification. For the phagocytosis experiments, the antibody directly conjugated to Alexa Fluor 647 using an Alexa Fluor 647 Antibody Labelling Kit (ThermoFisher).

### High-content antibody binding assay

Overnight bacterial cultures were diluted 1:200 in LB broth and 50μl bacterial suspension was added per well in CellCarrier Ultra 96 plates (Perkin Elmer) and left in a static incubator at 37°C for 2 hours. The supernatant and any non-adhered bacteria were removed and the remaining bacteria were fixed with 4% paraformaldehyde (PFA) in phosphate buffered saline (PBS) for 15 minutes at room temperature, then washed with PBS. KM467 was diluted to 1μg/ml in PBS+1% bovine serum albumin (BSA) and added to the plate for 1 hour (100μl per well) at room temperature, after which it was replaced by 100μl 2μg/ml Alexa Fluor 647 Goat Anti-Human IgG (ThermoFisher) plus 2μg/ml DAPI (ThermoFisher) in PBS+1% BSA for 30 min in the dark. The plate was washed once with PBS, then 50μl PBS was added to each well. The plates were imaged on the Opera Phenix using the Alexa Flour 647 and DAPI channels and the 63x water immersion objective and 16 fields and 5 Z-stacks were imaged per well.

### Opera Phenix image analysis

Image analysis was carried out using the Perkin Elmer Harmony version 4.8. Supplementary Table 2 shows the detailed analysis workflow and outputs. Exported image data was plotted in R.

### LPS analysis

LPS was extracted using a modified hot-phenol method^39^. Ten ml bacterial overnight cultures were centrifuged for 10 minutes at 9000 x g. The pellets were resuspended in 0.75 ml distilled water, then 0.5 ml 90% phenol was added. The mixture was vortexed then heated to 65 °C for 15 minutes, vortexing every 5 minutes. It was then cooled on ice after which the layers were separated by centrifugation at 9000 x g for 20 minutes at 4 °C. The aqueous layer was dialysed against distilled water in 6000–8000 Dalton (Da) molecular weight cut off dialysis tubing (Spectra/Por, Spectrum) for 48 hours, changing the water 5 times. 10μl of each LPS was mixed with 2x Novex™ Tris-Glycine SDS Sample Buffer and separated by gel electrophoresis for 1.5 hours at 30mA using Novex WedgeWell 12% Tris-Glycine Mini gels in Novex™ Tris-Glycine SDS Running Buffer (ThermoFisher). The gels were stained using sodium-m-periodate silver staining as described previously^40^.

### Immunoblotting

Five μl of each LPS was separated by electrophoresis as described above. LPS was transferred to 0.2 μm polyvinylidene difluoride (PVDF) membranes using the Trans-Blot Turbo blotting system (Bio-Rad) with a high-molecular-weight (MW) program (1.3A up to 25V for 10 min). The membranes were blocked overnight in 5% BSA in PBS plus 0.1% Tween 20 (PBST), then incubated with 1μg/ml KM467 or polyclonal anti-O25 typing sera (abcam, diluted 1:10) in 0.5% BSA in PBST for 2h at RT. The membranes were washed 3 × 10 min in PBST, then incubated with 0.1μg/ml HRP-conjugated Goat Anti-Human IgG H&L or HRP-conjugated Goat Anti-Rabbit IgG H&L (abcam) in 0.5% BSA in PBST for 1h at RT. The membranes were washed 6 x 10 min in PBST, then developed with SuperSignal West Pico PLUS (Thermo Scientific).

### Phylogenomic and SNP analysis

For SNP analysis, paired end reads from 86 *E. coli* ST131 O25b:H4 isolates were mapped to the reference sequence of *E. coli* strain Escherichia_coli_UPEC_ST131/NCTC13441 (accession number: GCA_900448475.1) using the RedDog mapping pipeline (V1beta.10.3), available at: http://github.com/katholt/reddog. For phylogenetic analyses, SNPs with confident homozygous allele calls (i.e. phred score >20) outside of phage regions identified using PHAST^41^, and repetitive regions were identified using mummer (v3.23)^42^ (784,141 bp), and all genomes were concatenated to produce an alignment of alleles at 23,924 variant sites. Any further recombinant regions were identified using Gubbins (v2.3.2)^43^ and excluded resulting in a final set of 3,300 SNPs identified from an alignment of 5,174,631 bp for the 86 isolates. Maximum likelihood (ML) phylogenetic trees were inferred from this alignment using RAxML (v8.2.9)^44^, with a generalized time-reversible model, a Gamma distribution to model site-specific rate variation (the GTR+ Γ substitution model; GTRGAMMA in RAxML), and 100 bootstrap pseudo-replicates to assess branch support. The resulting tree was visualized using iTOL^45^.

SNPs in the Quinolone Resistance Determining Region (QRDR) of genes *gyrA, and parC* were extracted from the whole genome SNP alignments.

### Gene content and mobile genetic element analysis

SRST2 (v0.2.0)^18^ was used to identify AMR genes (ARGannot^46^), *E. coli* serotypes (EcOH database^19^), as well as their precise alleles. ISMapper^23^ was run with default parameters to screen all read sets for the presence of the transposases IS*1* (accession number: X52534.1) and IS*Ec10* (accession number: WP_000174402.1) relative to the NCTC13441 reference chromosome sequence.

Raw read data was assembled *de novo* with Unicycler (v0.4.7)^20^, visualised with Bandage^47^ and annotated with PROKKA (v1.13.3)^48^, and operon alignment figures generated using EasyFig2.2.2^49^. Individual genes as well as operons were extracted manually and analysed using NCBI BLAST, MAUVE^22^ and Clustal Omega^50^.

### Electron microscopy

Bacterial colonies were picked off LB agar plates and placed into planchettes with Hexedecene for cryopreservation in a Baltec HPM010 high pressure freezer. Frozen samples were post-fixed by freeze-substitution as described previously^51^. Ultrathin section were cut on a Leica UC6 ultramicrotome, contrasted with uranyl acetate and lead citrate and imaged on an FEI Spirit Biotwin TEM using a Tietz F416 CCD. The thickness of the surface matrix was quantified directly at 60K magnification using EM Menu Measure at 20 locations from a total of 10 individual cells representing each isolate.

### Serum resistance

Bacterial overnights were diluted 1:1000 in LB and 50μl of each isolate was mixed with 25μl of Dulbecco’s Phosphate-Buffered Saline with calcium and magnesium (DPBS++) and 25μl baby rabbit complement serum (BRC) (Bio-Rad, C12CA, lot 148004). As controls, 50μl of the diluted bacterial culture mixed with 25μl DPBS++ and 25μl LB was used. Plates were shaken at 200rpm at 37°C and OD600 values were obtained by an automated plate reader (BMG Fluostar) after 4 hours. In parallel, 10-fold serial dilutions were plated in triplicate and colony forming units (cfu) were counted and the mean cfu/ml calculated.

### Antibody induced serum killing

Bacterial overnights were diluted 1:1000 in LB. The sample was split into 3 tubes and KM467 was added to two tubes at final concentrations of 2 μg/ml or 40 μg/ml. The third sample was left as a control without antibody. These were incubated for 30 minutes shaking at 200 RPM at 37°C degrees, then 50μl of these were mixed with 25μl DPBS++ and BRC, and assayed as described above.

### Phagocytosis and opsonophagocytosis assays

An average of 5×10^4^ THP1cells per well were differentiated in CellCarrier ultra 96 microplates with 10ng/ml phorbol-12-myristate-13 acetate (PMA) in RPMI + HEPES + heat inactivated foetal calf serum (HiFCS) (labtech) for 3 days after which the media was changed to RPMI + HEPES + HiFCS. Overnight bacterial cultures were adjusted to an OD600 of 1 and diluted 1:100 in LB and pre-incubated with either KM467 (final concentration of 4μg/ml) or LB (control). Bacteria were incubated for 1 hour at 200rpm at 37°C, after which 50µl was used to infect each well. Cells were infected for 90 minutes at 37°C, after which the infection media was aspirated and extracellular bacteria were removed by 3 × 50μl PBS washes. The plates were then fixed with 4% PFA. Macrophages were permeabilized with 0.1% Triton X-100 for 10 minutes followed blocking for 30 minutes with 10% BSA at room temperature in the dark. Bacteria were stained with 2μg/ml Alexa 647 labelled KM467 antibody or serum from mice immunised with *E. coli* MG1655 outer-membrane vesicles (provided by Kymab Ltd) followed by an anti-mouse IgG Alexa Fluor 647 secondary (abcam), diluted in 2% BSA for one hour in the dark. The cells were then stained with DAPI and CellMask Orange (Thermo) for 10 minutes in the dark. The plates were imaged on the Opera Phenix using the CellMask Orange, Alexa Flour 647 and DAPI channels and the 40x air objective, and 11 fields and 3 Z-stacks were imaged per well.

Opsonophagocytosis assays were performed in a same way as the phagocytosis assays, except the dilution of the overnights was done in LB with 10% BRC.

## Data availability

Whole genome sequenced raw read data is available at the European Nucleotide Archive (ENA) and individual sample accession numbers are listed in Supplementary Table S1. The image data analysed in this study can be found in Supplementary Table S3, and images generated during this study are available from the corresponding author on reasonable request.

## Acknowledgements

We thank Sally Forrest and Beth Blane for laboratory assistance, Sharon Peacock and Nicholas Brown for access to the bacterial isolates, and James Hutt for help with the image analysis pipeline. This work was supported by the MRC Proximity to Discovery: Industry Engagement Fund Biomedical Research Exchange Programme. MM and JBS are funded by the National Institute for Health Research (Cambridge Biomedical Research Centre at the Cambridge University Hospitals NHS Foundation Trust).

## Author Contributions

STR, PK, GD and JBS conceived the project. MM and JBS optimised the HCI screen and image data analysis and carried out most of the experiments. MM, ZAD and JBS carried out the bioinformatic analysis. SES managed the antibody production and validation at Kymab Ltd. TEM imaging was done by DAG. CL provided the *E. coli* ST131 isolates and sequencing data. SB assisted in data interpretation. JBS wrote the main manuscript and MM and JBS prepared the Figures. All authors reviewed the manuscript.

## Supplementary Information

### Direct TMB ELISA

High bind ELISA plates (Costar) were coated with 50μL of LPS from *E. coli* NCTC13441 or control LPS from *E. coli* O111:B4 (Sigma) in duplicate at an estimated 30μg/mL, 20μg/mL, 10μg/mL or 2.5 μg/mL in phosphate buffered saline (PBS) overnight at 4°C. Plates were washed 3x with PBS+0.1% Tween (PBST), blocked with PBS+1% bovine serum albumin (BSA) for 1h at RT, then washed again 3x with PBST. Serial 3-fold dilutions of KM467 (50μg/ml starting concentration) were prepared in PBS+1% BSA and 50ul per well was added to the coated wells and incubated for 1h at RT. Plates was washed 3x with PBST, after which 50μL of horseradish peroxidase (HRP) conjugated goat anti-human IgG diluted 1:3000 in PBS+1% BSA was added for 1 hr at RT. Plates was washed 3x with PBST, then 50μl of TMB substrate (Sigma) was added and incubated at RT protected from light. Then 50µl/well of stop solution (sulphuric acid, Fisher) was added and absorbance was read at 450nm (and correct at 540nm).

**Supplementary Figure S1: Titration of KM467 against *E. coli* ST131 NCTC13441 LPS by ELISA.** ELISA plates were coated with different amounts of isolated LPS from *E. coli* ST131 NCTC13441 O25b:H4 and control LPS from *E. coli* O111:B4, and serial 3-fold dilutions of KM467 (top concentration 50μg/ml) was added and binding was measured using TMB ELISA substrate at 405nm. Error bars represent standard deviation of 3 replicates.

**Supplementary Figure S2: Antibody binding titration.** *E. coli* ST131 O25b reference isolate NCTC13441 and an O16 clinical isolate (VREC0673) were incubated with different concentrations of KM467 (ranging from 0-20μg/ml). Bacteria were stained with DAPI and an Alexa Flour 647-conjugated secondary antibody (**a**). Percentage binding (**b**) was plotted against concentration. Error bars represent standard deviation of 4 replicates.

**Supplementary Figure S3: Antibody binding titration against different phenotypes confirms weak binding.** Representative isolates exhibiting different binding phenotype were incubated with different concentrations of KM467 (ranging from 0-20μg/ml). Bacteria were stained with DAPI and an Alexa Flour 647-conjugated secondary antibody. Images taken at 1μg/ml are shown (**a**). Percentage binding (**b**) was plotted against concentration. Error bars represent standard deviation of 3 replicates.

**Supplementary Figure S4: Silver stained LPS gels**. Isolated LPS from 35 isolates was analysed by gel electrophoresis followed by silver staining. Order on gel (1-35): VREC1101, VREC1106, VRES1160, VREC1403, VREC1428, VREC1592, VRES1610, VRES1619, VREC1630, NCTC13441, ECO0056, ECO0061, VREC0645, VREC0693, VRES0700, VREC0708, VRES0710, VRES0739, VREC0829, VREC0865, VREC0926, VREC1013, VREC1073, VRES1100, ECO0172, ECO0211, ECO0218, ECO0224, ECO0353, ECO0387, ECO0395, ECO0407, ECO0422, ECO0431, ECO0433. Any identified disrupted genes are indicated below gels.

**Supplementary Figure S5: Wxy, WekB, WbbL, RfaJ, RfaZ Clustal Omega amino acid alignments.**

**Supplementary Figure S6: Background differences in phagocytosis for isolates without antibody present.** Macrophages were stained with CellMasks Orange (cytoplasm) and DAPI (nucleic acid) and bacteria were visualised using KM467 labelled with Alexa Flour 647.

Images were obtained using a 40x air objective on the Opera Phenix. Images were analysed in Harmony using predefined building blocks to segment nuclei and cytoplasm to define the number of cells, and to count individual bacteria (using Find Spots) within the cells. Error bars represent standard deviation of 3 replicates.

**Supplementary Figure S7: Serum Resistance Reproducibility.** Growth in LB broth in the presence or absence of 25% BRC was measured by OD600 after 4 hours and plotted as percentage growth in the presence of serum compared to LB alone (**a**). Growth in LB broth in the presence or absence of 25% BRC was also measured by cfu counting after 4 hours (**b)** and **(c**). The average of representative isolates per phenotype (SB=4, SAB=4, WB=3, NB =3) of 3 replicates is shown, and error bars represent standard deviation. Significance was determined by *t*-test (*=<0.05, **= between 0.001 and 0.05, *** <0.001).

**Supplementary Table S1: List of isolates used in the study.**

**Supplementary Table S2: Antibody binding analysis pipeline (Harmony).**

**Supplementary Table S3: Image data from three replicate screens.**

**Supplementary Table S4: Presence of IS*Ec10* or IS*1* in genes realted to capsule, LPS core and O-antigen biosynthesis.** ISmapper results showing the presence (+) or absence (-) of IS*1* or IS*Ec10* in LPS or capsule biosynthesis genes. * presence of IS but an imprecise hit.

**Supplementary Table S5: Macrophage phagocytosis analysis pipeline (Harmony).**

## References

1. O’Neill. Review on Antimicrobial Resistance. Tackling drug-resistant infections globally: final report and recommendations. London, United Kingdom. (2016). Available at: https://amr-review.org/sites/default/files/160525_Finalpaper_withcover.pdf.

2. Alexander, H. E. Treatment of Haemophilus influenza infections and of Meningococcic meningitis. Am. J. Dis. Child. 66, 172–187 (1943).

3. Stiehm, R.E. Passive Immunization. in Textbook of pedriatrics infectious disease (ed. Feigin R.D., C. J. D.) 2769–2802 (The W.B Saunders CO. Philadelphia, Pa, 1998, 1998).

4. Singh, S. et al. Monoclonal Antibodies: A Review. Curr. Clin. Pharmacol. 13, 85–99 (2018).

5. The Impact-RSV Study Group. Palivizumab, a humanized respiratory syncytial virus monoclonal antibody, reduces hospitalization from respiratory syncytial virus infection in high-risk infants. Pediatrics 102, 531–537 (1998).

6. McConnell, M. J. Where are we with monoclonal antibodies for multidrug-resistant infections? Drug Discov. Today 24, 1132–1138 (2019).

7. Holden, M. T. G. et al. A genomic portrait of the emergence, evolution, and global spread of a methicillin-resistant Staphylococcus aureus pandemic. Genome Res. 23, 653–664 (2013).

8. Wong, V. K. et al. Phylogeographical analysis of the dominant multidrug-resistant H58 clade of Salmonella Typhi identifies inter- and intracontinental transmission events. Nat. Genet. 47, 632–639 (2015).

9. Wyres, K. L. & Holt, K. E. Klebsiella pneumoniae Population Genomics and Antimicrobial-Resistant Clones. Trends Microbiol. 24, 944–956 (2016).

10. Petty, N. K. et al. Global dissemination of a multidrug resistant Escherichia coli clone. Proc. Natl. Acad. Sci. U. S. A. 111, 5694–5699 (2014).

11. Szijártó, V. et al. Bactericidal monoclonal antibodies specific to the lipopolysaccharide O antigen from multidrug-resistant Escherichia coli clone ST131-O25b:H4 elicit protection in mice. Antimicrob. Agents Chemother. 59, 3109–3116 (2015).

12. Guachalla, L. M. et al. Multiple Modes of Action of a Monoclonal Antibody against Multidrug-Resistant Escherichia coli Sequence Type 131-H30. Antimicrob. Agents Chemother. 61, e01428–17 (2017).

13. van Vliet, E. et al. Current approaches and future role of high content imaging in safety sciences and drug discovery. ALTEX 31, 479–493 (2014).

14. Bray, M.-A. et al. Cell Painting, a high-content image-based assay for morphological profiling using multiplexed fluorescent dyes. Nat. Protoc. 11, 1757–1774 (2016).

15. Christophe, T., Ewann, F., Jeon, H. K., Cechetto, J. & Brodin, P. High-content imaging of Mycobacterium tuberculosis-infected macrophages: an in vitro model for tuberculosis drug discovery. Future Med. Chem. 2, 1283–1293 (2010).

16. Greenwood, D. J. et al. Subcellular antibiotic visualization reveals a dynamic drug reservoir in infected macrophages. Science 364, 1279–1282 (2019).

17. Zoffmann, S. et al. Machine learning-powered antibiotics phenotypic drug discovery. Sci. Rep. 9, 5013 (2019).

18. Inouye, M. et al. SRST2: Rapid genomic surveillance for public health and hospital microbiology labs. Genome Med. 6, 90 (2014).

19. Ingle, D. J. et al. In silico serotyping of E. coli from short read data identifies limited novel O-loci but extensive diversity of O:H serotype combinations within and between pathogenic lineages. Microb. genomics 2, e000064–e000064 (2016).

20. Wick, R. R., Judd, L. M., Gorrie, C. L. & Holt, K. E. Unicycler: Resolving bacterial genome assemblies from short and long sequencing reads. PLoS Comput. Biol. 13, e1005595–e1005595 (2017).

21. Szijártó, V. et al. Diagnostic potential of monoclonal antibodies specific to the unique O-antigen of multidrug-resistant epidemic Escherichia coli clone ST131-O25b:H4. Clin. Vaccine Immunol. 21, 930–939 (2014).

22. Darling, A. C. E., Mau, B., Blattner, F. R. & Perna, N. T. Mauve: Multiple alignment of conserved genomic sequence with rearrangements. Genome Res. 14, 1394–1403 (2004).

23. Hawkey, J. et al. ISMapper: identifying transposase insertion sites in bacterial genomes from short read sequence data. BMC Genomics 16, 667 (2015).

24. Yethon, J. A., Vinogradov, E., Perry, M. B. & Whitfield, C. Mutation of the lipopolysaccharide core glycosyltransferase encoded by waaG destabilizes the outer membrane of Escherichia coli by interfering with core phosphorylation. J. Bacteriol. 182, 5620–5623 (2000).

25. Pagnout, C. et al. Pleiotropic effects of rfa-gene mutations on Escherichia coli envelope properties. Sci. Rep. 9, (2019).

26. Phan, M.-D. et al. The serum resistome of a globally disseminated multidrug resistant uropathogenic Escherichia coli clone. PLoS Genet. 9, e1003834–e1003834 (2013).

27. Ranjan, A. et al. ESBL-plasmid carriage in E. coli enhances in vitro bacterial competition fitness and serum resistance in some strains of pandemic sequence types without overall fitness cost. Gut Pathog. 10, 24 (2018).

28. Itzek, A., Chen, Z., Merritt, J. & Kreth, J. Effect of salivary agglutination on oral streptococcal clearance by human polymorphonuclear neutrophil granulocytes. Mol. Oral Microbiol. 32, 197–210 (2017).

29. Sarkar, S., Ulett, G. C., Totsika, M., Phan, M. D. & Schembri, M. A. Role of capsule and O antigen in the virulence of uropathogenic Escherichia coli. PLoS One 9, (2014).

30. Sharp, C. et al. O-antigen-dependent colicin insensitivity of uropathogenic Escherichia coli. J. Bacteriol. 201, (2019).

31. Ondari, E. M. et al. Rapid transcriptional responses to serum exposure are associated with sensitivity and resistance to antibody-mediated complement killing in invasive salmonella typhimurium st313 [version 1; peer review: 2 approved]. Wellcome Open Res. 4, (2019).

32. Goh, Y. S. et al. Bactericidal Immunity to Salmonella in Africans and Mechanisms Causing Its Failure in HIV Infection. PLoS Negl. Trop. Dis. 10, e0004604 (2016).

33. Wells, T. J. et al. Increased severity of respiratory infections associated with elevated anti-LPS IgG2 which inhibits serum bactericidal killing. J. Exp. Med. 211, 1893–1904 (2014).

34. Coggon, C. F. et al. A novel method of serum resistance by Escherichia coli that causes urosepsis. MBio 9, (2018).

35. DiGiandomenico, A. et al. Identification of broadly protective human antibodies to Pseudomonas aeruginosa exopolysaccharide Psl by phenotypic screening. J. Exp. Med. 209, 1273–1287 (2012).

36. DiGiandomenico, A. et al. A multifunctional bispecific antibody protects against Pseudomonas aeruginosa. Sci. Transl. Med. 6, 262ra155–262ra155 (2014).

37. Ali, S. O. et al. Phase 1 study of MEDI3902, an investigational anti-Pseudomonas aeruginosa PcrV and Psl bispecific human monoclonal antibody, in healthy adults. Clin. Microbiol. Infect. 25, 629.e1-629.e6 (2019).

38. Ludden, C. et al. One Health Genomic Surveillance of Escherichia coli Demonstrates Distinct Lineages and Mobile Genetic Elements in Isolates from Humans versus Livestock. MBio 10, e02693–18 (2019).

39. Westphal, O. J., Jann, K. Bacterial lipopolysaccharides:extraction with phenol-water and further applications of the procedure. Methods Carbohydr. Chem. 5, 83–91 (1965).

40. Tsai, C. M. & Frasch, C. E. A sensitive silver stain for detecting lipopolysaccharides in polyacrylamide gels. Anal. Biochem. 119, 115–119 (1982).

41. Zhou, Y., Liang, Y., Lynch, K. H., Dennis, J. J. & Wishart, D. S. PHAST: A Fast Phage Search Tool. Nucleic Acids Res. 39, W347–W352 (2011).

42. Kurtz, S. et al. Versatile and open software for comparing large genomes. Genome Biol. 5, R12–R12 (2004).

43. Croucher, N. J. et al. Rapid phylogenetic analysis of large samples of recombinant bacterial whole genome sequences using Gubbins. Nucleic Acids Res. 43, e15–e15 (2015).

44. Stamatakis, A. RAxML-VI-HPC: maximum likelihood-based phylogenetic analyses with thousands of taxa and mixed models. Bioinformatics 22, 2688–2690 (2006).

45. Letunic, I. & Bork, P. Interactive Tree Of Life (iTOL): An online tool for phylogenetic tree display and annotation. Bioinformatics 23, 127–128 (2007).

46. Gupta, S. K. et al. ARG-ANNOT, a new bioinformatic tool to discover antibiotic resistance genes in bacterial genomes. Antimicrob. Agents Chemother. 58, 212–20 (2014).

47. Wick, R. R., Schultz, M. B., Zobel, J. & Holt, K. E. Bandage: interactive visualization of de novo genome assemblies. Bioinformatics 31, 3350–2 (2015).

48. Seemann, T. Prokka: rapid prokaryotic genome annotation. Bioinformatics 30, 2068–2069 (2014).

49. Sullivan, M. j., Petty, n.K & Beatson, S. A. Easyfig: a genome comparison visualizer. Available at: https://www.ncbi.nlm.nih.gov/pmc/articles/PMC3065679/. (Accessed: 6th March 2020)

50. Madeira, F. et al. The EMBL-EBI search and sequence analysis tools APIs in 2019. Nucleic Acids Res. 47, W636–W641 (2019).

51. Dorman, M. J., Feltwell, T., Goulding, D. A., Parkhill, J. & Short, F. L. The capsule regulatory network of klebsiella pneumoniae defined by density-traDISort. MBio 9, (2018).

