## Supplementary Fig. S1 for "A novel therapeutic antibody screening method using bacterial high-content imaging reveals functional antibody binding phenotypes of *Escherichia coli* ST131"

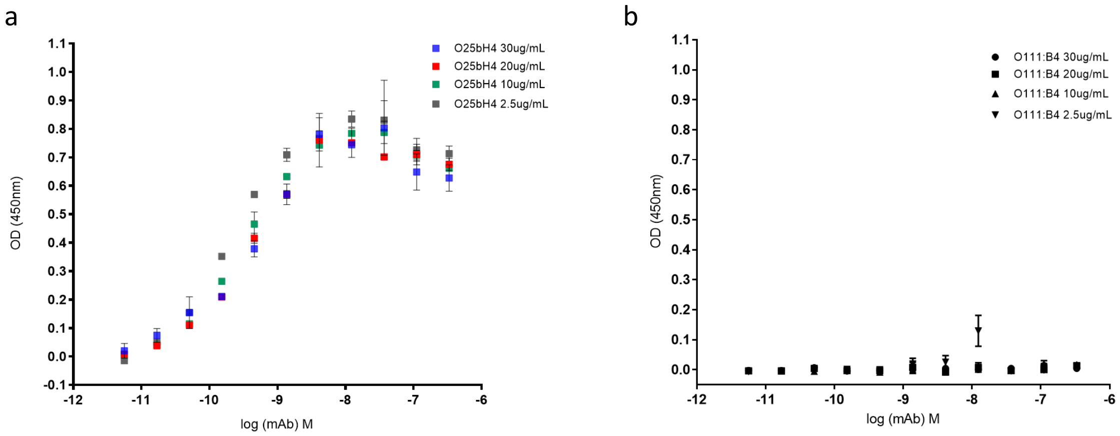


**Supplementary Figure S1: Titration of KM467 against *E. coli* ST131 NCTC13441 LPS by ELISA.** ELISA plates were coated with different amounts of isolated LPS from *E. coli* ST131 NCTC13441 O25b:H4 and control LPS from *E. coli* O111:B4, and serial 3-fold dilutions of KM467 (top concentration 50µg/ml) was added and binding was measured using TMB ELISA substrate at 405nm. Error bars represent standard deviation of 3 replicates.
