## Supplementary Fig. S2 for "A novel therapeutic antibody screening method using bacterial high-content imaging reveals functional antibody binding phenotypes of *Escherichia coli* ST131"

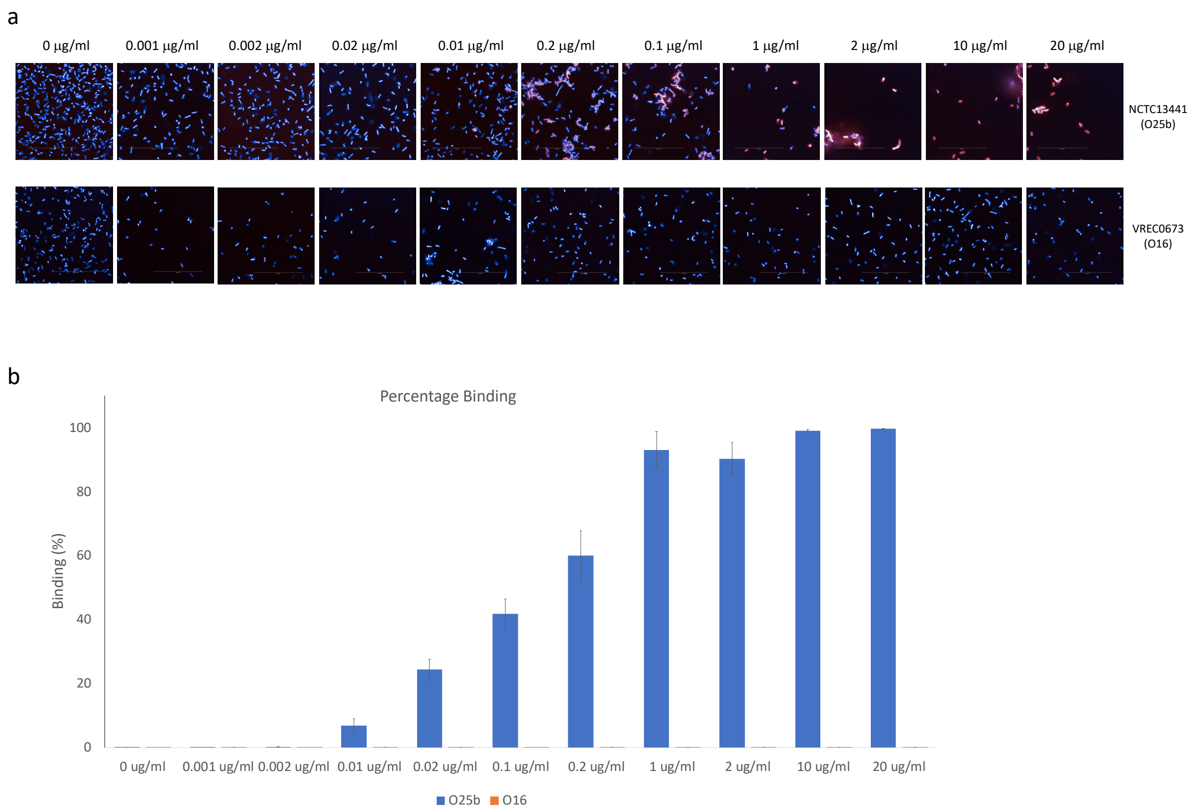


**Supplementary Figure S2: Antibody binding titration.** *E. coli* ST131 O25b reference isolate NCTC13441 and an O16 clinical isolate (VREC0673) were incubated with different concentrations of KM467 (ranging from 0-20µg/ml). Bacteria were stained with DAPI and an Alexa Flour 647-conjugated secondary antibody (**a**). Percentage binding was plotted against antibody concentration (**b**). Error bars represent standard deviation of 4 replicates.
