## Supplementary Fig. S3 for "A novel therapeutic antibody screening method using bacterial high-content imaging reveals functional antibody binding phenotypes of *Escherichia coli* ST131"

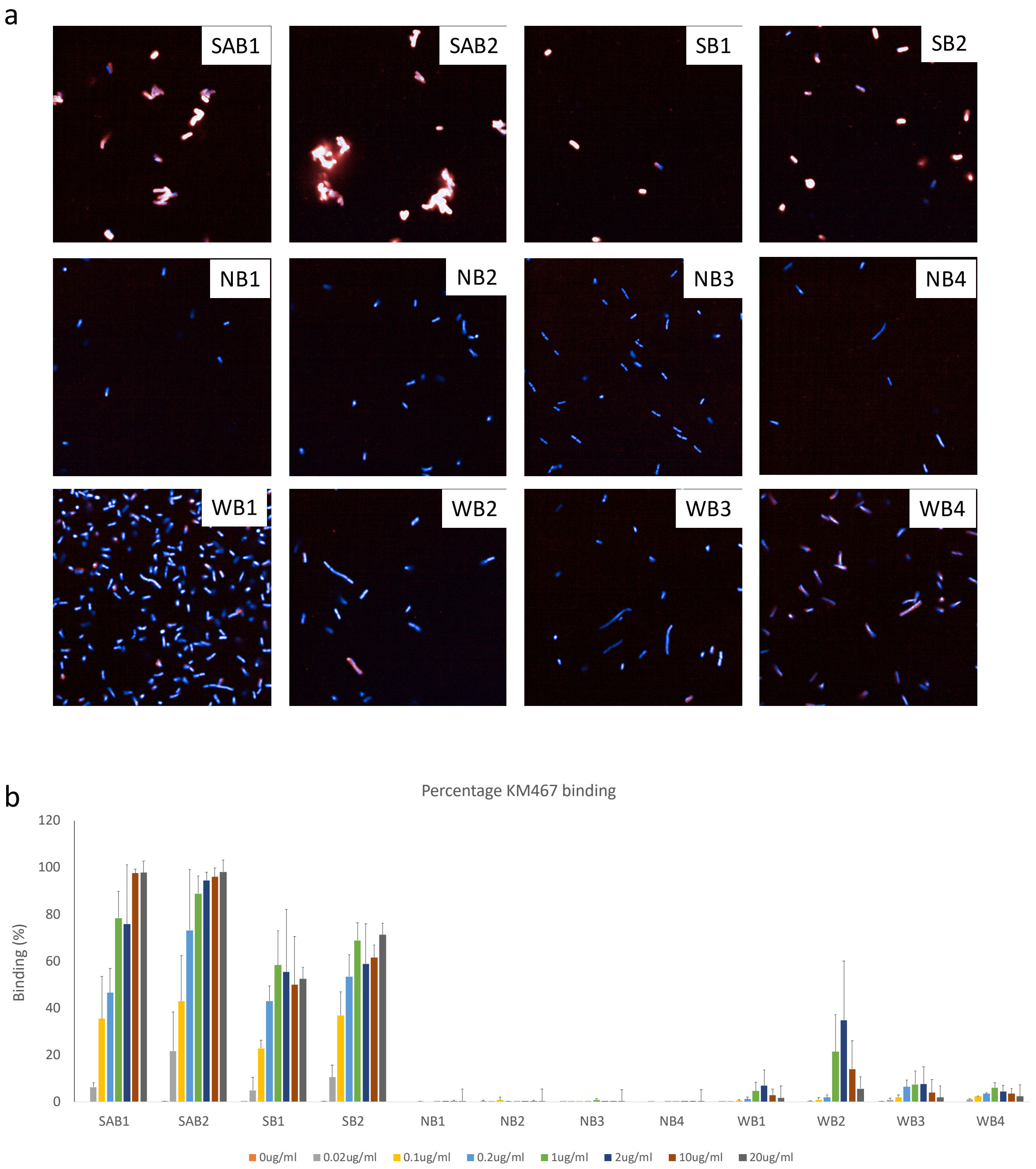


**Supplementary Figure S3: Antibody binding titration against different phenotypes confirms weak binding.** Representative isolates exhibiting different binding phenotype were incubated with different concentrations of KM467 (ranging from 0-20µg/ml). Bacteria were stained with DAPI and an Alexa Flour 647-conjugated secondary antibody. Images taken at 1µg/ml are shown (**a**). Percentage binding (**b**) was plotted against concentration. Error bars represent standard deviation of 3 replicates.
