## Supplementary Fig. S4 for "A novel therapeutic antibody screening method using bacterial high-content imaging reveals functional antibody binding phenotypes of *Escherichia coli* ST131"

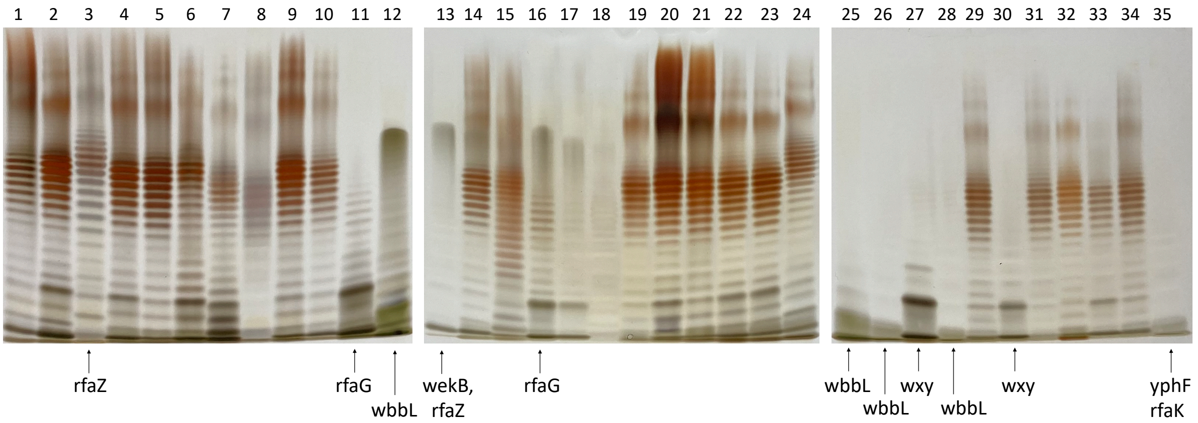


**Supplementary Figure S4: Silver stained LPS gels**. Isolated LPS from 35 isolates was analysed by gel electrophoresis followed by silver staining. Order on gel (1-35): VREC1101, VREC1106, VRES1160, VREC1403, VREC1428, VREC1592, VRES1610, VRES1619, VREC1630, NCTC13441, ECO0056, ECO0061, VREC0645, VREC0693, VRES0700, VREC0708, VRES0710, VRES0739, VREC0829, VREC0865, VREC0926, VREC1013, VREC1073, VRES1100, ECO0172, ECO0211, ECO0218, ECO0224, ECO0353, ECO0387, ECO0395, ECO0407, ECO0422, ECO0431, ECO0433. Any identified disrupted genes are indicated below gels.
