## Supplementary Fig. S5 for "A novel therapeutic antibody screening method using bacterial high-content imaging reveals functional antibody binding phenotypes of *Escherichia coli* ST131"

**Supplementary Figure 5: Wxy, WekB, WbbL, RfaJ, RfaZ Clustal Omega amino acid alignments.**

CLUSTAL O(1.2.4) multiple sequence alignment **WbbL**

```
ECO0211        MVYIIIVSHGHDDYIENLLLNLKLPSGRFKIIVRDNKSSMVLKKTCEKNCVTYLHGGQYE  60
NCTC13441      MVYIIIVSHGHDDYIENLLLNLKLPSGRFKIIVRDNKSSMVLKKTCEKNCVTYLHGGQYG  60
VREC1403       MVYIIIVSHGHDDYIENLLLNLKLPSGRFKIIVRDNKSSMVLKKTCEKNCVTYLHGGQYG  60
ECO0218        MVYIIIVSHGHDDYIENLLLNLKLPSGRFKIIVRDNKSSMVLKKTCEKNCVTYLHGGQYG  60
ECO0061        MVYIIIVSHGHDDYIENLLLNLKLPSGRFKIIVRDNKSSMVLKKTCEKNCVTYLHGGQYG  60
ECO0387        MVYIIIVSHGHDDYIENLLLNLKLPSGRFKIIVRDNKSSMVLKKTCEKNCVTYLHGGQYG  60
VRES1610       MVYIIIVSHGHDDYIENLLLNLKLPSGRFKIIVRDNKSSMVLKKTCEKNCVTYLHGGQYG  60
VRES1619       MVYIIIVSHGHDDYIENLLLNLKLPSGRFKIIVRDNKSSMVLKKTCEKNCVTYLHGGQYG  60
VRES0739       MVYIIIVSHGHDDYIENLLLNLKLPSGRFKIIVRDNKSSMVLKKTCEKNCVTYLHGGQYG  60
VREC0645       MVYIIIVSHGHDDYIENLLLNLKLPSGRFKIIVRDNKSSMVLKKTCEKNCVTYLHGGQYG  60
VRES1160       MVYIIIVSHGHDDYIENLLLNLKLPSGRFKIIVRDNKSSMVLKKTCEKNCVTYLHGGQYG  60
VRES0700       MVYIIIVSHGHDDYIENLLLNLKLPSGRFKIIVRDNKSSMVLKKTCEKNCVTYLHGGQYG  60
VREC1630       MVYIIIVSHGHDDYIENLLLNLKLPSGRFKIIVRDNKSSMVLKKTCEKNCVTYLHGGQYG  60
VREC1428       MVYIIIVSHGHDDYIENLLLNLKLPSGRFKIIVRDNKSSMVLKKTCEKNCVTYLHGGQYG  60
VREC1106       MVYIIIVSHGHDDYIENLLLNLKLPSGRFKIIVRDNKSSMVLKKTCEKNCVTYLHGGQYG  60
VREC1101       MVYIIIVSHGHDDYIENLLLNLKLPSGRFKIIVRDNKSSMVLKKTCEKNCVTYLHGGQYG  60
VREC1073       MVYIIIVSHGHDDYIENLLLNLKLPSGRFKIIVRDNKSSMVLKKTCEKNCVTYLHGGQYG  60
VREC1013       MVYIIIVSHGHDDYIENLLLNLKLPSGRFKIIVRDNKSSMVLKKTCEKNCVTYLHGGQYG  60
VREC0926       MVYIIIVSHGHDDYIENLLLNLKLPSGRFKIIVRDNKSSMVLKKTCEKNCVTYLHGGQYG  60
VREC0865       MVYIIIVSHGHDDYIENLLLNLKLPSGRFKIIVRDNKSSMVLKKTCEKNCVTYLHGGQYG  60
VREC0829       MVYIIIVSHGHDDYIENLLLNLKLPSGRFKIIVRDNKSSMVLKKTCEKNCVTYLHGGQYG  60
VREC0693       MVYIIIVSHGHDDYIENLLLNLKLPSGRFKIIVRDNKSSMVLKKTCEKNCVTYLHGGQYG  60
ECO0433        MVYIIIVSHGHDDYIENLLLNLKLPSGRFKIIVRDNKSSMVLKKTCEKNCVTYLHGGQYG  60
ECO0224        MVYIIIVSHGHDDYIENLLLNLKLPSGRFKIIVRDNKSSMVLKKTCEKNCVTYLHGGQYG  60
ECO0411        MVYIIIVSHGHDDYIENLLLNLKLPSGRFKIIVRDNKSSMVLKKTCEKNCVTYLHGGQYG  60
ECO0431        MVYIIIVSHGHDDYIENLLLNLKLPSGRFKIIVRDNKSSMVLKKTCEKNCVTYLHGGQYG  60
ECO0353        MVYIIIVSHGHDDYIENLLLNLKLPSGRFKIIVRDNKSSMVLKKTCEKNCVTYLHGGQYG  60
               ***********************************************************

ECO0211        FGHNNNIAVSYIINNFMIMNNDYFLFLNPDVFITSESLINYVDYIISNDYKFSTLCLYRD  120
NCTC13441      FGHNNNIAVSYIINNFMIMNNDYFLFLNPDVFITSESLINYVDYIISNDYKFSTLCLYRD  120
VREC1403       FGHNNNIAVSYIINNFMIMNNDYFLFLNPDVFITSESLINYVDYIISNDYKFSTLCLYRD  120
ECO0218        FGHNNNIAVSYIINNFMIMNNDYFLFLNPDVFITSESLINYVDYIISNDYKFSTLCLYRD  120
ECO0061        FGHNNNIAVSYIINNFMIMNNDYFLFLNPDVFITSESLINYVDYIISNDYKFSTLCLYRD  120
ECO0387        FGHNNNIAVSYIINNFMIMNNDYFLFLNPDVFITSESLINYVDYIISNDYKFSTLCLYRD  120
VRES1610       FGHNNNIAVSYIINNFMIMNNDYFLFLNPDVFITSESLINYVDYIISNDYKFSTLCLYRD  120
VRES1619       FGHNNNIAVSYIINNFMIMNNDYFLFLNPDVFITSESLINYVDYIISNDYKFSTLCLYRD  120
VRES0739       FGHNNNIAVSYIINNFMIMNNDYFLFLNPDVFITSESLINYVDYIISNDYKFSTLCLYRD  120
VREC0645       FGHNNNIAVSYIINNFMIMNNDYFLFLNPDVFITSESLINYVDYIISNDYKFSTLCLYRD  120
VRES1160       FGHNNNIAVSYIINNFMIMNNDYFLFLNPDVFITSESLINYVDYIISNDYKFSTLCLYRD  120
VRES0700       FGHNNNIAVSYIINNFMIMNNDYFLFLNPDVFITSESLINYVDYIISNDYKFSTLCLYRD  120
VREC1630       FGHNNNIAVSYIINNFMIMNNDYFLFLNPDVFITSESLINYVDYIISNDYKFSTLCLYRD  120
VREC1428       FGHNNNIAVSYIINNFMIMNNDYFLFLNPDVFITSESLINYVDYIISNDYKFSTLCLYRD  120
VREC1106       FGHNNNIAVSYIINNFMIMNNDYFLFLNPDVFITSESLINYVDYIISNDYKFSTLCLYRD  120
VREC1101       FGHNNNIAVSYIINNFMIMNNDYFLFLNPDVFITSESLINYVDYIISNDYKFSTLCLYRD  120
VREC1073       FGHNNNIAVSYIINNFMIMNNDYFLFLNPDVFITSESLINYVDYIISNDYKFSTLCLYRD  120
VREC1013       FGHNNNIAVSYIINNFMIMNNDYFLFLNPDVFITSESLINYVDYIISNDYKFSTLCLYRD  120
VREC0926       FGHNNNIAVSYIINNFMIMNNDYFLFLNPDVFITSESLINYVDYIISNDYKFSTLCLYRD  120
VREC0865       FGHNNNIAVSYIINNFMIMNNDYFLFLNPDVFITSESLINYVDYIISNDYKFSTLCLYRD  120
VREC0829       FGHNNNIAVSYIINNFMIMNNDYFLFLNPDVFITSESLINYVDYIISNDYKFSTLCLYRD  120
VREC0693       FGHNNNIAVSYIINNFMIMNNDYFLFLNPDVFITSESLINYVDYIISNDYKFSTLCLYRD  120
ECO0433        FGHNNNIAVSYIINNFMIMNNDYFLFLNPDVFITSESLINYVDYIISNDYKFSTLCLYRD  120
ECO0224        FGHNNNIAVSYIINNFMIMNNDYFLFLNPDVFITSESLINYVDYIISNDYKFSTLCLYRD  120
ECO0411        FGHNNNIAVSYIINNFMIMNNDYFLFLNPDVFITSESLINYVDYIISNDYKFSTLCLYRD  120
ECO0431        FGHNNNIAVSYIINNFMIMNNDYFLFLNPDVFITSESLINYVDYIISNDYKFSTLCLYRD  120
ECO0353        FGHNNNIAVSYIINNFMIMNNDYFLFLNPDVFITSESLINYVDYIISNDYKFSTLCLYRD  120
               ************************************************************
```

```
ECO0211         FTKSKHDYSIRSFPTLYDFLCSFLLGVNKSKIKKENILSDTVVDWCAGSFMLIHALSFLN  180
NCTC13441       FTKSKHDYSIRSFPTLYDFLCSFLLGVNKSKIKKENILSDTVVDWCAGSFMLIHALSFLN  180
VREC1403        FTKSKHDYSIRSFPTLYDFLCSFLLGVNKSKIKKENILSDTVVDWCAGSFMLIHALSFLN  180
ECO0218         FTKSKHDYSIRSFPTLYDFLCSFLLGVNKSKIKKENILSDTVVDWCAGSFMLIHALSFLN  180
ECO0061         FTKSKHDYSIRSFPTLYDFLCSFLLGVNKSKIKKENILSDTVVDWCAGSFMLIHALSFLN  180
ECO0387         FTKSKHDYSIRSFPTLYDFLCSFLLGVNKSKIKKENILSDTVVDWCAGSFMLIHALSFLN  180
VRES1610        FTKSKHDYSIRSFPTLYDFLCSFLLGVNKSKIKKENILSDTVVDWCAGSFMLIHALSFLN  180
VRES1619        FTKSKHDYSIRSFPTLYDFLCSFLLGVNKSKIKKENILSDTVVDWCAGSFMLIHALSFLN  180
VRES0739        FTKSKHDYSIRSFPTLYDFLCSFLLGVNKSKIKKENILSDTVVDWCAGSFMLIHALSFLN  180
VREC0645        FTKSKHDYSIRSFPTLYDFLCSFLLGVNKSKIKKENILSDTVVDWCAGSFMLIHALSFLN  180
VRES1160        FTKSKHDYSIRSFPTLYDFLCSFLLGVNKSKIKKENILSDTVVDWCAGSFMLIHALSFLN  180
VRES0700        FTKSKHDYSIRSFPTLYDFLCSFLLGVNKSKIKKENILSDTVVDWCAGSFMLIHALSFLN  180
VREC1630        FTKSKHDYSIRSFPTLYDFLCSFLLGVNKSKIKKENILSDTVVDWCAGSFMLIHALSFLN  180
VREC1428        FTKSKHDYSIRSFPTLYDFLCSFLLGVNKSKIKKENILSDTVVDWCAGSFMLIHALSFLN  180
VREC1106        FTKSKHDYSIRSFPTLYDFLCSFLLGVNKSKIKKENILSDTVVDWCAGSFMLIHALSFLN  180
VREC1101        FTKSKHDYSIRSFPTLYDFLCSFLLGVNKSKIKKENILSDTVVDWCAGSFMLIHALSFLN  180
VREC1073        FTKSKHDYSIRSFPTLYDFLCSFLLGVNKSKIKKENILSDTVVDWCAGSFMLIHALSFLN  180
VREC1013        FTKSKHDYSIRSFPTLYDFLCSFLLGVNKSKIKKENILSDTVVDWCAGSFMLIHALSFLN  180
VREC0926        FTKSKHDYSIRSFPTLYDFLCSFLLGVNKSKIKKENILSDTVVDWCAGSFMLIHALSFLN  180
VREC0865        FTKSKHDYSIRSFPTLYDFLCSFLLGVNKSKIKKENILSDTVVDWCAGSFMLIHALSFLN  180
VREC0829        FTKSKHDYSIRSFPTLYDFLCSFLLGVNKSKIKKENILSDTVVDWCAGSFMLIHALSFLN  180
VREC0693        FTKSKHDYSIRSFPTLYDFLCSFLLGVNKSKIKKENILSDTVVDWCAGSFMLIHALSFLN  180
ECO0433         FTKSKHDYSIRSFPTLYDFLCSFLLGVNKSKIKKENILSDTVVDWCAGSFMLIHALSFLN  180
ECO0224         FTKSKHDYSIRSFPTLYDFLCSFLLGVNKSKIKKENILSDTVVDWCAGSFMLIHALSFLN  180
ECO0411         FTKSKHDYSIRSFPTLYDFLCSFLLGVNKSKIKKENILSDTVVDWCAGSFMLIHALSFLN  180
ECO0431         FTKSKHDYSIRSFPTLYDFLCSFLLGVNKSKIKKENILSDTVVDWCAGSFMLIHALSFLN  180
ECO0353         FTKSKHDYSIRSFPTLYDFLCSFLLGVNKSKIKKENILSDTVVDWCAGSFMLIHALSFLN  180
                ************************************************************

ECO0211         VNGFDQKYFMYCEDIDLCMRLKLSGVDLYYTPHFDAIHYAQHENRRIFTKAFRWHIRSIT  240
NCTC13441       VNGFDQKYFMYCEDIDLCMRLKLSGVDLYYTPHFDAIHYAQHENRRIFTKAFRWHIRSIT  240
VREC1403        VNGFDQKYFMYCEDIDLCMRLKLSGVDLYYTPHFDAIHYAQHENRRIFTKAFRWHIRSIT  240
ECO0218         VNGFDQKYFMYCEDIDLCMRLKLSGVDLYYTPHFDAIHYAQHENRRIFTKAFRWHIRSIT  240
ECO0061         VNGFDQKYFMYCEDIDLCMRLKLSGVDLYYTPHFDAIHYAQHENRRIFTKAF--------  232
ECO0387         VNGFDQKYFMYCEDIDLCMRLKLSGVDLYYTPHFDAIHYAQHENRRIFTKAFRWHIRSIT  240
VRES1610        VNGFDQKYFMYCEDIDLCMRLKLSGVDLYYTPHFDAIHYAQHENRRIFTKAFRWHIRSIT  240
VRES1619        VNGFDQKYFMYCEDIDLCMRLKLSGVDLYYTPHFDAIHYAQHENRRIFTKAFRWHIRSIT  240
VRES0739        VNGFDQKYFMYCEDIDLCMRLKLSGVDLYYTPHFDAIHYAQHENRRIFTKAFRWHIRSIT  240
VREC0645        VNGFDQKYFMYCEDIDLCMRLKLSGVDLYYTPHFDAIHYAQHENRRIFTKAFRWHIRSIT  240
VRES1160        VNGFDQKYFMYCEDIDLCMRLKLSGVDLYYTPHFDAIHYAQHENRRIFTKAFRWHIRSIT  240
VRES0700        VNGFDQKYFMYCEDIDLCMRLKLSGVDLYYTPHFDAIHYAQHENRRIFTKAFRWHIRSIT  240
VREC1630        VNGFDQKYFMYCEDIDLCMRLKLSGVDLYYTPHFDAIHYAQHENRRIFTKAFRWHIRSIT  240
VREC1428        VNGFDQKYFMYCEDIDLCMRLKLSGVDLYYTPHFDAIHYAQHENRRIFTKAFRWHIRSIT  240
VREC1106        VNGFDQKYFMYCEDIDLCMRLKLSGVDLYYTPHFDAIHYAQHENRRIFTKAFRWHIRSIT  240
VREC1101        VNGFDQKYFMYCEDIDLCMRLKLSGVDLYYTPHFDAIHYAQHENRRIFTKAFRWHIRSIT  240
VREC1073        VNGFDQKYFMYCEDIDLCMRLKLSGVDLYYTPHFDAIHYAQHENRRIFTKAFRWHIRSIT  240
VREC1013        VNGFDQKYFMYCEDIDLCMRLKLSGVDLYYTPHFDAIHYAQHENRRIFTKAFRWHIRSIT  240
VREC0926        VNGFDQKYFMYCEDIDLCMRLKLSGVDLYYTPHFDAIHYAQHENRRIFTKAFRWHIRSIT  240
VREC0865        VNGFDQKYFMYCEDIDLCMRLKLSGVDLYYTPHFDAIHYAQHENRRIFTKAFRWHIRSIT  240
VREC0829        VNGFDQKYFMYCEDIDLCMRLKLSGVDLYYTPHFDAIHYAQHENRRIFTKAFRWHIRSIT  240
VREC0693        VNGFDQKYFMYCEDIDLCMRLKLSGVDLYYTPHFDAIHYAQHENRRIFTKAFRWHIRSIT  240
ECO0433         VNGFDQKYFMYCEDIDLCMRLKLSGVDLYYTPHFDAIHYAQHENRRIFTKAFRWHIRSIT  240
ECO0224         VNGFDQKYFMYCEDIDLCMRLKLSGVDLYYTPHFDAIHYAQHENRRIFTKAFR-------  233
ECO0411         VNGFDQKYFMYCEDIDLCMRLKLSGVDLYYTPHFDAIHYAQHENRRIFTKAFRWHIRSIT  240
ECO0431         VNGFDQKYFMYCEDIDLCMRLKLSGVDLYYTPHFDAIHYAQHENRRIFTKAFRWHIRSIT  240
ECO0353         VNGFDQKYFMYCEDIDLCMRLKLSGVDLYYTPHFDAIHYAQHENRRIFTKAFRWHIRSIT  240
                ****************************************************

ECO0211         RYILRKPILSYKNYRKITSELVK 263
NCTC13441       RYILRKPILSYKNYRKITSELVK 263
VREC1403        RYILRKPILSYKNYRKITSELVK 263
ECO0218         RYILRKPILSYKNYRKITSELVK 263
ECO0061         ----------------------- 232
ECO0387         RYILRKPILSYKNYRKITSELVK 263
VRES1610        RYILRKPILSYKNYRKITSELVK 263
VRES1619        RYILRKPILSYKNYRKITSELVK 263
VRES0739        RYILRKPILSYKNYRKITSELVK 263
VREC0645        RYILRKPILSYKNYRKITSELVK 263
VRES1160        RYILRKPILSYKNYRKITSELVK 263
VRES0700        RYILRKPILSYKNYRKITSELVK 263
VREC1630        RYILRKPILSYKNYRKITSELVK 263
VREC1428        RYILRKPILSYKNYRKITSELVK 263
VREC1106        RYILRKPILSYKNYRKITSELVK 263
VREC1101        RYILRKPILSYKNYRKITSELVK 263
VREC1073        RYILRKPILSYKNYRKITSELVK 263
VREC1013        RYILRKPILSYKNYRKITSELVK 263
VREC0926        RYILRKPILSYKNYRKITSELVK 263
```

```
VREC0865      RYILRKPILSYKNYRKITSELVK263
VREC0829      RYILRKPILSYKNYRKITSELVK263
VREC0693      RYILRKPILSYKNYRKITSELVK263
ECO0433       RYILRKPILSYKNYRKITSELVK263
ECO0224       -----------------------233
ECO0411       RYILRKPILSYKNYRKITSELVK263
ECO0431       RYILRKPILSYKNYRKITSELVK263
ECO0353       RYILRKPILSYKNYRKITSELVK263
```

CLUSTAL O(1.2.4) multiple sequence alignment - **WekB**

```
NCTC13441       MKVAFLSAYDPLSTSSWSGTPYYMLKALSKRNISIEILGPVNSYMIYMLKVYKLILRCFG  60
VREC1403        MKVAFLSAYDPLSTSSWSGTPYYMLKALSKRNISIEILGPVNSYMIYMLKVYKLILRCFG  60
ECO0218         MKVAFLSAYDPLSTSSWSGTPYYMLKALSKRNISIEILGPVNSYMIYMLKVYKLILRCFG  60
ECO0061         MKVAFLSAYDPLSTSSWSGTPYYMLKALSKRNISIEILGPVNSYMIYMLKVYKLILRCFG  60
ECO0211         MKVAFLSAYDPLSTSSWSGTPYYMLKALSKRNISIEILGPVNSYMIYMLKVYKLILRCFG  60
ECO0387         MKVAFLSAYDPLSTSSWSGTPYYMLKALSKRNISIEILGPVNSYMIYMLKVYKLILRCFG  60
VRES1610        MKVAFLSAYDPLSTSSWSGTPYYMLKALSKRNISIEILGPVNSYMIYMLKVYKLILRCFG  60
VRES1619        MKVAFLSAYDPLSTSSWSGTPYYMLKALSKRNISIEILGPVNSYMIYMLKVYKLILRCFG  60
VRES0739        MKVAFLSAYDPLSTSSWSGTPYYMLKALSKRNISIEILGPVNSYMIYMLKVYKLILRCFG  60
VREC0645        MKVAFLSAYDPLSTSSWSGTPYYMLKALSKRNISIEILGPVNSYMIYMLKVYKLILRCFG  60
VRES1160        MKVAFLSAYDPLSTSSWSGTPYYMLKALSKRNISIEILGPVNSYMIYMLKVYKLILRCFG  60
VRES0700        MKVAFLSAYDPLSTSSWSGTPYYMLKALSKRNISIEILGPVNSYMIYMLKVYKLILRCFG  60
VREC1630        MKVAFLSAYDPLSTSSWSGTPYYMLKALSKRNISIEILGPVNSYMIYMLKVYKLILRCFG  60
VREC1428        MKVAFLSAYDPLSTSSWSGTPYYMLKALSKRNISIEILGPVNSYMIYMLKVYKLILRCFG  60
VREC1106        MKVAFLSAYDPLSTSSWSGTPYYMLKALSKRNISIEILGPVNSYMIYMLKVYKLILRCFG  60
VREC1101        MKVAFLSAYDPLSTSSWSGTPYYMLKALSKRNISIEILGPVNSYMIYMLKVYKLILRCFG  60
VREC1073        MKVAFLSAYDPLSTSSWSGTPYYMLKALSKRNISIEILGPVNSYMIYMLKVYKLILRCFG  60
VREC1013        MKVAFLSAYDPLSTSSWSGTPYYMLKALSKRNISIEILGPVNSYMIYMLKVYKLILRCFG  60
VREC0926        MKVAFLSAYDPLSTSSWSGTPYYMLKALSKRNISIEILGPVNSYMIYMLKVYKLILRCFG  60
VREC0865        MKVAFLSAYDPLSTSSWSGTPYYMLKALSKRNISIEILGPVNSYMIYMLKVYKLILRCFG  60
VREC0829        MKVAFLSAYDPLSTSSWSGTPYYMLKALSKRNISIEILGPVNSYMIYMLKVYKLILRCFG  60
VREC0693        MKVAFLSAYDPLSTSSWSGTPYYMLKALSKRNISIEILGPVNSYMIYMLKVYKLILRCFG  60
ECO0433         MKVAFLSAYDPLSTSSWSGTPYYMLKALSKRNISIEILGPVNSYMIYMLKVYKLILRCFG  60
ECO0224         MKVAFLSAYDPLSTSSWSGTPYYMLKALSKRNISIEILGPVNSYMIYMLKVYKLILRCFG  60
ECO0411         MKVAFLSAYDPLSTSSWSGTPYYMLKALSKRNISIEILGPVNSYMIYMLKVYKLILRCFG  60
ECO0431         MKVAFLSAYDPLSTSSWSGTPYYMLKALSKRNISIEILGPVNSYMIYMLKVYKLILRCFG  60
ECO0353         MKVAFLSAYDPLSTSSWSGTPYYMLKALSKRNISIEILGPVNSYMIYMLKVYKLILRCFG  60
                ************************************************************

NCTC13441       KEYDYSHSKLLSRYYGRIFGRKLKKIDGLDFIIAPAGSSQIAFLKTTIPIIYLSDTTYDQ  120
VREC1403        KEYDYSHSKLLSRYYGRIFGRKLKKIDGLDFIIAPAGSSQIAFLKTTIPIIYLSDTTYDQ  120
ECO0218         KEYDYSHSKLLSRYYGRIFGRKLKKIDGLDFIIAPAGSSQIAFLKTTIPIIYLSDTTYDQ  120
ECO0061         KEYDYSHSKLLSRYYGRIFGRKLKKIDGLDFIIAPAGSSQIAFLKTTIPIIYLSDTTYDQ  120
ECO0211         KEYDYSHSKLLSRYYGRIFGRKLKKIDGLDFIIAPAGSSQIAFLKTTIPIIYLSDTTYDQ  120
ECO0387         KEYDYSHSKLLSRYYGRIFGRKLKKIDGLDFIIAPAGSSQIAFLKTTIPIIYLSDTTYDQ  120
VRES1610        KEYDYSHSKLLSRYYGRIFGRKLKKIDGLDFIIAPAGSSQIAFLKTTIPIIYLSDTTYDQ  120
VRES1619        KEYDYSHSKLLSRYYGRIFGRKLKKIDGLDFIIAPAGSSQIAFLKTTIPIIYLSDTTYDQ  120
VRES0739        KEYDYSHSKLLSRYYGRIFGRKLKKIDGLDFIIAPAGSSQIAFLKTTIPIIYLSDTTYDQ  120
VREC0645        KEYDYSHSKLLSRYYGRIFGRKLKKIDGLDFIIAPAGSSQIAFLKTTIPIIYLSDTTYDQ  120
VRES1160        KEYDYSHSKLLSRYYGRIFGRKLKKIDGLDFIIAPAGSSQIAFLKTTIPIIYLSDTTYDQ  120
VRES0700        KEYDYSHSKLLSRYYGRIFGRKLKKIDGLDFIIAPAGSSQIAFLKTTIPIIYLSDTTYDQ  120
VREC1630        KEYDYSHSKLLSRYYGRIFGRKLKKIDGLDFIIAPAGSSQIAFLKTTIPIIYLSDTTYDQ  120
VREC1428        KEYDYSHSKLLSRYYGRIFGRKLKKIDGLDFIIAPAGSSQIAFLKTTIPIIYLSDTTYDQ  120
VREC1106        KEYDYSHSKLLSRYYGRIFGRKLKKIDGLDFIIAPAGSSQIAFLKTTIPIIYLSDTTYDQ  120
VREC1101        KEYDYSHSKLLSRYYGRIFGRKLKKIDGLDFIIAPAGSSQIAFLKTTIPIIYLSDTTYDQ  120
VREC1073        KEYDYSHSKLLSRYYGRIFGRKLKKIDGLDFIIAPAGSSQIAFLKTTIPIIYLSDTTYDQ  120
VREC1013        KEYDYSHSKLLSRYYGRIFGRKLKKIDGLDFIIAPAGSSQIAFLKTTIPIIYLSDTTYDQ  120
VREC0926        KEYDYSHSKLLSRYYGRIFGRKLKKIDGLDFIIAPAGSSQIAFLKTTIPIIYLSDTTYDQ  120
VREC0865        KEYDYSHSKLLSRYYGRIFGRKLKKIDGLDFIIAPAGSSQIAFLKTTIPIIYLSDTTYDQ  120
VREC0829        KEYDYSHSKLLSRYYGRIFGRKLKKIDGLDFIIAPAGSSQIAFLKTTIPIIYLSDTTYDQ  120
VREC0693        KEYDYSHSKLLSRYYGRIFGRKLKKIDGLDFIIAPAGSSQIAFLKTTIPIIYLSDTTYDQ  120
ECO0433         KEYDYSHSKLLSRYYGRIFGRKLKKIDGLDFIIAPAGSSQIAFLKTTIPIIYLSDTTYDQ  120
ECO0224         KEYDYSHSKLLSRYYGRIFGRKLKKIDGLDFIIAPAGSSQIAFLKTTIPIIYLSDTTYDQ  120
ECO0411         KEYDYSHSKLLSRYYGRIFGRKLKKIDGLDFIIAPAGSSQIAFLKTTIPIIYLSDTTYDQ  120
ECO0431         KEYDYSHSKLLSRYYGRIFGRKLKKIDGLDFIIAPAGSSQIAFLKTTIPIIYLSDTTYDQ  120
ECO0353         KEYDYSHSKLLSRYYGRIFGRKLKKIDGLDFIIAPAGSSQIAFLKTTIPIIYLSDTTYDQ  120
                ************************************************************

NCTC13441       LKSYYPNLNKKTIINDEDASLIERKAIEKATVVSFPSKWAMDFCRNYYRLDFDKLVEIPW  180
VREC1403        LKSYYPNLNKKTIINDEDASLIERKAIEKATVVSFPSKWAMDFCRNYYRLDFDKLVEIPW  180
ECO0218         LKSYYPNLNKKTIINDEDASLIERKAIEKATVVSFPSKWAMDFCRNYYRLDFDKLVEIPW  180
ECO0061         LKSYYPNLNKKTIINDEDASLIERKAIEKATVVSFPSKWAMDFCRNYYRLDFDKLVEIPW  180
ECO0211         LKSYYPNLNKKTIINDEDASLIERKAIEKATVVSFPSKWAMDFCRNYYRLDFDKLVEIPW  180
ECO0387         LKSYYPNLNKKTIINDEDASLIERKAIEKATVVSFPSKWAMDFCRNYYRLDFDKLVEIPW  180
VRES1610        LKSYYPNLNKKTIINDEDASLIERKAIEKATVVSFPSKWAMDFCRNYYRLDFDKLVEIPW  180
VRES1619        LKSYYPNLNKKTIINDEDASLIERKAIEKATVVSFPSKWAMDFCRNYYRLDFDKLVEIPW  180
VRES0739        LKSYYPNLNKKTIINDEDASLIERKAIEKATVVSFPSKWAMDFCRNYYRLDFDKLVEIPW  180
VREC0645        LKSYYPNLNKKTIINDEDASLIERKAIEKATVVSFPSKWAMDFCRNYYRLDFDKLVEIPW  180
VRES1160        LKSYYPNLNKKTIINDEDASLIERKAIEKATVVSFPSKWAMDFCRNYYRLDFDKLVEIPW  180
VRES0700        LKSYYPNLNKKTIINDEDASLIERKAIEKATVVSFPSKWAMDFCRNYYRLDFDKLVEIPW  180
VREC1630        LKSYYPNLNKKTIINDEDASLIERKAIEKATVVSFPSKWAMDFCRNYYRLDFDKLVEIPW  180
VREC1428        LKSYYPNLNKKTIINDEDASLIERKAIEKATVVSFPSKWAMDFCRNYYRLDFDKLVEIPW  180
VREC1106        LKSYYPNLNKKTIINDEDASLIERKAIEKATVVSFPSKWAMDFCRNYYRLDFDKLVEIPW  180
```

```
VREC1101       LKSYYPNLNKKTIINDEDASLIERKAIEKATVVSFPSKWAMDFCRNYYRLDFDKLVEIPW 180
VREC1073       LKSYYPNLNKKTIINDEDASLIERKAIEKATVVSFPSKWAMDFCRNYYRLDFDKLVEIPW 180
VREC1013       LKSYYPNLNKKTIINDEDASLIERKAIEKATVVSFPSKWAMDFCRNYYRLDFDKLVEIPW 180
VREC0926       LKSYYPNLNKKTIINDEDASLIERKAIEKATVVSFPSKWAMDFCRNYYRLDFDKLVEIPW 180
VREC0865       LKSYYPNLNKKTIINDEDASLIERKAIEKATVVSFPSKWAMDFCRNYYRLDFDKLVEIPW 180
VREC0829       LKSYYPNLNKKTIINDEDASLIERKAIEKATVVSFPSKWAMDFCRNYYRLDFDKLVEIPW 180
VREC0693       LKSYYPNLNKKTIINDEDASLIERKAIEKATVVSFPSKWAMDFCRNYYRLDFDKLVEIPW 180
ECO0433        LKSYYPNLNKKTIINDEDASLIERKAIEKATVVSFPSKWAMDFCRNYYRLDFDKLVEIPW 180
ECO0224        LKSYYPNLNKKTIINDEDASLIERKAIEKATVVSFPSKWAMDFCRNYYRLDFDKLVEIPW 180
ECO0411        LKSYYPNLNKKTIINDEDASLIERKAIEKATVVSFPSKWAMDFCRNYYRLDFDKLVEIPW 180
ECO0431        LKSYYPNLNKKTIINDEDASLIERKAIEKATVVSFPSKWAMDFCRNYYRLDFDKLVEIPW 180
ECO0353        LKSYYPNLNKKTIINDEDASLIERKAIEKATVVSFPSKWAMDFCRNYYRLDFDKLVEIPW 180
               ************************************************************

NCTC13441      GANLFDDIHFANKNIIQKNSYTCLFLGVDWERKGGKTALKAIEYVRQLYGIDVRLKICGC 240
VREC1403       GANLFDDIHFANKNIIQKNSYTCLFLGVDWERKGGKTALKAIEYVRQLYGIDVRLKICGC 240
ECO0218        GANLFDDIHFANKNIIQKNSYTCLFLGVDWERKGGKTALKAIEYVRQLYGIDVRLKICGC 240
ECO0061        GANLFDDIHFANKNIIQKNSYTCLFLGVDWERKGGKTALKAIEYVRQLYGIDVRLKICGC 240
ECO0211        GANLFDDIHFANKNIIQKNSYTCLFLGVDWERKGGKTALKAIEYVRQLYGIDVRLKICGC 240
ECO0387        GANLFDDIHFANKNIIQKNSYTCLFLGVDWERKGGKTALKAIEYVRQLYGIDVRLKICGC 240
VRES1610       GANLFDDIHFANKNIIQKNSYTCLFLGVDWERKGGKTALKAIEYVRQLYGIDVRLKICGC 240
VRES1619       GANLFDDIHFANKNIIQKNSYTCLFLGVDWERKGGKTALKAIEYVRQLYGIDVRLKICGC 240
VRES0739       GANLFDDIHFANKNIIQKNSYTCLFLGVDWERKGGKTALKAIEYVRQLYGIDVRLKICGC 240
VREC0645       GANLFDDIHFANKNIIQKNSYTCLFLGVDWERKGGKTALKAIEYVRQLYGIDVRLKICGC 240
VRES1160       GANLFDDIHFANKNIIQKNSYTCLFLGVDWERKGGKTALKAIEYVRQLYGIDVRLKICGC 240
VRES0700       GANLFDDIHFANKNIIQKNSYTCLFLGVDWERKGGKTALKAIEYVRQLYGIDVRLKICGC 240
VREC1630       GANLFDDIHFANKNIIQKNSYTCLFLGVDWERKGGKTALKAIEYVRQLYGIDVRLKICGC 240
VREC1428       GANLFDDIHFANKNIIQKNSYTCLFLGVDWERKGGKTALKAIEYVRQLYGIDVRLKICGC 240
VREC1106       GANLFDDIHFANKNIIQKNSYTCLFLGVDWERKGGKTALKAIEYVRQLYGIDVRLKICGC 240
VREC1101       GANLFDDIHFANKNIIQKNSYTCLFLGVDWERKGGKTALKAIEYVRQLYGIDVRLKICGC 240
VREC1073       GANLFDDIHFANKNIIQKNSYTCLFLGVDWERKGGKTALKAIEYVRQLYGIDVRLKICGC 240
VREC1013       GANLFDDIHFANKNIIQKNSYTCLFLGVDWERKGGKTALKAIEYVRQLYGIDVRLKICGC 240
VREC0926       GANLFDDIHFANKNIIQKNSYTCLFLGVDWERKGGKTALKAIEYVRQLYGIDVRLKICGC 240
VREC0865       GANLFDDIHFANKNIIQKNSYTCLFLGVDWERKGGKTALKAIEYVRQLYGIDVRLKICGC 240
VREC0829       GANLFDDIHFANKNIIQKNSYTCLFLGVDWERKGGKTALKAIEYVRQLYGIDVRLKICGC 240
VREC0693       GANLFDDIHFANKNIIQKNSYTCLFLGVDWERKGGKTALKAIEYVRQLYGIDVRLKICGC 240
ECO0433        GANLFDDIHFANKNIIQKNSYTCLFLGVDWERKGGKTALKAIEYVRQLYGIDVRLKICGC 240
ECO0224        GANLFDDIHFANKNIIQKNSYTCLFLGVDWERKGGKTALKAIEYVRQLYGIDVRLKICGC 240
ECO0411        GANLFDDIHFANKNIIQKNSYTCLFLGVDWERKGGKTALKAIEYVRQLYGIDVRLKICGC 240
ECO0431        GANLFDDIHFANKNIIQKNSYTCLFLGVDWERKGGKTALKAIEYVRQLYGIDVRLKICGC 240
ECO0353        GANLFDDIHFANKNIIQKNSYTCLFLGVDWERKGGKTALKAIEYVRQLYGIDVRLKICGC 240
               ************************************************************

NCTC13441      TPNQKILPTWVELIDKVDKNNVDEYQKFIDVLSNADILLLPTIAECYGMVFCEAAAFGLP 300
VREC1403       TPNQKILPTWVELIDKVDKNNVDEYQKFIDVLSNADILLLPTIAECYGMVFCEAAAFGLP 300
ECO0218        TPNQKILPTWVELIDKVDKNNVDEYQKFIDVLSNADILLLPTIAECYGMVFCEAAAFGLP 300
ECO0061        TPNQKILPTWVELIDKVDKNNVDEYQKFIDVLSNADILLLPTIAECYGMVFCEAAAFGLP 300
ECO0211        TPNQKILPTWVELIDKVDKNNVDEYQKFIDVLSNADILLLPTIAECYGMVFCEAAAFGLP 300
ECO0387        TPNQKILPTWVELIDKVDKNNVDEYQKFIDVLSNADILLLPTIAECYGMVFCEAAAFGLP 300
VRES1610       TPNQKILPTWVELIDKVDKNNVDEYQKFIDVLSNADILLLPTIAECYGMVFCEAAAFGLP 300
VRES1619       TPNQKILPTWVELIDKVDKNNVDEYQKFIDVLSNADILLLPTIAECYGMVFCEAAAFGLP 300
VRES0739       TPNQKILPTWVELIDKVDKNNVDEYQKFIDVLSNADILLLPTIAECYGMVFCEAAAFGLP 300
VREC0645       TPNQKILPTWVELIDKVDKNNVDEYQKFIDVLSNADILLLPTIAECYGMVFCEAAAFGLP 300
VRES1160       TPNQKILPTWVELIDKVDKNNVDEYQKFIDVLSNADILLLPTIAECYGMVFCEAAAFGLP 300
VRES0700       TPNQKILPTWVELIDKVDKNNVDEYQKFIDVLSNADILLLPTIAECYGMVFCEAAAFGLP 300
VREC1630       TPNQKILPTWVELIDKVDKNNVDEYQKFIDVLSNADILLLPTIAECYGMVFCEAAAFGLP 300
VREC1428       TPNQKILPTWVELIDKVDKNNVDEYQKFIDVLSNADILLLPTIAECYGMVFCEAAAFGLP 300
VREC1106       TPNQKILPTWVELIDKVDKNNVDEYQKFIDVLSNADILLLPTIAECYGMVFCEAAAFGLP 300
VREC1101       TPNQKILPTWVELIDKVDKNNVDEYQKFIDVLSNADILLLPTIAECYGMVFCEAAAFGLP 300
VREC1073       TPNQKILPTWVELIDKVDKNNVDEYQKFIDVLSNADILLLPTIAECYGMVFCEAAAFGLP 300
VREC1013       TPNQKILPTWVELIDKVDKNNVDEYQKFIDVLSNADILLLPTIAECYGMVFCEAAAFGLP 300
VREC0926       TPNQKILPTWVELIDKVDKNNVDEYQKFIDVLSNADILLLPTIAECYGMVFCEAAAFGLP 300
VREC0865       TPNQKILPTWVELIDKVDKNNVDEYQKFIDVLSNADILLLPTIAECYGMVFCEAAAFGLP 300
VREC0829       TPNQKILPTWVELIDKVDKNNVDEYQKFIDVLSNADILLLPTIAECYGMVFCEAAAFGLP 300
VREC0693       TPNQKILPTWVELIDKVDKNNVDEYQKFIDVLSNADILLLPTIAECYGMVFCEAAAFGLP 300
ECO0433        TPNQKILPTWVELIDKVDKNNVDEYQKFIDVLSNADILLLPTIAECYGMVFCEAAAFGLP 300
ECO0224        TPNQKILPTWVELIDKVDKNNVDEYQKFIDVLSNADILLLPTIAECYGMVFCEAAAFGLP 300
ECO0411        TPNQKILPTWVELIDKVDKNNVDEYQKFIDVLSNADILLLPTIAECYGMVFCEAAAFGLP 300
ECO0431        TPNQKILPTWVELIDKVDKNNVDEYQKFIDVLSNADILLLPTIAECYGMVFCEAAAFGLP 300
ECO0353        TPNQKILPTWVELIDKVDKNNVDEYQKFIDVLSNADILLLPTIAECYGMVFCEAAAFGLP 300
               ************************************************************

NCTC13441      VVATDTGGVSSIVINERTGILIKDPLDYKHFGNAIHKIISSVETYQNYSQNARIRYNNIL 360
VREC1403       VVATDTGGVSSIVINERTGILIKDPLDYKHFGNAIHKIISSVETYQNYSQNARIRYNNIL 360
ECO0218        VVATDTGGVSSIVINERTGILIKDPLDYKHFGNAIHKIISSVETYQNYSQNARIRYNNIL 360
ECO0061        VVATDTGGVSSIVINERTGILIKDPLDYKHFGNAIHKIISSVETYQNYSQNARIRYNNIL 360
ECO0211        VVATDTGGVSSIVINERTGILIKDPLDYKHFGNAIHKIISSVETYQNYSQNARIRYNNIL 360
```

```
ECO0387        VVATDTGGVSSIVINERTGILIKDPLDYKHFGNAIHKIISSVETYQNYSQNARIRYNNIL 360
VRES1610       VVATDTGGVSSIVINERTGILIKDPLDYKHFGNAIHKIISSVETYQNYSQNARIRYNNIL 360
VRES1619       VVATDTGGVSSIVINERTGILIKDPLDYKHFGNAIHKIISSVETYQNYSQNARIRYNNIL 360
VRES0739       VVATDTGGVSSIVINERTGILIKDPLDYKHFGNAIHKIISSVETYQNYSQNARIRYNNIL 360
VREC0645       VVATDTGGVSSIVINERTGILIKDPLDYKHFGNAIHKIISSVETYQN------------- 347
VRES1160       VVATDTGGVSSIVINERTGILIKDPLDYKHFGNAIHKIISSVETYQNYSQNARIRYNNIL 360
VRES0700       VVATDTGGVSSIVINERTGILIKDPLDYKHFGNAIHKIISSVETYQNYSQNARIRYNNIL 360
VREC1630       VVATDTGGVSSIVINERTGILIKDPLDYKHFGNAIHKIISSVETYQNYSQNARIRYNNIL 360
VREC1428       VVATDTGGVSSIVINERTGILIKDPLDYKHFGNAIHKIISSVETYQNYSQNARIRYNNIL 360
VREC1106       VVATDTGGVSSIVINERTGILIKDPLDYKHFGNAIHKIISSVETYQNYSQNARIRYNNIL 360
VREC1101       VVATDTGGVSSIVINERTGILIKDPLDYKHFGNAIHKIISSVETYQNYSQNARIRYNNIL 360
VREC1073       VVATDTGGVSSIVINERTGILIKDPLDYKHFGNAIHKIISSVETYQNYSQNARIRYNNIL 360
VREC1013       VVATDTGGVSSIVINERTGILIKDPLDYKHFGNAIHKIISSVETYQNYSQNARIRYNNIL 360
VREC0926       VVATDTGGVSSIVINERTGILIKDPLDYKHFGNAIHKIISSVETYQNYSQNARIRYNNIL 360
VREC0865       VVATDTGGVSSIVINERTGILIKDPLDYKHFGNAIHKIISSVETYQNYSQNARIRYNNIL 360
VREC0829       VVATDTGGVSSIVINERTGILIKDPLDYKHFGNAIHKIISSVETYQNYSQNARIRYNNIL 360
VREC0693       VVATDTGGVSSIVINERTGILIKDPLDYKHFGNAIHKIISSVETYQNYSQNARIRYNNIL 360
ECO0433        VVATDTGGVSSIVINERTGILIKDPLDYKHFGNAIHKIISSVETYQNYSQNARIRYNNIL 360
ECO0224        VVATDTGGVSSIVINERTGILIKDPLDYKHFGNAIHKIISSVETYQNYSQNARIRYNNIL 360
ECO0411        VVATDTGGVSSIVINERTGILIKDPLDYKHFGNAIHKIISSVETYQNYSQNARIRYNNIL 360
ECO0431        VVATDTGGVSSIVINERTGILIKDPLDYKHFGNAIHKIISSVETYQNYSQNARIRYNNIL 360
ECO0353        VVATDTGGVSSIVINERTGILIKDPLDYKHFGNAIHKIISSVETYQNYSQNARIRYNNIL 360
               ***********************************************

NCTC13441      HWDNWAKKIIEIMYEHKNRRIK 382
VREC1403       HWDNWAKKIIEIMYEHKNRRIK 382
ECO0218        HWDNWAKKIIEIMYEHKNRRIK 382
ECO0061        HWDNWAKKIIEIMYEHKNRRIK 382
ECO0211        HWDNWAKKIIEIMYEHKNRRIK 382
ECO0387        HWDNWAKKIIEIMYEHKNRRIK 382
VRES1610       HWDNWAKKIIEIMYEHKNRRIK 382
VRES1619       HWDNWAKKIIEIMYEHKNRRIK 382
VRES0739       HWDNWAKKIIEIMYEHKNRRIK 382
VREC0645       ---------------------- 347
VRES1160       HWDNWAKKIIEIMYEHKNRRIK 382
VRES0700       HWDNWAKKIIEIMYEHKNRRIK 382
VREC1630       HWDNWAKKIIEIMYEHKNRRIK 382
VREC1428       HWDNWAKKIIEIMYEHKNRRIK 382
VREC1106       HWDNWAKKIIEIMYEHKNRRIK 382
VREC1101       HWDNWAKKIIEIMYEHKNRRIK 382
VREC1073       HWDNWAKKIIEIMYEHKNRRIK 382
VREC1013       HWDNWAKKIIEIMYEHKNRRIK 382
VREC0926       HWDNWAKKIIEIMYEHKNRRIK 382
VREC0865       HWDNWAKKIIEIMYEHKNRRIK 382
VREC0829       HWDNWAKKIIEIMYEHKNRRIK 382
VREC0693       HWDNWAKKIIEIMYEHKNRRIK 382
ECO0433        HWDNWAKKIIEIMYEHKNRRIK 382
ECO0224        HWDNWAKKIIEIMYEHKNRRIK 382
ECO0411        HWDNWAKKIIEIMYEHKNRRIK 382
ECO0431        HWDNWAKKIIEIMYEHKNRRIK 382
ECO0353        HWDNWAKKIIEIMYEHKNRRIK 382
```

CLUSTAL O(1.2.4) multiple sequence alignment - **Wzy**

```
ECO0353         MSIRIEESNSTKRIICLFILFLVFPDFLFYTLGVDNFSISTIISITLLFVFLRAKNICKD        60
ECO0431         MSIRIEESNSTKRIICLFILFLVFPDFLFYTLGVDNFSISTIISITLLFVFLRAKNICKD        60
ECO0411         MSIRIEESNSTKRIICLFILFLVFPDFLFYTLGVDNFSISTIISITLLFVFLRAKNICKD        60
ECO0224         MSIRIEESNSTKRIICLFILFLVFPDFLFYTLGVDNFSISTIISITLLFVFLRAKNICKD        60
ECO0433         MSIRIEESNSTKRIICLFILFLVFPDFLFYTLGVDNFSISTIISITLLFVFLRAKNICKD        60
VREC0693        MSIRIEESNSTKRIICLFILFLVFPDFLFYTLGVDNFSISTIISITLLFVFLRAKNICKD        60
VREC0829        MSIRIEESNSTKRIICLFILFLVFPDFLFYTLGVDNFSISTIISITLLFVFLRAKNICKD        60
VREC0865        MSIRIEESNSTKRIICLFILFLVFPDFLFYTLGVDNFSISTIISITLLFVFLRAKNICKD        60
VREC0926        MSIRIEESNSTKRIICLFILFLVFPDFLFYTLGVDNFSISTIISITLLFVFLRAKNICKD        60
VREC1013        MSIRIEESNSTKRIICLFILFLVFPDFLFYTLGVDNFSISTIISITLLFVFLRAKNICKD        60
VREC1073        MSIRIEESNSTKRIICLFILFLVFPDFLFYTLGVDNFSISTIISITLLFVFLRAKNICKD        60
VREC1101        MSIRIEESNSTKRIICLFILFLVFPDFLFYTLGVDNFSISTIISITLLFVFLRAKNICKD        60
VREC1106        MSIRIEESNSTKRIICLFILFLVFPDFLFYTLGVDNFSISTIISITLLFVFLRAKNICKD        60
VREC1428        MSIRIEESNSTKRIICLFILFLVFPDFLFYTLGVDNFSISTIISITLLFVFLRAKNICKD        60
VREC1630        MSIRIEESNSTKRIICLFILFLVFPDFLFYTLGVDNFSISTIISITLLFVFLRAKNICKD        60
VRES0700        MSIRIEESNSTKRIICLFILFLVFPDFLFYTLGVDNFSISTIISITLLFVFLRAKNICKD        60
VRES1160        MSIRIEESNSTKRIICLFILFLVFPDFLFYTLGVDNFSISTIISITLLFVFLRAKNICKD        60
VREC0645        MSIRIEESNSTKRIICLFILFLVFPDFLFYTLGVDNFSISTIISITLLFVFLRAKNICKD        60
VRES0739        MSIRIEESNSTKRIICLFILFLVFPDFLFYTLGVDNFSISTIISITLLFVFLRAKNICKD        60
VRES1619        MSIRIEESNSTKRIICLFILFLVFPDFLFYTLGVDNFSISTIISITLLFVFLRAKNICKD        60
VRES1610        MSIRIEESNSTKRIICLFILFLVFPDFLFYTLGVDNFSISTIISITLLFVFLRAKNICKD        60
ECO0211         MSIRIEESNSTKRIICLFILFLVFPDFLFYTLGVDNFSISTIISITLLFVFLRAKNICKD        60
ECO0061         MSIRIEESNSTKRIICLFILFLVFPDFLFYTLGVDNFSISTIISITLLFVFLRAKNICKD        60
VREC1403        MSIRIEESNSTKRIICLFILFLVFPDFLFYTLGVDNFSISTIISITLLFVFLRAKNICKD        60
NCTC13441       MSIRIEESNSTKRIICLFILFLVFPDFLFYTLGVDNFSISTIISITLLFVFLRAKNICKD        60
ECO0218orf2     ------------------------------------------------------------        0
ECO0387orf2     ------------------------------------------------------------        0
ECO0218orf1     MSIRIEESNSTKRIICLFILFLVFPDFLFYTLGVDNFSISTIISITLLFVFLRAKNICKD        60
ECO0387orf1     MSIRIEESNSTKRIICLFILFLVFPDFLFYTLGVDNFSISTIISITLLFVFLRAKNICKD        60

ECO0353         NFLIIVALFILLCFNCLLSMLFNIEQALTFKVVLSIYSILIMAYVSSCYAQTLWLCSEEI        120
ECO0431         NFLIIVALFILLCFNCLLSMLFNIEQALTFKVVLSIYSILIMAYVSSCYAQTLWLCSEEI        120
ECO0411         NFLIIVALFILLCFNCLLSMLFNIEQALTFKVVLSIYSILIMAYVSSCYAQTLWLCSEEI        120
ECO0224         NFLIIVALFILLCFNCLLSMLFNIEQALTFKVVLSIYSILIMAYVSSCYAQTLWLCSEEI        120
ECO0433         NFLIIVALFILLCFNCLLSMLFNIEQALTFKVVLSIYSILIMAYVSSCYAQTLWLCSEEI        120
VREC0693        NFLIIVALFILLCFNCLLSMLFNIEQALTFKVVLSIYSILIMAYVSSCYAQTLWLCSEEI        120
VREC0829        NFLIIVALFILLCFNCLLSMLFNIEQALTFKVVLSIYSILIMAYVSSCYAQTLWLCSEEI        120
VREC0865        NFLIIVALFILLCFNCLLSMLFNIEQALTFKVVLSIYSILIMAYVSSCYAQTLWLCSEEI        120
VREC0926        NFLIIVALFILLCFNCLLSMLFNIEQALTFKVVLSIYSILIMAYVSSCYAQTLWLCSEEI        120
VREC1013        NFLIIVALFILLCFNCLLSMLFNIEQALTFKVVLSIYSILIMAYVSSCYAQTLWLCSEEI        120
VREC1073        NFLIIVALFILLCFNCLLSMLFNIEQALTFKVVLSIYSILIMAYVSSCYAQTLWLCSEEI        120
VREC1101        NFLIIVALFILLCFNCLLSMLFNIEQALTFKVVLSIYSILIMAYVSSCYAQTLWLCSEEI        120
VREC1106        NFLIIVALFILLCFNCLLSMLFNIEQALTFKVVLSIYSILIMAYVSSCYAQTLWLCSEEI        120
VREC1428        NFLIIVALFILLCFNCLLSMLFNIEQALTFKVVLSIYSILIMAYVSSCYAQTLWLCSEEI        120
VREC1630        NFLIIVALFILLCFNCLLSMLFNIEQALTFKVVLSIYSILIMAYVSSCYAQTLWLCSEEI        120
VRES0700        NFLIIVALFILLCFNCLLSMLFNIEQALTFKVVLSIYSILIMAYVSSCYAQTLWLCSEEI        120
VRES1160        NFLIIVALFILLCFNCLLSMLFNIEQALTFKVVLSIYSILIMAYVSSCYAQTLWLCSEEI        120
VREC0645        NFLIIVALFILLCFNCLLSMLFNIEQALTFKVVLSIYSILIMAYVSSCYAQTLWLCSEEI        120
VRES0739        NFLIIVALFILLCFNCLLSMLFNIEQALTFKVVLSIYSILIMAYVSSCYAQTLWLCSEEI        120
VRES1619        NFLIIVALFILLCFNCLLSMLFNIEQALTFKVVLSIYSILIMAYVSSCYAQTLWLCSEEI        120
VRES1610        NFLIIVALFILLCFNCLLSMLFNIEQALTFKVVLSIYSILIMAYVSSCYAQTLWLCSEEI        120
ECO0211         NFLIIVALFILLCFNCLLSMLFNIEQALTFKVVLSIYSILIMAYVSSCYAQTLWLCSEEI        120
ECO0061         NFLIIVALFILLCFNCLLSMLFNIEQALTFKVVLSIYSILIMAYVSSCYAQTLWLCSEEI        120
VREC1403        NFLIIVALFILLCFNCLLSMLFNIEQALTFKVVLSIYSILIMAYVSSCYAQTLWLCSEEI        120
NCTC13441       NFLIIVALFILLCFNCLLSMLFNIEQALTFKVVLSIYSILIMAYVSSCYAQTLWLCSEEI        120
ECO0218orf2     ------------------------------------------------------------        0
ECO0387orf2     ------------------------------------------------------------        0
ECO0218orf1     NFLIIVALFILLCFNCLLSMLFNIEQALTFKVVLSIYSILIMAYVSSCYAQTLWLCSEEI        120
ECO0387orf1     NFLIIVALFILLCFNCLLSMLFNIEQALTFKVVLSIYSILIMAYVSSCYAQTLWLCSEEI        120

ECO0353         LKRSVFYLFAFLCLIGIISILLQKTEIIHDKSMILFPEPSAFALVFIPIFSFCLYYTRGG        180
ECO0431         LKRSVFYLFAFLCLIGIISILLQKTEIIHDKSMILFPEPSAFALVFIPIFSFCLYYTRGG        180
ECO0411         LKRSVFYLFAFLCLIGIISILLQKTEIIHDKSMILFPEPSAFALVFIPIFSFCLYYTRGG        180
ECO0224         LKRSVFYLFAFLCLIGIISILLQKTEIIHDKSMILFPEPSAFALVFIPIFSFCLYYTRGG        180
ECO0433         LKRSVFYLFAFLCLIGIISILLQKTEIIHDKSMILFPEPSAFALVFIPIFSFCLYYTRGG        180
VREC0693        LKRSVFYLFAFLCLIGIISILLQKTEIIHDKSMILFPEPSAFALVFIPIFSFCLYYTRGG        180
VREC0829        LKRSVFYLFAFLCLIGIISILLQKTEIIHDKSMILFPEPSAFALVFIPIFSFCLYYTRGG        180
VREC0865        LKRSVFYLFAFLCLIGIISILLQKTEIIHDKSMILFPEPSAFALVFIPIFSFCLYYTRGG        180
VREC0926        LKRSVFYLFAFLCLIGIISILLQKTEIIHDKSMILFPEPSAFALVFIPIFSFCLYYTRGG        180
VREC1013        LKRSVFYLFAFLCLIGIISILLQKTEIIHDKSMILFPEPSAFALVFIPIFSFCLYYTRGG        180
VREC1073        LKRSVFYLFAFLCLIGIISILLQKTEIIHDKSMILFPEPSAFALVFIPIFSFCLYYTRGG        180
```

```
VREC1101        LKRSVFYLFAFLCLIGIISILLQKTEIIHDKSMILFPEPSAFALVFIPIFSFCLYYTRGG    180
VREC1106        LKRSVFYLFAFLCLIGIISILLQKTEIIHDKSMILFPEPSAFALVFIPIFSFCLYYTRGG    180
VREC1428        LKRSVFYLFAFLCLIGIISILLQKTEIIHDKSMILFPEPSAFALVFIPIFSFCLYYTRGG    180
VREC1630        LKRSVFYLFAFLCLIGIISILLQKTEIIHDKSMILFPEPSAFALVFIPIFSFCLYYTRGG    180
VRES0700        LKRSVFYLFAFLCLIGIISILLQKTEIIHDKSMILFPEPSAFALVFIPIFSFCLYYTRGG    180
VRES1160        LKRSVFYLFAFLCLIGIISILLQKTEIIHDKSMILFPEPSAFALVFIPIFSFCLYYTRGG    180
VREC0645        LKRSVFYLFAFLCLIGIISILLQKTEIIHDKSMILFPEPSAFALVFIPIFSFCLYYTRGG    180
VRES0739        LKRSVFYLFAFLCLIGIISILLQKTEIIHDKSMILFPEPSAFALVFIPIFSFCLYYTRGG    180
VRES1619        LKRSVFYLFAFLCLIGIISILLQKTEIIHDKSMILFPEPSAFALVFIPIFSFCLYYTRGG    180
VRES1610        LKRSVFYLFAFLCLIGIISILLQKTEIIHDKSMILFPEPSAFALVFIPIFSFCLYYTRGG    180
ECO0211         LKRSVFYLFAFLCLIGIISILLQKTEIIHDKSMILFPEPSAFALVFIPIFSFCLYYTRGG    180
ECO0061         LKRSVFYLFAFLCLIGIISILLQKTEIIHDKSMILFPEPSAFALVFIPIFSFCLYYTRGG    180
VREC1403        LKRSVFYLFAFLCLIGIISILLQKTEIIHDKSMILFPEPSAFALVFIPIFSFCLYYTRGG    180
NCTC13441       LKRSVFYLFAFLCLIGIISILLQKTEIIHDKSMILFPEPSAFALVFIPIFSFCLYYTRGG    180
ECO0218orf2     ------------------------------------------------------------    0
ECO0387orf2     ------------------------------------------------------------    0
ECO0218orf1     LKRSVFYLFAFLCLIGIISILLQKTEIIHDKSMILFPEPSAFALVFIPIFSFCLYYTRGG    180
ECO0387orf1     LKRSVFYLFAFLCLIGIISILLQKTEIIHDKSMILFPEPSAFALVFIPIFSFCLYYTRGG    180


ECO0353         GLLLLYILSLGIALGIQNLTMLVGIVISVFVMKKITIRQTIVILLGAWIFSMILSDLDIS    240
ECO0431         GLLLLYILSLGIALGIQNLTMLVGIVISVFVMKKITIRQTIVILLGAWIFSMILSDLDIS    240
ECO0411         GLLLLYILSLGIALGIQNLTMLVGIVISVFVMKKITIRQTIVILLGAWIFSMILSDLDIS    240
ECO0224         GLLLLYILSLGIALGIQNLTMLVGIVISVFVMKKITIRQTIVILLGAWIFSMILSDLDIS    240
ECO0433         GLLLLYILSLGIALGIQNLTMLVGIVISVFVMKKITIRQTIVILLGAWIFSMILSDLDIS    240
VREC0693        GLLLLYILSLGIALGIQNLTMLVGIVISVFVMKKITIRQTIVILLGAWIFSMILSDLDIS    240
VREC0829        GLLLLYILSLGIALGIQNLTMLVGIVISVFVMKKITIRQTIVILLGAWIFSMILSDLDIS    240
VREC0865        GLLLLYILSLGIALGIQNLTMLVGIVISVFVMKKITIRQTIVILLGAWIFSMILSDLDIS    240
VREC0926        GLLLLYILSLGIALGIQNLTMLVGIVISVFVMKKITIRQTIVILLGAWIFSMILSDLDIS    240
VREC1013        GLLLLYILSLGIALGIQNLTMLVGIVISVFVMKKITIRQTIVILLGAWIFSMILSDLDIS    240
VREC1073        GLLLLYILSLGIALGIQNLTMLVGIVISVFVMKKITIRQTIVILLGAWIFSMILSDLDIS    240
VREC1101        GLLLLYILSLGIALGIQNLTMLVGIVISVFVMKKITIRQTIVILLGAWIFSMILSDLDIS    240
VREC1106        GLLLLYILSLGIALGIQNLTMLVGIVISVFVMKKITIRQTIVILLGAWIFSMILSDLDIS    240
VREC1428        GLLLLYILSLGIALGIQNLTMLVGIVISVFVMKKITIRQTIVILLGAWIFSMILSDLDIS    240
VREC1630        GLLLLYILSLGIALGIQNLTMLVGIVISVFVMKKITIRQTIVILLGAWIFSMILSDLDIS    240
VRES0700        GLLLLYILSLGIALGIQNLTMLVGIVISVFVMKKITIRQTIVILLGAWIFSMILSDLDIS    240
VRES1160        GLLLLYILSLGIALGIQNLTMLVGIVISVFVMKKITIRQTIVILLGAWIFSMILSDLDIS    240
VREC0645        GLLLLYILSLGIALGIQNLTMLVGIVISVFVMKKITIRQTIVILLGAWIFSMILSDLDIS    240
VRES0739        GLLLLYILSLGIALGIQNLTMLVGIVISVFVMKKITIRQTIVILLGAWIFSMILSDLDIS    240
VRES1619        GLLLLYILSLGIALGIQNLTMLVGIVISVFVMKKITIRQTIVILLGAWIFSMILSDLDIS    240
VRES1610        GLLLLYILSLGIALGIQNLTMLVGIVISVFVMKKITIRQTIVILLGAWIFSMILSDLDIS    240
ECO0211         GLLLLYILSLGIALGIQNLTMLVGIVISVFVMKKITIRQTIVILLGAWIFSMILSDLDIS    240
ECO0061         GLLLLYILSLGIALGIQNLTMLVGIVISVFVMKKITIRQTIVILLGAWIFSMILSDLDIS    240
VREC1403        GLLLLYILSLGIALGIQNLTMLVGIVISVFVMKKITIRQTIVILLGAWIFSMILSDLDIS    240
NCTC13441       GLLLLYILSLGIALGIQNLTMLVGIVISVFVMKKITIRQTIVILLGAWIFSMILSDLDIS    240
ECO0218orf2     ---------------------------------------------------MILSDLDIS    9
ECO0387orf2     --------------------MLVGIVISVFVMKKITIRQTIVILLGAWIFSMILSDLDIS    40
ECO0218orf1     GATIALYIIFGYCVRYPEFNNVGRH-CD--------------------------------    207
ECO0387orf1     GYYCSIYYLWVLR-----------------------------------------------    193


ECO0353         YYTSRLDFKNTTNLSVLVYLSGIERAFLNFITSYGLGIGFQQMGVNGEIGIYQQILAELD    300
ECO0431         YYTSRLDFKNTTNLSVLVYLSGIERAFLNFITSYGLGIGFQQMGVNGEIGIYQQILAELD    300
ECO0411         YYTSRLDFKNTTNLSVLVYLSGIERAFLNFITSYGLGIGFQQMGVNGEIGIYQQILAELD    300
ECO0224         YYTSRLDFKNTTNLSVLVYLSGIERAFLNFITSYGLGIGFQQMGVNGEIGIYQQILAELD    300
ECO0433         YYTSRLDFKNTTNLSVLVYLSGIERAFLNFITSYGLGIGFQQMGVNGEIGIYQQILAELD    300
VREC0693        YYTSRLDFKNTTNLSVLVYLSGIERAFLNFITSYGLGIGFQQMGVNGEIGIYQQILAELD    300
VREC0829        YYTSRLDFKNTTNLSVLVYLSGIERAFLNFITSYGLGIGFQQMGVNGEIGIYQQILAELD    300
VREC0865        YYTSRLDFKNTTNLSVLVYLSGIERAFLNFITSYGLGIGFQQMGVNGEIGIYQQILAELD    300
VREC0926        YYTSRLDFKNTTNLSVLVYLSGIERAFLNFITSYGLGIGFQQMGVNGEIGIYQQILAELD    300
VREC1013        YYTSRLDFKNTTNLSVLVYLSGIERAFLNFITSYGLGIGFQQMGVNGEIGIYQQILAELD    300
VREC1073        YYTSRLDFKNTTNLSVLVYLSGIERAFLNFITSYGLGIGFQQMGVNGEIGIYQQILAELD    300
VREC1101        YYTSRLDFKNTTNLSVLVYLSGIERAFLNFITSYGLGIGFQQMGVNGEIGIYQQILAELD    300
VREC1106        YYTSRLDFKNTTNLSVLVYLSGIERAFLNFITSYGLGIGFQQMGVNGEIGIYQQILAELD    300
VREC1428        YYTSRLDFKNTTNLSVLVYLSGIERAFLNFITSYGLGIGFQQMGVNGEIGIYQQILAELD    300
VREC1630        YYTSRLDFKNTTNLSVLVYLSGIERAFLNFITSYGLGIGFQQMGVNGEIGIYQQILAELD    300
VRES0700        YYTSRLDFKNTTNLSVLVYLSGIERAFLNFITSYGLGIGFQQMGVNGEIGIYQQILAELD    300
VRES1160        YYTSRLDFKNTTNLSVLVYLSGIERAFLNFITSYGLGIGFQQMGVNGEIGIYQQILAELD    300
VREC0645        YYTSRLDFKNTTNLSVLVYLSGIERAFLNFITSYGLGIGFQQMGVNGEIGIYQQILAELD    300
VRES0739        YYTSRLDFKNTTNLSVLVYLSGIERAFLNFITSYGLGIGFQQMGVNGEIGIYQQILAELD    300
VRES1619        YYTSRLDFKNTTNLSVLVYLSGIERAFLNFITSYGLGIGFQQMGVNGEIGIYQQILAELD    300
VRES1610        YYTSRLDFKNTTNLSVLVYLSGIERAFLNFITSYGLGIGFQQMGVNGEIGIYQQILAELD    300
ECO0211         YYTSRLDFKNTTNLSVLVYLSGIERAFLNFITSYGLGIGFQQMGVNGEIGIYQQILAELD    300
ECO0061         YYTSRLDFKNTTNLSVLVYLSGIERAFLNFITSYGLGIGFQQMGVNGEIGIYQQILAELD    300
VREC1403        YYTSRLDFKNTTNLSVLVYLSGIERAFLNFITSYGLGIGFQQMGVNGEIGIYQQILAELD    300
NCTC13441       YYTSRLDFKNTTNLSVLVYLSGIERAFLNFITSYGLGIGFQQMGVNGEIGIYQQILAELD    300
ECO0218orf2     YYTSRLDFKNTTNLSVLVYLSGIERAFLNFITSYGLGIGFQQMGVNGEIGIYQQILAELD    69
```

```
ECO0387orf2      YYTSRLDFKNTTNLSVLVYLSGIERAFLNFITSYGLGIGFQQMGVNGEIGIYQQILAELD      100
ECO0218orf1      ------------------------------------------------------------      207
ECO0387orf1      ------------------------------------------------------------      193

ECO0353          APMLNIYDGSFISSKLISEFGVIGALMCIFYFFYFSRFYLRFKKSKRYSPQYILAYSFYM      360
ECO0431          APMLNIYDGSFISSKLISEFGVIGALMCIFYFFYFSRFYLRFKKSKRYSPQYILAYSFYM      360
ECO0411          APMLNIYDGSFISSKLISEFGVIGALMCIFYFFYFSRFYLRFKKSKRYSPQYILAYSFYM      360
ECO0224          APMLNIYDGSFISSKLISEFGVIGALMCIFYFFYFSRFYLRFKKSKRYSPQYILAYSFYM      360
ECO0433          APMLNIYDGSFISSKLISEFGVIGALMCIFYFFYFSRFYLRFKKSKRYSPQYILAYSFYM      360
VREC0693         APMLNIYDGSFISSKLISEFGVIGALMCIFYFFYFSRFYLRFKKSKRYSPQYILAYSFYM      360
VREC0829         APMLNIYDGSFISSKLISEFGVIGALMCIFYFFYFSRFYLRFKKSKRYSPQYILAYSFYM      360
VREC0865         APMLNIYDGSFISSKLISEFGVIGALMCIFYFFYFSRFYLRFKKSKRYSPQYILAYSFYM      360
VREC0926         APMLNIYDGSFISSKLISEFGVIGALMCIFYFFYFSRFYLRFKKSKRYSPQYILAYSFYM      360
VREC1013         APMLNIYDGSFISSKLISEFGVIGALMCIFYFFYFSRFYLRFKKSKRYSPQYILAYSFYM      360
VREC1073         APMLNIYDGSFISSKLISEFGVIGALMCIFYFFYFSRFYLRFKKSKRYSPQYILAYSFYM      360
VREC1101         APMLNIYDGSFISSKLISEFGVIGALMCIFYFFYFSRFYLRFKKSKRYSPQYILAYSFYM      360
VREC1106         APMLNIYDGSFISSKLISEFGVIGALMCIFYFFYFSRFYLRFKKSKRYSPQYILAYSFYM      360
VREC1428         APMLNIYDGSFISSKLISEFGVIGALMCIFYFFYFSRFYLRFKKSKRYSPQYILAYSFYM      360
VREC1630         APMLNIYDGSFISSKLISEFGVIGALMCIFYFFYFSRFYLRFKKSKRYSPQYILAYSFYM      360
VRES0700         APMLNIYDGSFISSKLISEFGVIGALMCIFYFFYFSRFYLRFKKSKRYSPQYILAYSFYM      360
VRES1160         APMLNIYDGSFISSKLISEFGVIGALMCIFYFFYFSRFYLRFKKSKRYSPQYILAYSFYM      360
VREC0645         APMLNIYDGSFISSKLISEFGVIGALMCIFYFFYFSRFYLRFKKSKRYSPQYILAYSFYM      360
VRES0739         APMLNIYDGSFISSKLISEFGVIGALMCIFYFFYFSRFYLRFKKSKRYSPQYILAYSFYM      360
VRES1619         APMLNIYDGSFISSKLISEFGVIGALMCIFYFFYFSRFYLRFKKSKRYSPQYILAYSFYM      360
VRES1610         APMLNIYDGSFISSKLISEFGVIGALMCIFYFFYFSRFYLRFKKSKRYSPQYILAYSFYM      360
ECO0211          APMLNIYDGSFISSKLISEFGVIGALMCIFYFFYFSRFYLRFKKSKRYSPQYILAYSFYM      360
ECO0061          APMLNIYDGSFISSKLISEFGVIGALMCIFYFFYFSRFYLRFKKSKRYSPQYILAYSFYM      360
VREC1403         APMLNIYDGSFISSKLISEFGVIGALMCIFYFFYFSRFYLRFKKSKRYSPQYILAYSFYM      360
NCTC13441        APMLNIYDGSFISSKLISEFGVIGALMCIFYFFYFSRFYLRFKKSKRYSPQYILAYSFYM      360
ECO0218orf2      APMLNIYDGSFISSKLISEFGVIGALMCIFYFFYFSRFYLRFKKSKRYSPQYILAYSFYM      129
ECO0387orf2      APMLNIYDGSFISSKLISEFGVIGALMCIFYFFYFSRFYLRFKKSKRYSPQYILAYSFYM      160
ECO0218orf1      ------------------------------------------------------------      207
ECO0387orf1      ------------------------------------------------------------      193

ECO0353          CFFIPLFIRGAGYINPYVFMLFSSIFLCKYHAKNILMKSNVQIAI     405
ECO0431          CFFIPLFIRGAGYINPYVFMLFSSIFLCKYHAKNILMKSNVQIAI     405
ECO0411          CFFIPLFIRGAGYINPYVFMLFSSIFLCKYHAKNILMKSNVQIAI     405
ECO0224          CFFIPLFIRGAGYINPYVFMLFSSIFLCKYHAKNILMKSNVQIAI     405
ECO0433          CFFIPLFIRGAGYINPYVFMLFSSIFLCKYHAKNILMKSNVQIAI     405
VREC0693         CFFIPLFIRGAGYINPYVFMLFSSIFLCKYHAKNILMKSNVQIAI     405
VREC0829         CFFIPLFIRGAGYINPYVFMLFSSIFLCKYHAKNILMKSNVQIAI     405
VREC0865         CFFIPLFIRGAGYINPYVFMLFSSIFLCKYHAKNILMKSNVQIAI     405
VREC0926         CFFIPLFIRGAGYINPYVFMLFSSIFLCKYHAKNILMKSNVQIAI     405
VREC1013         CFFIPLFIRGAGYINPYVFMLFSSIFLCKYHAKNILMKSNVQIAI     405
VREC1073         CFFIPLFIRGAGYINPYVFMLFSSIFLCKYHAKNILMKSNVQIAI     405
VREC1101         CFFIPLFIRGAGYINPYVFMLFSSIFLCKYHAKNILMKSNVQIAI     405
VREC1106         CFFIPLFIRGAGYINPYVFMLFSSIFLCKYHAKNILMKSNVQIAI     405
VREC1428         CFFIPLFIRGAGYINPYVFMLFSSIFLCKYHAKNILMKSNVQIAI     405
VREC1630         CFFIPLFIRGAGYINPYVFMLFSSIFLCKYHAKNILMKSNVQIAI     405
VRES0700         CFFIPLFIRGAGYINPYVFMLFSSIFLCKYHAKNILMKSNVQIAI     405
VRES1160         CFFIPLFIRGAGYINPYVFMLFSSIFLCKYHAKNILMKSNVQIAI     405
VREC0645         CFFIPLFIRGAGYINPYVFMLFSSIFLCKYHAKNILMKSNVQIAI     405
VRES0739         CFFIPLFIRGAGYINPYVFMLFSSIFLCKYHAKNILMKSNVQIAI     405
VRES1619         CFFIPLFIRGAGYINPYVFMLFSSIFLCKYHAKNILMKSNVQIAI     405
VRES1610         CFFIPLFIRGAGYINPYVFMLFSSIFLCKYHAKNILMKSNVQIAI     405
ECO0211          CFFIPLFIRGAGYINPYVFMLFSSIFLCKYHAKNILMKSNVQIAI     405
ECO0061          CFFIPLFIRGAGYINPYVFMLFSSIFLCKYHAKNILMKSNVQIAI     405
VREC1403         CFFIPLFIRGAGYINPYVFMLFSSIFLCKYHAKNILMKSNVQIAI     405
NCTC13441        CFFIPLFIRGAGYINPYVFMLFSSIFLCKYHAKNILMKSNVQIAI     405
ECO0218orf2      CFFIPLFIRGAGYINPYVFMLFSSIFLCKYHAKNILMKSNVQIAI     174
ECO0387orf2      CFFIPLFIRGAGYINPYVFMLFSSIFLCKYHAKNILMKSNVQIAI     205
ECO0218orf1      ---------------------------------------------     207
ECO0387orf1      ---------------------------------------------     193
```

CLUSTAL O(1.2.4) multiple sequence alignment **RfaZ**

```
VREC0645_1      MKNIRYIDKKDVENLIENKTSDDVIIFLSGPTSQKTPLSVLRTKDIIAVNGSVQYLFKS- 59
VRES1160        MKNIRYIDKKDVENLIENKTSDDVIIFLSGPTSQKTPLSVLRTKDIIAVNGSVQYLLSHN 60
VREC0645_2      -------------------------------------------------MVLCNICLSHN 11
VRES0708        MKNIRYIDKKDVENLIENKTSDDVIIFLSGPTSQKTPLSVLRTKDIIAVNGSVQYLLSHN 60
ECO0216         MKNIRYIDKKDVENLIENKTSDDVIIFLSGPTSQKTPLSVLRTKDIIAVNGSVQYLLSHN 60
ECO0172         MKNIRYIDKKDVENLIENKTSDDVIIFLSGPTSQKTPLSVLRTKDIIAVNGSVQYLLSHN 60
ECO0056         MKNIRYIDKKDVENLIENKTSDDVIIFLSGPTSQKTPLSVLRTKDIIAVNGSVQYLLSHN 60
NCTC13441       MKNIRYIDKKDVENLIENKTSDDVIIFLSGPTSQKTPLSVLRTKDIIAVNGSVQYLLSHN 60
VRES1100        MKNIRYIDKKDVENLIENKTSDDVIIFLSGPTSQKTPLSVLRTKDIIAVNGSVQYLLSHN 60
VRES0739        MKNIRYIDKKDVENLIENKTSDDVIIFLSGPTSQKTPLSVLRTKDIIAVNGSVQYLLSHN 60
VRES0710        MKNIRYIDKKDVENLIENKTSDDVIIFLSGPTSQKTPLSVLRTKDIIAVNGSVQYLLSHN 60
ECO0257         MKNIRYIDKKDVENLIENKTSDDVIIFLSGPTSQKTPLSVLRTKDIIAVNGSVQYLLSHN 60
VRES1619        MKNIRYIDKKDVENLIENKTSDDVIIFLSGPTSQKTPLSVLRTKDIIAVNGSVQYLLSHN 60
VRES1610        MKNIRYIDKKDVENLIENKTSDDVIIFLSGPTSQKTPLSVLRTKDIIAVNGSVQYLLSHN 60
                                                                     :  :.

VREC0645_1      ------------------------------------------------------------ 59
VRES1160        IVPFIYVLTDVRFLHQRRDDFYKFSQRSRYTIVNVDVYEHASKEDKLYILQNCLVLRSFY 120
VREC0645_2      IVPFIYVLTDVRFLHQRRDDFYKFSQRSRYTIVNVDVYEHASKEDKLYILQNCLVLRSFY 71
VRES0708        IVPFIYVLTDVRFLHQRRDDFYKFSQRSRYTIVNVDVYEHASKEDKLYILQNCLVLRSFY 120
ECO0216         IVPFIYVLTDVRFLHQRRDDFYKFSQRSRYTIVNVDVYEHASKEDKLYILQNCLVLRSFY 120
ECO0172         IVPFIYVLTDVRFLHQRRDDFYKFSQRSRYTIVNVDVYEHASKEDKLYILQNCLVLRSFY 120
ECO0056         IVPFIYVLTDVRFLHQRRDDFYKFSQRSRYTIVNVDVYEHASKEDKLYILQNCLVLRSFY 120
NCTC13441       IVPFIYVLTDVRFLHQRRDDFYKFSQRSRYTIVNVDVYEHASKEDKLYILQNCLVLRSFY 120
VRES1100        IVPFIYVLTDVRFLHQRRDDFYKFSQRSRYTIVNVDVYEHASKEDKLYILQNCLVLRSFY 120
VRES0739        IVPFIYVLTDVRFLHQRRDDFYKFSQRSRYTIVNVDVYEHASKEDKLYILQNCLVLRSFY 120
VRES0710        IVPFIYVLTDVRFLHQRRDDFYKFSQRSRYTIVNVDVYEHASKEDKLYILQNCLVLRSFY 120
ECO0257         IVPFIYVLTDVRFLHQRRDDFYKFSQRSRYTIVNVDVYEHASKEDKLYILQNCLVLRSFY 120
VRES1619        IVPFIYVLTDVRFLHQRRDDFYKFSQRSRYTIVNVDVYEHASKEDKLYILQNCLVLRSFY 120
VRES1610        IVPFIYVLTDVRFLHQRRDDFYKFSQRSRYTIVNVDVYEHASKEDKLYILQNCLVLRSFY 120

VREC0645_1      ------------------------------------------------------------ 59
VRES1160        RREKGGFIKKIKFNILSQIHKELLISVPLSKKGRLVGFCKDISFGYCSCHTIAFAAIQIA 180
VREC0645_2      RREKGGFIKKIKFNILSQIHKELLISVPLSKKGRLVGFCKDISFGYCSCHTIAFAAIQIA 131
VRES0708        RREKGGFIKKIKFNILSQIHKELLISVPLSKKGRLVGFCKDISFGYCSCHTIAFAAIQIA 180
ECO0216         RREKGGFIKKIKFNILSQIHKELLISVPLSKKGRLVGFCKDISFGYCSCHTIAFAAIQIA 180
ECO0172         RREKGGFIKKIKFNILSQIHKELLISVPLSKKGRLVGFCKDISFGYCSCHTIAFAAIQIA 180
ECO0056         RREKGGFIKKIKFNILSQIHKELLISVPLSKKGRLVGFCKDISFGYCSCHTIAFAAIQIA 180
NCTC13441       RREKGGFIKKIKFNILSQIHKELLISVPLSKKGRLVGFCKDISFGYCSCHTIAFAAIQIA 180
VRES1100        RREKGGFIKKIKFNILSQIHKELLISVPLSKKGRLVGFCKDISFGYCSCHTIAFAAIQIA 180
VRES0739        RREKGGFIKKIKFNILSQIHKELLISVPLSKKGRLVGFCKDISFGYCSCHTIAFAAIQIA 180
VRES0710        RREKGGFIKKIKFNILSQIHKELLISVPLSKKGRLVGFCKDISFGYCSCHTIAFAAIQIA 180
ECO0257         RREKGGFIKKIKFNILSQIHKELLISVPLSKKGRLVGFCKDISFGYCSCHTIAFAAIQIA 180
VRES1619        RREKGGFIKKIKFNILSQIHKELLISVPLSKKGRLVGFCKDISFGYCSCHTIAFAAIQIA 180
VRES1610        RREKGGFIKKIKFNILSQIHKELLISVPLSKKGRLVGFCKDISFGYCSCHTIAFAAIQIA 180

VREC0645_1      ------------------------------------------------------------ 59
VRES1160        YSLKYARIICSGLDLTGNCSRFYDENNNPMPSELSKDLFKILPFFRFMRDNVEDINIYNL 240
VREC0645_2      YSLKYARIICSGLDLTGNCSRFYDENNNPMPSELSKDLFKILPFFRFMRDNVEDINIYNL 191
VRES0708        YSLKYARIICSGLDLTGNCSRFYDENNNPMPSELSKDLFKILSFFRFMRDNVEDINIYNL 240
ECO0216         YSLKYARIICSGLDLTGNCSRFYDENNNPMPSELSKDLFKILPFFRFMRDNVEDINIYNL 240
ECO0172         YSLKYARIICSGLDLTGNCSRFYDENNNPMPSELSKDLFKILPFFRFMRDNVEDINIYNL 240
ECO0056         YSLKYARIICSGLDLTGNCSRFYDENNNPMPSELSKDLFKILPFFRFMRDNVEDINIYNL 240
NCTC13441       YSLKYARIICSGLDLTGNCSRFYDENNNPMPSELSKDLFKILPFFRFMRDNVEDINIYNL 240
VRES1100        YSLKYARIICSGLDLTGNCSRFYDENNNPMPSELSKDLFKILPFFRFMRDNVEDINIYNL 240
VRES0739        YSLKYARIICSGLDLTGNCSRFYDENNNPMPSELSKDLFKILPFFRFMRDNVEDINIYNL 240
VRES0710        YSLKYARIICSGLDLTGNCSRFYDENNNPMPSELSKDLFKILPFFRFMRDNVEDINIYNL 240
ECO0257         YSLKYARIICSGLDLTGNCSRFYDENNNPMPSELSKDLFKILPFFRFMRDNVEDINIYNL 240
VRES1619        YSLKYARIICSGLDLTGNCSRFYDENNNPMPSKLSKDLFKILPFFRFMRDNVEDINIYNL 240
VRES1610        YSLKYARIICSGLDLTGNCSRFYDENNNPMPSKLSKDLFKILPFFRFMRDNVEDINIYNL 240

VREC0645_1      ------------------------------------------------------------ 59
VRES1160        SDDTAISYDVIPFIKFQDISREESKDKTRKKCNIELQLILMLINHPETKVIWYKNAFRNF 300
VREC0645_2      SDDTAISYDVIPFIKFQDISREESKDKTRKKMQYRTSTDSYA------------N----- 234
VRES0708        SDDTAISYDVIPFIKFQDISREESKDKTRKKMQYRTSTDSYA------------N----- 283
ECO0216         SDDTAISYDVIPFIKFQDISREESKDKTRKKMQYRTSTDSYA------------N----- 283
ECO0172         SDDTAISYDVIPFIKFQDISREESKDKTRKKMQYRTSTDSYA------------N----- 283
ECO0056         SDDTAISYDVIPFIKFQDISREESKDKTRKKMQYRTSTDSYA------------N----- 283
NCTC13441       SDDTAISYDVIPFIKFQDISREESKDKTRKKMQYRTSTDSYA------------N----- 283
VRES1100        SDDTAISYDVIPFIKFQDISREESKDKTRKKMQYRTSTDSYA------------N----- 283
VRES0739        SDDTAISYDVIPFIKFQDISREESKDKTRKKMQYRTSTDSYA------------N----- 283
VRES0710        SDDTAISYDVIPFIKFQDISREESKDKTRKKMQYRTSTDSYA------------N----- 283
```

```
ECO0257         SDDTAISYDVIPFIKFQDISREESKDKTRKKMQYRTSTDSYA------------N-----283
VRES1619        SDDTAISYDVIPFIKFQDISREESKDKTRKKMQYRTSTDSYA------------N-----283
VRES1610        SDDTAISYDVIPFIKFQDISREESKDKTRKKMQYRTSTDSYA------------N-----283

VREC0645_1      -------------         59
VRES1160        SQKEKILYKQNKN         313
VREC0645_2      -------------         234
VRES0708        -------------         283
ECO0216         -------------         283
ECO0172         -------------         283
ECO0056         -------------         283
NCTC13441       -------------         283
VRES1100        -------------         283
VRES0739        -------------         283
VRES0710        -------------         283
ECO0257         -------------         283
VRES1619        -------------         283
VRES1610        -------------         283
```

CLUSTAL O(1.2.4) multiple sequence alignment **RfaJ**

```
ECO0216         MDSFPAIEIDKVKAWDFRLANINTSECLNVAYGVDANYLDGVGVSITSIVLNNRHINLDF  60
ECO0172         MDSFPAIEIDKVKAWDFRLANINTSECLNVAYGVDANYLDGVGVSITSIVLNNRHINLDF  60
ECO0056         MDSFPAIEIDKVKAWDFRLANINTSECLNVAYGVDANYLDGVGVSITSIVLNNRHINLDF  60
NCTC13441       MDSFPAIEIDKVKAWDFRLANINTSECLNVAYGVDANYLDGVGVSITSIVLNNRHINLDF  60
VRES1619        MDSFPAIEIDKVKAWDFRLANINTSECLNVAYGVDANYLDGVGVSITSIVLNNRHINLDF  60
VRES1610        MDSFPAIEIDKVKAWDFRLANINTSECLNVAYGVDANYLDGVGVSITSIVLNNRHINLDF  60
VRES1160        MDSFPAIEIDKVKAWDFRLANINTSECLNVAYGVDANYLDGVGVSITSIVLNNRHINLDF  60
VRES1100        MDSFPAIEIDKVKAWDFRLANINTSECLNVAYGVDANYLDGVGVSITSIVLNNRHINLDF  60
VRES0739        MDSFPAIEIDKVKAWDFRLANINTSECLNVAYGVDANYLDGVGVSITSIVLNNRHINLDF  60
VRES0710        MDSFPAIEIDKVKAWDFRLANINTSECLNVAYGVDANYLDGVGVSITSIVLNNRHINLDF  60
VRES0708        MDSFPAIEIDKVKAWDFRLANINTSECLNVAYGVDANYLDGVGVSITSIVLNNRHINLDF  60
VREC0645        MDSFPAIEIDKVKAWDFRLANINTSECLNVAYGVDANYLDGVGVSITSIVLNNRHINLDF  60
ECO0257         MDSFPAIEIDKVKAWDFRLANINTSECLNVAYGVDANYLDGVGVSITSIVLNNRHINLDF  60
                ************************************************************

ECO0216         YIIADVYNDAFFQKVAKLAEQYQLRITLYRINTDKLQCLPCTQVWSRAMYFRLFAFQLLG  120
ECO0172         YIIADVYNDAFFQKVAKLAEQYQLRITLYRINTDKLQCLPCTQVWSRAMYFRLFAFQLLG  120
ECO0056         YIIADVYNDAFFQKVAKLAEQYQLRITLYRINTDKLQCLPCTQVWSRAMYFRLFAFQLLG  120
NCTC13441       YIIADVYNDAFFQKVAKLAEQYQLRITLYRINTDKLQCLPCTQVWSRAMYFRLFAFQLLG  120
VRES1619        YIIADVYNDAFFQKVAKLAEQYQLRITLYRINTDKLQCLPCTQVWSRAMYFRLFAFQLLG  120
VRES1610        YIIADVYNDAFFQKVAKLAEQYQLRITLYRINTDKLQCLPCTQVWSRAMYFRLFAFQLLG  120
VRES1160        YIIADVYNDAFFQKVAKLAEQYQLRITLYRINTDKLQCLPCTQVWSRAMYFRLFAFQLLG  120
VRES1100        YIIADVYNDAFFQKVAKLAEQYQLRITLYRINTDKLQCLPCTQVWSRAMYFRLFAFQLLG  120
VRES0739        YIIADVYNDAFFQKVAKLAEQYQLRITLYRINTDKLQCLPCTQVWSRAMYFRLFAFQLLG  120
VRES0710        YIIADVYNDAFFQKVAKLAEQYQLRITLYRINTDKLQCLPCTQVWSRAMYFRLFAFQLLG  120
VRES0708        YIIADVYNDAFFQKVAKLAEQYQLRITLYRINTDKLQCLPCTQVWSRAMYFRLFAFQLLG  120
VREC0645        YIIADVYNDAFFQKVAKLAEQYQLRITLYRINTDKLQCLPCTQVWSRAMYFRLFAFQLLG  120
ECO0257         YIIADVYNDAFFQKVAKLAEQYQLRITLYRINTDKLQCLPCTQVWSRAMYFRLFAFQLLG  120
                ************************************************************

ECO0216         LTLDRLLYLDADVVCKGDISQLLHLDLNGAVAAVVKDVDPMQEKAASRLSDPELLGQYFN  180
ECO0172         LTLDRLLYLDADVVCKGDISQLLHLDLNGAVAAVVKDVDPMQEKAASRLSDPELLGQYFN  180
ECO0056         LTLDRLLYLDADVVCKGDISQLLHLDLNGAVAAVVKDVDPMQEKAASRLSDPELLGQYFN  180
NCTC13441       LTLDRLLYLDADVVCKGDISQLLHLDLNGAVAAVVKDVDPMQEKAASRLSDPELLGQYFN  180
VRES1619        LTLDRLLYLDADVVCKGDISQLLHLDLNGAVAAVVKDVDPMQEKAASRLSDPELLGQYFN  180
VRES1610        LTLDRLLYLDADVVCKGDISQLLHLDLNGAVAAVVKDVDPMQEKAASRLSDPELLGQYFN  180
VRES1160        LTLDRLLYLDADVVCKGDISQLLHLDLNGAVAAVVKDVDPMQEKAASRLSDPELLGQYFN  180
VRES1100        LTLDRLLYLDADVVCKGDISQLLHLDLNGAVAAVVKDVDPMQEKAASRLSDPELLGQYFN  180
VRES0739        LTLDRLLYLDADVVCKGDISQLLHLDLNGAVAAVVKDVDPMQEKAASRLSDPELLGQYFN  180
VRES0710        LTLDRLLYLDADVVCKGDISQLLHLDLNGAVAAVVKDVDPMQEKAASRLSDPELLGQYFN  180
VRES0708        LTLDRLLYLDADVVCKGDISQLLHLDLNGAVAAVVKDVDPMQEKAASRLSDPELLGQYFN  180
VREC0645        LTLDRLLYLDADVVCKGDISQLLHLDLNGAVAAVVKDVDPMQEKAASRLSDPELLGQYFN  180
ECO0257         LTLDRLLYLDADVVCKGDISQLLHLDLNGAVAAVVKDVDPMQEKAASRLSDPELLGQYFN  180
                ************************************************************

ECO0216         SCVVYLDLKKWANAKLTEKALSILMSKDNVYKYPDQDVMNVLLKGMTIFLPRGYNTIYTI  240
ECO0172         SGVVYLDLKKWANAKLTEKALSILMSKDNVYKYPDQDVMNVLLKGMTIFLPRGYNTIYTI  240
ECO0056         SGVVYLDLKKWANAKLTEKALSILMSKDNVYKYPDQDVMNVLLKGMTIFLPRGYNTIYTI  240
NCTC13441       SGVVYLDLKKWANAKLTEKALSILMSKDNVYKYPDQDVMNVLLKGMTIFLPRGYNTIYTI  240
VRES1619        SGVVYLDLKKWANAKLTEKALSILMSKDNVYKYPDQDVMNVLLKGMTIFLPRGYNTIYTI  240
VRES1610        SGVVYLDLKKWANAKLTEKALSILMSKDNVYKYPDQDVMNVLLKGMTIFLPRGYNTIYTI  240
VRES1160        SGVVYLDLKKWANAKLTEKALSILMSKDNVYKYPDQDVMNVLLKGMTIFLPRGYNTIYTI  240
VRES1100        SGVVYLDLKKWANAKLTEKALSILMSKDNVYKYPDQDVMNVLLKGMTIFLPRGYNTIYTI  240
VRES0739        SGVVYLDLKKWANAKLTEKALSILMSKDNVYKYPDQDVMNVLLKGMTIFLPRGYNTIYTI  240
VRES0710        SGVVYLDLKKWANAKLTEKALSILMSKDNVYKYPDQDVMNVLLKGMTIFLPRGYNTIYTI  240
VRES0708        SGVVYLDLKKWANAKLTEKALSILMSKDNVYKYPDQDVMNVLLKGMTIFLPRGYNTIYTI  240
VREC0645        SGVVYLDLKKWANAKLTEKALSILMSKDNVYKYPDQDVMNVLLKGMTIFLPRGYNTIYTI  240
ECO0257         SGVVYLDLKKWANAKLTEKALSILMSKDNVYKYPDQDVMNVLLKGMTIFLPRGYNTIYTI  240
                * **********************************************************

ECO0216         KSELKDKTHQNYKKLIIENTLLIHYTGATKPWHKWAIYPSTKYYKIALDNSPWKDDSPRD  300
ECO0172         KSELKDKTHQNYKKLIIENTLLIHYTGATKPWHKWAIYPSTKYYKIALDNSPWKDDSPRD  300
ECO0056         KSELKDKTHQNYKKLIIENTLLIHYTGATKPWHKWAIYPSTKYYKIALDNSPWKDDSPRD  300
NCTC13441       KSELKDKTHQNYKKLIIENTLLIHYTGATKPWHKWAIYPSTKYYKIALDNSPWKDDSPRD  300
VRES1619        KSELKDKTHQNYKKLIIENTLLIHYTGATKPWHKWAIYPSTKYYKIALDNSPWKDDSPRD  300
VRES1610        KSELKDKTHQNYKKLIIENTLLIHYTGATKPWHKWAIYPSTKYYKIALDNSPWKDDSPRD  300
VRES1160        KSELKDKTHQNYKKLIIENTLLIHYTGATKPWHKWAIYPSTKYYKIALDNSPWKDDSPRD  300
VRES1100        KSELKDKTHQNYKKLIIENTLLIHYTGATKPWHKWAIYPSTKYYKIALDNSPWKDDSPRD  300
VRES0739        KSELKDKTHQNYKKLIIENTLLIHYTGATKPWHKWAIYPSTKYYKIALDNSPWKDDSPRD  300
VRES0710        KSELKDKTHQNYKKLIIENTLLIHYTGATKPWHKWAIYPSTKYYKIALDNSPWKDDSPRD  300
VRES0708        KSELKDKTHQNYKKLIIENTLLIHYTGATKPWHKWAIYPSTKYYKIALDNSPWKDDSPRD  300
VREC0645        KSELKDKTHQNYKKLIIENTLLIHYTGATKPWHKWAIYPSTKYYKIALDNSPWKDDSPRD  300
ECO0257         KSELKDKTHQNYKKLIIENTLLIHYTGATKPWHKWAIYPSTKYYKIALDNSPWKDDSPRD  300
                ************************************************************
```

```
ECO0216        AKSIIEFKKRYKHLLVQHHYISGIIAGVCYLCRKYYPK    338
ECO0172        AKSIIEFKKRYKHLLVQHHYISGIIAGVCYLCRKYYPK    338
ECO0056        AKSIIEFKKRYKHLLVQHHYISGIIAGVCYLCRKYYPK    338
NCTC13441      AKSIIEFKKRYKHLLVQHHYISGIIAGVCYLCRKYYPK    338
VRES1619       AKSIIEFKKRYKHLLVQHHYISGIIAGVCYLCRKYYPK    338
VRES1610       AKSIIEFKKRYKHLLVQHHYISGIIAGVCYLCRKYYPK    338
VRES1160       AKSIIEFKKRYKHLLVQHHYISGIIAGVCYLCRKYYPK    338
VRES1100       AKSIIEFKKRYKHLLVQHHYISGIIAGVCYLCRKYYPK    338
VRES0739       AKSIIEFKKRYKHLLVQHHYISGIIAGVCYLCRKYYPK    338
VRES0710       AKSIIEFKKRYKHLLVQHHYISGIIAGVCYLCRKYYPK    338
VRES0708       AKSIIEFKKRYKHLLVQHHYISGIIAGVCYLCRKYYPK    338
VREC0645       AKSIIEFKKRYKHLLVQHHYISGIIAGVCYLCRKYYPK    338
ECO0257        AKSIIEFKKRYKHLLVQHHYISGIIAGVCYLCRKYYPK    338
               **************************************
```
