## Supplementary Fig. S6 for "A novel therapeutic antibody screening method using bacterial high-content imaging reveals functional antibody binding phenotypes of *Escherichia coli* ST131"

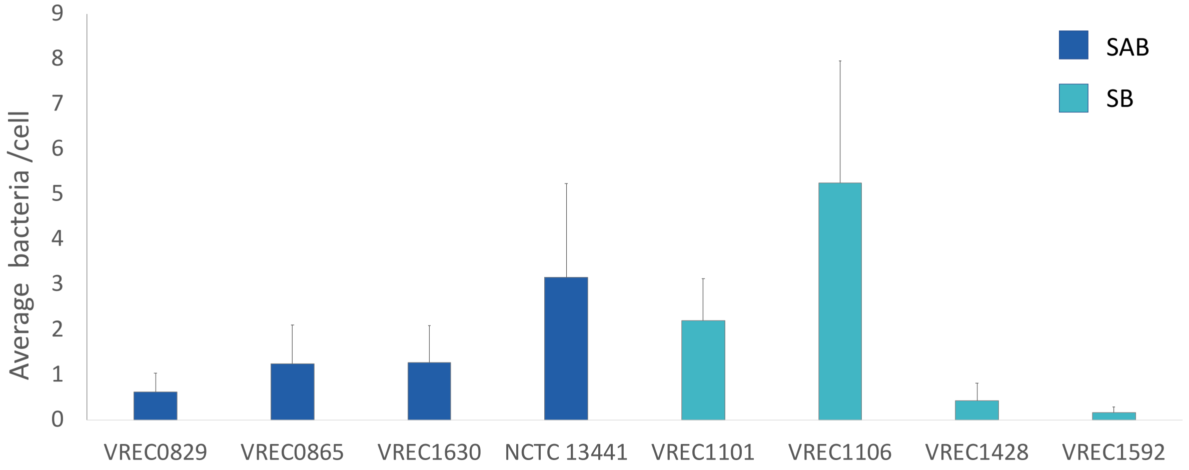


**Supplementary Figure S6: Background differences in phagocytosis for isolates without antibody present.** Macrophages were stained with CellMasks Orange (cytoplasm) and DAPI (nucleic acid) and bacteria were visualised using KM467 labelled with Alexa Flour 647. Images were obtained using a 40x air objective on the Opera Phenix. Images were analysed in Harmony using predefined building blocks to segment nuclei and cytoplasm to define the number of cells, and to count individual bacteria (using Find Spots) within the cells. Error bars represent standard deviation of 3 replicates.
