## Supplementary Fig. S7 for "A novel therapeutic antibody screening method using bacterial high-content imaging reveals functional antibody binding phenotypes of *Escherichia coli* ST131"

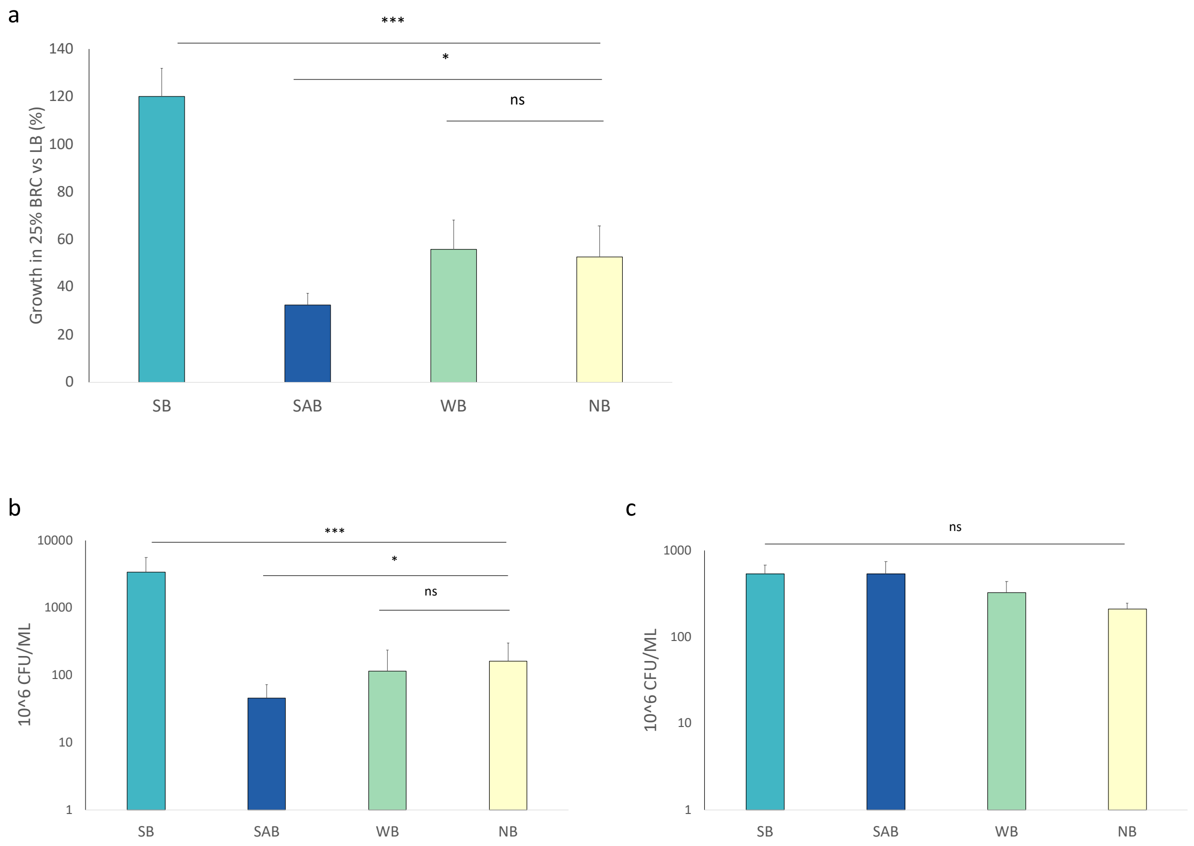


**Supplementary Figure S7: Serum Resistance Reproducibility.** Growth in LB broth in the presence or absence of 25% BRC was measured by OD600 after 4 hours and plotted as percentage growth in the presence of serum compared to LB alone (**a**). Growth in LB broth in the presence or absence of 25% BRC was also measured by cfu counting after 4 hours (**b)** and **(c**). The average of representative isolates per phenotype (SB=4, SAB=4, WB=3, NB =3) of 3 replicates is shown, and error bars represent standard deviation. Significance was determined by *t*-test (*=<0.05, **= between 0.001 and 0.05, *** <0.001).
