## Supplementary Table S2 for "A novel therapeutic antibody screening method using bacterial high-content imaging reveals functional antibody binding phenotypes of *Escherichia coli* ST131"

### Supplementary Table 2: Antibody binding analysis pipeline (Harmony v4.8)

| | | | |
|---|---|---|---|
| **Input Image** | Input | | |
| | **Flatfield Correction**: Basic Brightfield Correction<br>**Stack Processing**: Individual Planes<br>Create Global Image<br>**Min. Global Binning**: Dynamic | | |
| **Find Spots** | Input | Method | Output |
| | **Channel**: DAPI<br>**ROI**: None | **Method**: D<br>Detection Sensitivity: 0.5<br>Splitting Sensitivity: 0.5<br>Background correction: 0.5<br>Calculate Spot Properties | Output Population: Spots |
| **Calculate Morphology Properties** | Input | Method | Output |
| | **Population**: Spots<br>**Region**: Spot | **Method**: Standard<br>Area<br>Length | Property Prefix: Spot morphology |
| **Select Population** | Input | Method | Output |
| | **Population**: Spots | **Method**: Filter by Property<br>Spot morphology Length[µm]: <5<br>Spot morphology area [$\mu m^2$]: >0.7<br>Boolean Operations: F1 and F2 | Output Population: Spots Selected |
| **Select Population (2)** | Input | Method | Output |
| | **Population**: Spots Selected | Method: Common Filters<br>Remove Border Objects<br>Region: Spot | Output Population: Spots Selected no border |
| **Select Region** | Input | Method | Output |
| | **Population**: Spots Selected no border<br>**Region**: Spot | **Method**: Resize Region [µm/px]<br>Outer Border: -0.5µm<br>Inner Border: INF µm | Output Population: final bacteria |
| **Calculate Intensity Properties** | Input | Method | Output |
| | **Channel**: Alexa 647<br>**Population**: Spots Selected no border<br>**Region**: final bacteria | **Method**: Standard<br>Mean<br>Standard Deviation<br>Coefficient of Variance<br>Median<br>Maximum<br>Minimum<br>Contrast | Property Prefix: Intensity final bacteria Alexa 647 |
| **Select Population (3)** | Input | Method | Output |
| | **Population**: Spots Selected no border | **Method**: Filter by Property<br>Intensity final bacteria Alexa 647 Mean: > 600 | Output Population: Strong ALEXA |
| **Select Population (4)** | Input | Method | Output |

| | | | |
|---|---|---|---|
| | **Population**: Spots Selected no border | **Method**: Filter by Property<br>Intensity final bacteria Alexa 647 Mean: < 599<br>Intensity final bacteria Alexa 647 Mean: > 180 | Output Population: Weak ALEXA |
| **Select Population (5)** | Input | Method | Output |
| | **Population**: Spots Selected no border | **Method**: Filter by Property<br>Intensity final bacteria Alexa 647 Mean: > 180 | Output Population: Total ALEXA |
| **Calculate Position Properties** | Input | Method | Output |
| | **Population**: Spots Selected no border<br>**Region**: final bacteria | **Method**: Standard<br>Nearest Neighbour Distance | Property Prefix: final bacteria nearest neighbour distance |
| **Select Population (6)** | Input | Method | Output |
| | **Population**: Spots Selected no border | **Method**: Filter by Property<br>: Filter by Property<br>Final bacteria nearest neighbour distance Nearest Neighbour Distance [µm] < 1 | Output Population: Neighbouring bacteria |
| **Modify Population** | Input | Method | Output |
| | **Population**: Neighbouring bacteria<br>**Region**: final bacteria | **Method**: Cluster by Distance<br>Distance: 0.2 µm<br>Area: > 2 $\mu m^2$ | Output Population: Clustered bacteria by distance<br>Output Region: Cluster of bacteria by distance |
| **Calculate Properties** | Input | Method | Output |
| | **Population**: Clustered bacteria by distance | **Method**: By Related Population<br>Related Population: Neighbouring bacteria<br>Number of Neighbouring bacteria | Property Suffix: bacteria in clusters |
| **Select Population (7)** | Input | Method | Output |
| | **Population**: Clustered bacteria by distance | **Method**: Filter by Property<br>Number of Neighbouring bacteria – bacteria in clusters: > 4 | Output Population: Cluster >4 bacteria |
| **Calculate Properties (2)** | Input | Method | Output |
| | **Population**: Cluster >4 bacteria | **Method**: By Related Population<br>Related Population: Neighbouring bacteria<br>Number of Neighbouring bacteria | Property Suffix: bacteria per real cluster |
| **Calculate Morphology Properties (2)** | Input | Method | Output |
| | **Population**: Clustered bacteria by distance<br>**Region**: Cluster of bacteria by distance | **Method**: Standard<br>Area | Property Prefix: area cluster of bacteria |
| **Select Population (8)** | Input | Method | Output |
| | **Population**: Clustered bacteria by distance | Method: Filter by Property<br>area cluster of bacteria Area [µm2]: >13 | Output Population: Agglutinated bacteria |
| **Calculate Properties (3)** | Input | Method | Output |
| | **Population**: Agglutinated by area | **Method**: By Related Population<br>Related Population: Neighbouring bacteria<br>Number of Neighbouring bacteria | Property Suffix: number of agglutinated bacteria |

### Well Results: list of outputs

**Population: Spots Selected no border**
Number of Objects
Relative Spot Intensity: Mean per Well
Corrected Spot Intensity: Mean per Well
Uncorrected Spot Intensity: Mean per Well
Spot Contrast: Mean per Well
Spot Background Intensity: Mean per Well
Spot Area [px$^2$]: Mean per Well
Region Intensity: Mean per Well
Spot to Region Intensity: Mean per Well
Spot morphology Area [μm2]: Mean per Well
Spot morphology Length [μm]: Mean per Well
Intensity final bacteria Alexa 647 Mean: Mean per Well
Intensity final bacteria Alexa 647 StdDev: Mean per Well
Intensity final bacteria Alexa 647 Median: Mean per Well
Intensity final bacteria Alexa 647 Maximum: Mean per Well
Intensity final bacteria Alexa 647 Minimum: Mean per Well
Intensity final bacteria Alexa 647 Su: Mean per Well
Intensity final bacteria Alexa 647 CV [%]: Mean per Well
Intensity final bacteria Alexa 647 Contrast: Mean per Well
Strong ALEXA: Mean per Well
Weak ALEXA: Mean per Well
Total ALEXA: Mean per Well
final bacteria nearest neighbour distance Nearest Neighbour Distance [μm]: Mean per Well
Neighbouring bacteria: Mean per Well

**Population: Strong ALEXA**
Number of Objects

**Population: Weak ALEXA**
Number of Objects

**Population: Total ALEXA**
Number of Objects

**Population: Clustered bacteria by distance**
Number of Neighbouring bacteria – bacteria in clusters: Mean per Well
Cluster >4 bacteria: Mean per Well
area cluster of bacteria Area [μm2]: Mean per Well
Agglutinated by area: mean and standard deviation

**Population: Cluster >4 bacteria**
Number of Neighbouring bacteria – bacteria in clusters: Mean per Well
Number of Neighbouring bacteria – bacteria per real cluster: Mean per Well

**Population: Agglutinated by area**
Number of Neighbouring bacteria – bacteria in clusters: Mean per Well
Cluster >4 bacteria: Mean per Well
area cluster of bacteria Area [μm2]: Mean per Well
Number of Neighbouring bacteria – number of agglutinating bacteria: Mean per Well

### Formula Outputs:

Formula: 100*(a/b)
Population a: Cluster >4 bacteria – Number of Neighbouring bacteria – bacteria per real cluster Sum
Population b: Spots Selected no border – Number of Objects
**% clustering**

Formula: 100*(a/b)
Population a: Agglutinated by area – Number of Neighbouring bacteria – number of agglutinated bacteria Sum
Population b: Spots Selected no border – Number of Objects
**% agglutination**

Formula: 100*(a/b)
Population a: Total ALEXA – Number of Objects
Population b: Spots Selected no border – Number of Objects
**% total binding**

Formula: 100*(a/b)
Population a: Strong ALEXA – Number of Objects
Population b: Spots Selected no border – Number of Objects
**% strong binding**

Formula: 100*(a/b)
Population a: Weak ALEXA – Number of Objects
Population b: Spots Selected no border – Number of Objects
**% weak binding**
