## Supplementary Table S5 for "A novel therapeutic antibody screening method using bacterial high-content imaging reveals functional antibody binding phenotypes of *Escherichia coli* ST131"

### Supplementary Table 5: Macrophage Phagocytosis Pipeline (Harmony v4.8)

| | | | |
|---|---|---|---|
| **Input Image** | Input | | |
| | **Flatfield Correction**: None, Brightfield Correction<br>**Stack Processing**: Maximum Projection<br>**Min. Global Binning**: Dynamic | | |
| **Find Nuclei** | Input | Method | Output |
| | **Channel**: DAPI<br>**ROI**: None | **Method**: B<br>Common Threshold: 0.4<br>Area: > 30um$^2$<br>Splitting Coefficient: 7<br>Individual Threshold: 0.4<br>Contrast: > 0.1 | Output Population: CELL |
| **Find Cytoplasm** | Input | Method | Output |
| | **Channel**: MitoTracker Orange<br>**Nuclei**: CELL | **Method**: D<br>Individual Threshold: 0.15 | |
| **Select Cell Region** | Input | Method | Output |
| | **Population**: CELL | **Method**: Resize Region (%)<br>Region Type: Cell Region<br>Outer Border: 1%<br>Inner Border: 100% | Output Region: Cell Region |
| **Find Spots** | Input | Method | Output |
| | **Channel**: Alexa 647<br>**ROI**: CELL<br>**ROI Region**: Cell | **Method**: B<br>Detection Sensitivity: 0.5<br>Splitting Sensitivity: 0.5<br>Calculate Spot Properties | Output Population: intracellular bacteria |

#### Results:

**Method**: List of outputs
Population: CELL
Number of objects

**Method**: Standard Output
CELL – Number of Objects: Object Count
Output Name: CELL – Number of Objects

**Method**: Standard Output
Intracellular bacteria – Number of Objects: Object Count
Output Name: intracellular bacteria – Number of Objects

**Method**: Formula Output
Formula: a/b
Population Type: Objects
Variable a: intracellular bacteria – Number of Objects
Variable b: CELL – Number of Objects
**Output: Number of intracellular bacteria per cell**
